# A Method to Analyze Low-Quality Archaic Human Genomes and its Application to the Teshik-Tash 1 Neandertal

**DOI:** 10.64898/2026.08.10.743885

**Authors:** Arev P. Sümer, Leonardo N. M. Iasi, Alba Bossoms Mesa, Viviane Slon, Elena Essel, Mateja Hajdinjak, Julia Zorn, Anna Schmidt, Sarah Nagel, Birgit Nickel, Bence Viola, Rustam Ziganshin, Alexandra Buzhilova, Anatoly Derevianko, Svante Pääbo, Benjamin M. Peter

## Abstract

The Teshik-Tash 1 child whose remains were found in Uzbekistan represents the southeastern-most extent of the known Neandertal range, providing an important link with the better studied Caucasus and Altai Mountain ranges. However, due to poor DNA preservation, studying the genetics of Teshik Tash 1 has remained elusive. Here we present analyses of the nuclear DNA from the Teshik-Tash 1, from extracts that are highly contaminated with present-day human DNA. To achieve this, we developed a new computational method, *admixslug*, that jointly models contamination and population relationships, in order to infer the relationship of a target individual from which only low-quality nuclear DNA is available, to high-quality archaic human genomes. After validating *admixslug*, we show that Teshik-Tash 1 is genetically more similar to later Neandertals from Western Eurasia than to older Neandertals from the Altai Mountains. We estimate that Teshik-Tash 1 split from the Western Eurasian lineage between 80,000 and 100,000 years ago. Despite the geographical proximity of Teshik-Tash 1 to the Denisovan range, we find no evidence for Denisovan ancestry in his genome. Our results demonstrate that *admixslug* enables the study of archaic human specimens in cases where DNA preservation was previously considered too poor for population genetic analyses.

## Main

Neandertals occupied most of Western Eurasia throughout the Middle Palaeolithic before their disappearance around 40 thousand years before present (ka)^1–5^. Two genetically distinct Neandertal lineages have been described previously: the Altai lineage represented by two high-coverage genomes from Denisova Cave, Russia (Neandertals D5 and D17), both older than 100 ka^4,6^, and a Western Eurasian lineage represented by the high-coverage genomes of ∼45 ka Neandertals from Vindija Cave, Croatia^3^ (Vi33.19) and Goyet, Belgium (GN1)^7^, and a ∼80 ka Neandertal from Chagyrskaya Cave, Russia (Chag 8)^8^. All other sequenced Neandertal genomes to date from Europe, Western Asia, as well as from Chagyrskaya and Okladnikov Caves, belong to the Western Eurasian lineage^2,3,5,8–11^ that persisted until the disappearance of Neandertals from the archaeological record. These two lineages are estimated to have split from one another ∼140 ka ago^3,8^.

Due to scarce DNA preservation, genetic studies of Neandertals from the southern part of their range are rare^9,12^, and in particular the relationship of the Neandertals in the south-east to others remains poorly understood^13^. Teshik-Tash Cave in Uzbekistan represents a site at the southeastern-most extent of the Neandertal range. It is more than 2,000 km away from the nearest Neandertal sites at Mezmaiskaya Cave in the Caucasus and Denisova Cave and other sites in the Altai Mountains from which genetic data have been generated (**Figure 1.A**). Importantly, Teshik-Tash is also located close to the areas possibly occupied by the Denisovans^14^. In 1938, the incomplete skeleton of an ∼8-12 year old Neandertal child, Teshik-Tash 1, was discovered in a shallow pit surrounded by Siberian ibex horn cores^15^. Because of the young age at death, morphological traits often used for biological sex determination had not yet formed. However, it has been suggested that the child was a male based on the robustness of the jaws and browridge^16^. Although the lithic assemblage found in association with Teshik-Tash 1 was distinct from the nearest Neandertal assemblages from which genetic information is available, such as the ones found in Chagyrskaya and Okladnikov Caves in the Altai Mountains^5,17^, Teshik-Tash 1 was assigned to the Middle Palaeolithic, with no direct dates on the specimen^15^.

**Figure 1:**
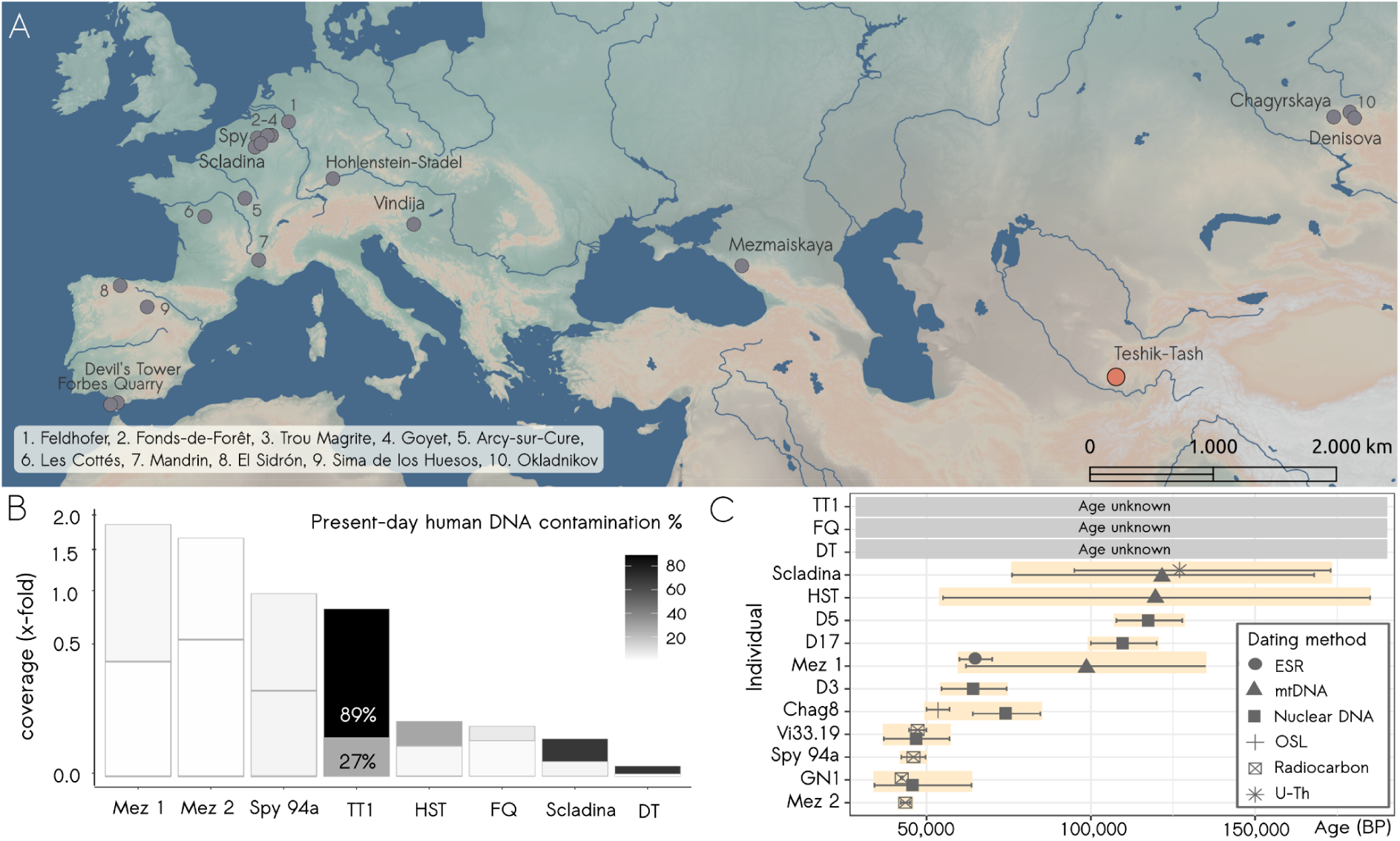
Geographical distribution and quality of the analysed Neandertal genomes. **A.** Map showing all the sites with available nuclear genomes from Neandertal remains. The sites with analyzed data in this study are labelled. All the other sites are numbered and listed at the left bottom of the panel. The base map was made with Natural Earth (www.naturalearthdata.com/). **B.** Average depth of coverage per individual in all sequences (top part on each bar) and only deaminated sequences (bottom part on each bar), and corresponding present-day human DNA contamination in these as estimated by *admixslug*. The values of the y-axis represent the fold-coverage (in square root scale to increase visibility of the lower coverage samples), and colour indicates the % of contamination. **C.** Estimated ages for the archaic humans mentioned or whose genomes are analysed in this study. Different shapes represent the methods used for obtaining the dates. Error bars represent the 95% CI. The y-axis lists the name of individuals while the x-axis represents the age estimates in years before present. abbreviations used in the figure are Mez 1: Mezmaiskaya 1, Mez 2: Mezmaiskaya 2, TT1: Teshik-Tash 1, HST: Hohlenstein-Stadel, FQ: Forbes Quarry, Scladina: Scladina_I-4A, DT: Devil’s Tower (**Extended Table 1**).

The relationship of Teshik-Tash 1 to other Neandertals has been proposed based on cranial morphology^18^ and a DNA sequence of the hypervariable region (HVR1) of the mitochondrial DNA (mtDNA) recovered from the specimen^12^. While the cranial morphology of Teshik-Tash 1 shows similarity to younger Western European Neandertals^18^, the partial mtDNA was found to be most similar to Neandertals older than 100 ka from Scladina in Belgium (Scladina_I-4A, a Western Eurasian Neandertal^10^)^12^, and to a lesser degree, two Neandertals from the Denisova Cave (D5 and D15)^6,19^.

To further understand the ancestry of the Teshik-Tash 1 child, we generated nuclear DNA data. We show that the recovered DNA is highly contaminated with the DNA of present-day humans, and thus cannot be analyzed using standard population genetic methods. We therefore develop *admixslug*, a method that co-estimates contamination and genetic relationships of the studied individual. We use *admixslug* to determine the genetic relationship of Teshik-Tash 1 to high-quality Neandertal and Denisovan genomes.

### High levels of contamination

We analyzed the DNA preservation in eight single-stranded libraries prepared from DNA extracted from Teshik-Tash 1 from a petrous bone in 2015 and from a long-bone fragment in 2019 (**SI3.1, ST.1**). All libraries had high-levels of present-day human DNA contamination (**Table SI3.2**). Despite this, mtDNA and/or nuclear DNA capture data from all libraries were produced as detailed in the Methods section and **SI3.1**.

We estimated between 49.4% and 97.5% contamination among the mtDNA sequences using the linear combination method, which is based on the derived allele frequencies in endogenous and contaminant sequences^20^ (**ST.2**). When restricting our analyses to deaminated sequences, estimates remained high (3.7-83.3%). Using four of the libraries that had less than 15% contamination among the deaminated sequences and requiring a minimum of four sequences with 75% agreement at each site, we called a partial consensus mtDNA sequence which covered ∼81% of the mitochondrial genome (3,075 of 16,569 sites were not covered) (**SI3.1**). We found that Teshik-Tash 1 mtDNA clustered with the Scladina Neandertal from Belgium (**Extended Figure 1**), as previously reported^12^.

Modern human nuclear DNA contamination ranged between 56% and 87% for all sequences, and up to 52% in the deaminated sequences, among the eight libraries (**Table SI3.6**). Merging the data from all libraries, we obtained a total of 0.84-fold genomic coverage using all sequences, and 0.044-fold coverage of deaminated sequences (**Figure 1.B**). The contamination estimates using the linear combination methods^5,7,10^ were 80.7% (95% CI: 79.6% - 81.8%), and 22.8% (95% CI: 19.7% - 26.3%) for these two data sets, respectively (**SI3.1**). To determine the genetic sex of the Teshik-Tash 1 child, we calculated the X-chromosome to autosome ratio, using the deaminated sequences. We found this ratio to be 0.51 (binomial 95% CI 0.45 - 0.57), indicating that the child was a male (**SI3.2**). This result is also supported by a recent proteomics study, based on amelogenin isoforms in the tooth enamel^21^ (**SI4.1**) (Buzhilova and Ziganshin, in press).

The recovered Teshik-Tash 1 genome is highly contaminated even among the deaminated DNA sequences, which is not uncommon for specimens excavated decades ago^9,10,20^. Therefore, the frequently used approach of restricting analyses to deaminated sequences that are assumed to have no or little contamination is not applicable for these types of data^9,22^. To address this limitation, we developed a new computational method, which we call *admixslug*.

### A new method for ultra-low coverage and highly contaminated genomes

The quality of sequenced Neandertal genomes is highly bimodal, with a small number of high-coverage genomes and a much larger number of genomes of low-coverage. Many of these are in addition highly contaminated by present-day human DNA. Here, we propose *admixslug*, a method to model the genetic relationship of a single low-quality target genome in the context of a set of high-coverage archaic and present-day human genomes, with the goal of co-estimating contamination and population relationships (**SI1.1, SI1.2**). The overall idea behind the model and the reason we do joint inference is that we can group the sequenced molecules in two ways. First, we can group them by molecular characteristics that have empirically been shown to impact contamination rates. Deaminated sequences typically have less contamination, longer human sequences are more likely to be contamination than shorter human sequences, shorter sequences are more likely to be bacterial than longer sequences, and techniques such as microsampling^2^ or bleach/phosphate pre-treatments^23^ yield libraries with different contamination rates. Second, we can group sequences by the genome to which they align best. If the target is a Neandertal, we would expect it to often share alleles with other Neandertals. Without contamination, these two groupings are orthogonal, i.e. molecular characteristics should not make molecules more Neandertal like, and the ancestry of a site should not inform deamination or molecule length. In contrast, when contamination is present, this is not true; deaminated reads, for example, tend to have lower contamination rates and thus will appear more Neandertal like, and we can use these differences to estimate differential contamination rates between deaminated and non-deaminated reads.

More formally, we use a mixture model with three main parts: First, a **contamination model** that estimates the probability for each sequence that it is a contaminant, in which case it is modelled as coming from a modern human population. The other possibility is classifying the molecule as endogenous, which implies it comes from the target individual, and is thus informative about its ancestry (**Figure 2**).

**Figure 2:**
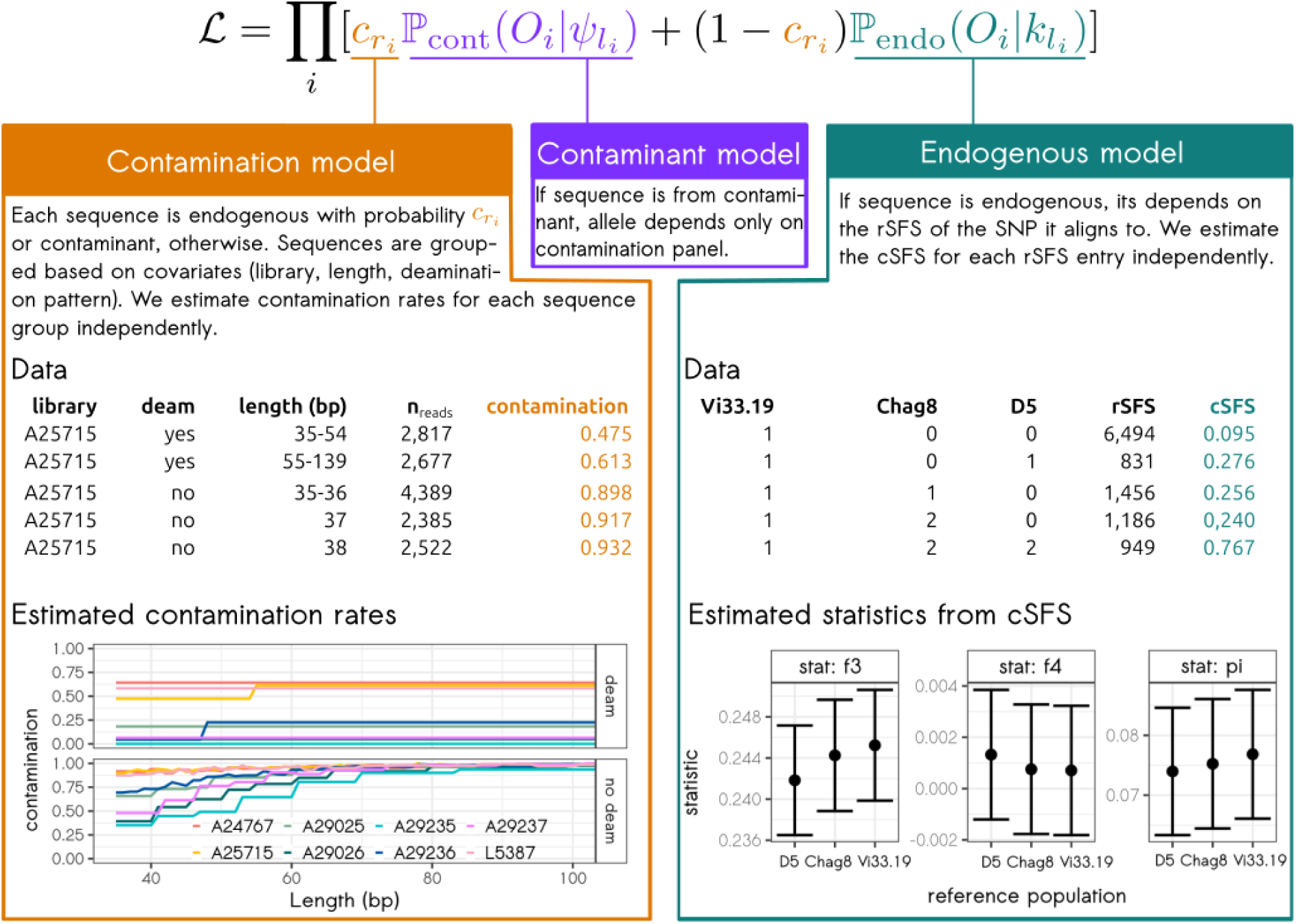
Schematic of the *admixslug* model. Top: the likelihood function is made up of three parts, the contamination model (orange) describing whether a sequence is contaminant or endogenous, a contaminant model (purple) for if the sequence comes from modern human contamination, and the endogenous model (green) if the sequence originates from the target individual. **Left:** We model contamination by subdividing the data into sequence groups based on shared characteristics (only 5 of 240 groups for Teshik-Tash 1 are shown), allowing us to estimate contamination from the data, coloured by library. **Right:** If data is endogenous, we use it to estimate population genetic statistics via the cSFS (only 5 of 135 entries are shown). Numbers for Vi33.19, Chag8 and D5 give the number of derived alleles for these Neandertals, rSFS and cSFS give the number of sites in that category and proportion of derived sites in Teshik Tash 1, respectively. Error bars represent one standard error.

We model the **endogenous** genome by assuming the high-quality reference data consists of *k* populations, where each population may be represented by a single individual. We first compute the *k*-dimensional reference site-frequency spectrum (rSFS), i.e., a tensor with each entry giving the number of sites that have a particular derived allele count in the reference data. If all reference individuals are part of the same population, this simplifies to the ordinary SFS. Many population genetic statistics within and between the reference populations, including heterozygosities, F_ST_, *f*-statistics and pairwise differences^24,25^ can be written in terms of the rSFS^26^. For each entry in the rSFS, we can then estimate the proportion of sites in this category for which the target individual has a derived allele. We call the resulting set of proportion, one per entry of the rSFS, the conditional site-frequency spectrum (cSFS). Since the rSFS is derived from putatively high-quality individuals, we assume it is known for inference. The entries τ_i_ of the cSFS, on the other hand, are unknown a priori and are inferred from the data (**Extended Figure 2**).

In **SI1.3** we show that many of the SFS-statistics can also be expressed in terms of the cSFS. The main exception are statistics that involve private sites in the target individual, as these are not captured by the cSFS. Thus, we cannot calculate F_ST_, the heterozygosity of the target individual or *f_2_*-statistics, but we can calculate *f_4_*, outgroup *f_3_*-statistics, F(A|B) and pairwise divergence between the target and the reference populations. Finally, our **contaminant model** incorporates modern human contamination by considering a second potential source of sequences that have derived alleles with probabilities given by a contamination panel.

In order to jointly estimate the parameters of the contamination and endogenous models, we use an accelerated expectation-maximization algorithm^27^, implemented in a program called *admixslug*. Specifically, we bin sequences into groups with similar characteristics (i.e., length, deamination, library), independent of where in the genome they align to, so that each bin contains approximately 2,000 sequences. We then estimate the contamination rates for each sequence group independently. The underlying idea is that without contamination, all sequences aligning to sites in the same rSFS-bin should have the same proportion of derived alleles, independent of molecular characteristics. Any systematic shift between sequences with different characteristics comes from contamination, allowing contamination rates to be estimated.

We extensively test our method on simulations and real data, including scenarios with very low coverages (down to 0.005x) and extremely high contamination rates (i.e. 95%) (**SI2**).

We evaluated the performance of *admixslug* on coalescent simulations^28^ using the demography in **Figure SI2.1**. *Admixslug* accurately estimated contamination in all cases, even when contamination occurred in both deaminated and non-deaminated sequences (**Figure 3**, **Figure SI2.4**). When comparing *f*-statistics calculated from *admixslug* to those based on ADMIXTOOLS^25^, we found that *admixslug’s* correction for contamination results in accurate estimates, while *f*-statistics calculated using ADMIXTOOLS^25^ are biased because they do not take contamination into account (**Figure 2**). The standard errors of the *f*-statistics from *admixslug* were consistently overestimated, in that 95% of the simulations were contained within 1.5 standard errors of the mean. Thus, we calibrate our confidence intervals (CIs) using simulations and report CIs based on ±1.5 standard errors for *f_4_*-statistics. We did not observe overestimation of standard errors for the F(A|B) statistics, and hence report ±2 standard errors (**SI2.1**).

**Figure 3:**
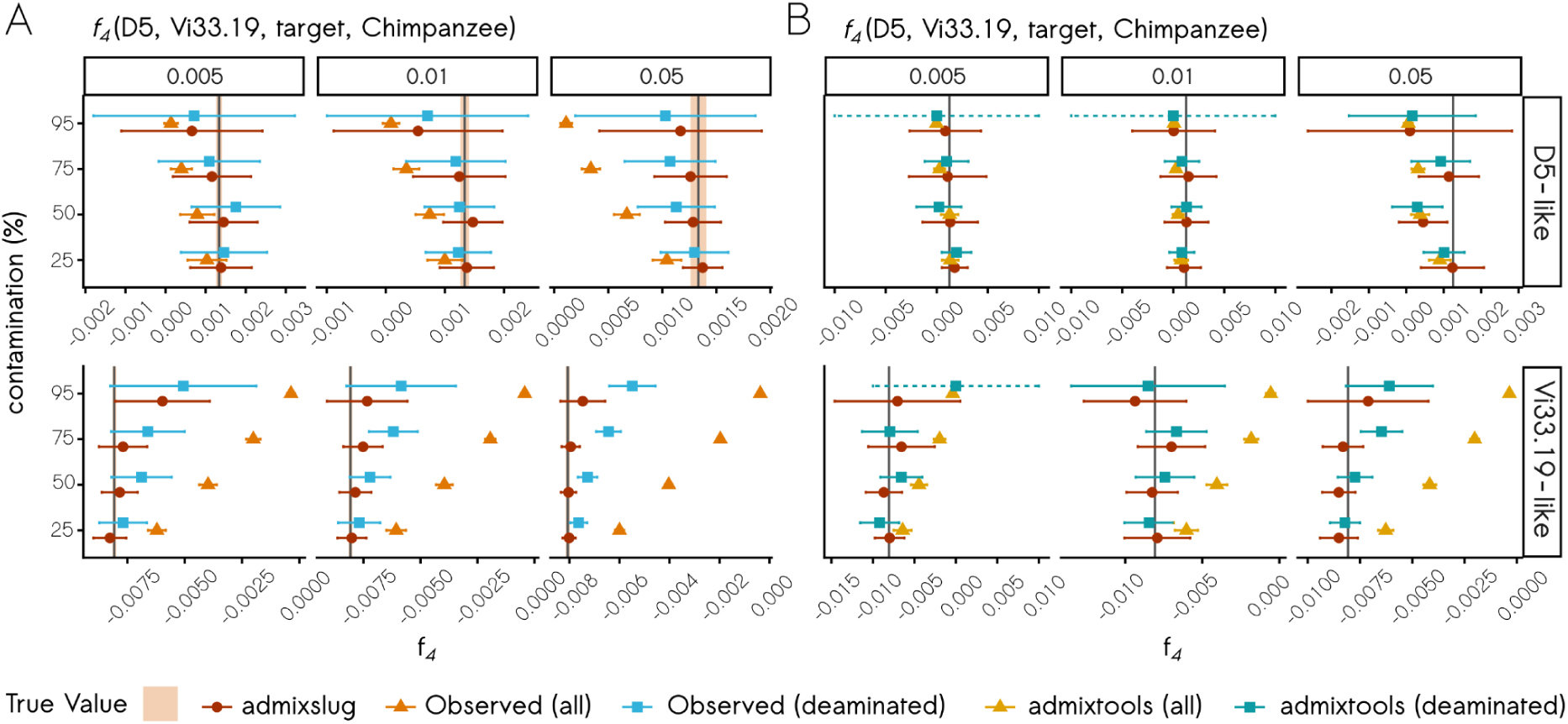
Comparisons of *f_4_(D5, Vi33.19, target, Chimpanzee)* statistics estimated on the simulated, downsampled and contaminated genomes, where deaminated sequences also carry present-day contamination. **A.** From 50 simulations in each scenario, comparison with direct estimates from simulated genotypes (Observed). The error bars correspond to the standard deviation. **B.** From a random simulation run in each scenario, comparison with ADMIXTOOLS. Simulated overall contamination shown on the y-axis, simulated Neandertal indicated on the right as row grids. Simulated average depth of coverage is on the x-axis panels on the top, while the x-axis at the bottom corresponds to the estimates of the *f_4_*-statistics. The colours of estimates represent the method used for each estimation, and the light orange area represents the central 95% range of true values obtained from repeated simulations The error bars on estimates indicate ±2 standard errors. The dotted error bars indicate intervals that extend beyond the displayed axis limits.

For tests on realistic data, we used the genome of the 45 ka-year-old Mezmaiskaya 2 Neandertal which has been sequenced to 1.7x coverage and has less than 1% modern human DNA contamination among all sequences^2^. To test the performance of *admixslug*, we introduced contamination up to 95% to the Mezmaiskaya 2 genome, and downsampled the data to various coverages (**SI2.2**). The lowest depth of coverage we tested was 0.005x, which corresponds to only ∼150 sequences stemming from the target Neandertal when contamination is 95%. In all cases, we find that the contamination rates estimated by *admixslug* overlap with the correct contamination rates (**Extended Figure 3**, **Figure SI2.12**), while simultaneously estimating contamination in groups of sequences. Thus, *admixslug* can also be used to estimate modern human DNA contamination in different groups of sequences and can be useful for filtering genomic data from archaic humans before performing analyses that are sensitive to present-day human DNA contamination.

### Teshik-Tash 1 was most similar to West Eurasian Neandertals

After establishing that *admixslug* performs well on highly contaminated and very low coverage Neandertal genomes, we applied it to the Teshik-Tash 1 nuclear DNA that we obtained through SNP enrichment. For comparison, we include seven Neandertal genomes of various quality in our *admixslug* analysis (**Figure 1.B, SI3.3**). These include the Neandertal genomes of Spy 94a and Mezmaiskaya 2^2^ that are from Neandertals younger than 50 ka years, the ∼65 ka years old Neandertal Mezmaiskaya 1^3^, two European Neandertals from Scladina and Hohlenstein-Stadel that are older than 100 ka years^10^, and two genomes from the Gibraltar Neandertals (Forbes Quarry and Devils Tower) that are of unclear age^9^ (**Figure 1.A, 1.C**). These genomes were all produced by whole-genome sequencing, from which we subsampled sequences covering the sites in the SNP enrichment ascertainment used for Teshik-Tash 1 (**SI3.1** and Skov et al., 2022^5^).

Contamination estimates generated by *admixslug* were slightly higher than those estimated by the linear combination method (**Figure 1.B, Figure S3.6, Table S3.7**). Using *admixslug*, we computed various *f_4_*-statistics (**Figure S3.8**), in particular *f_4_*(Vi33.19, D5, target, Chimpanzee) to measure if the target low-coverage Neandertal genome is closer to the Western Eurasian Neandertal lineage, represented by Vindija 33.19, or to the Altai lineage, represented by Neandertal D5. To estimate the relative order of split times from the Vindija 33.19 lineage within the Western Eurasian lineage without requiring direct comparisons between low-quality genomes, we used F(A|B) statistics^3,29^ comparing each Neandertal to the high-coverage Vindija 33.19 genome (**Figure 4.B, Extended Figure 4**).

**Figure 4:**
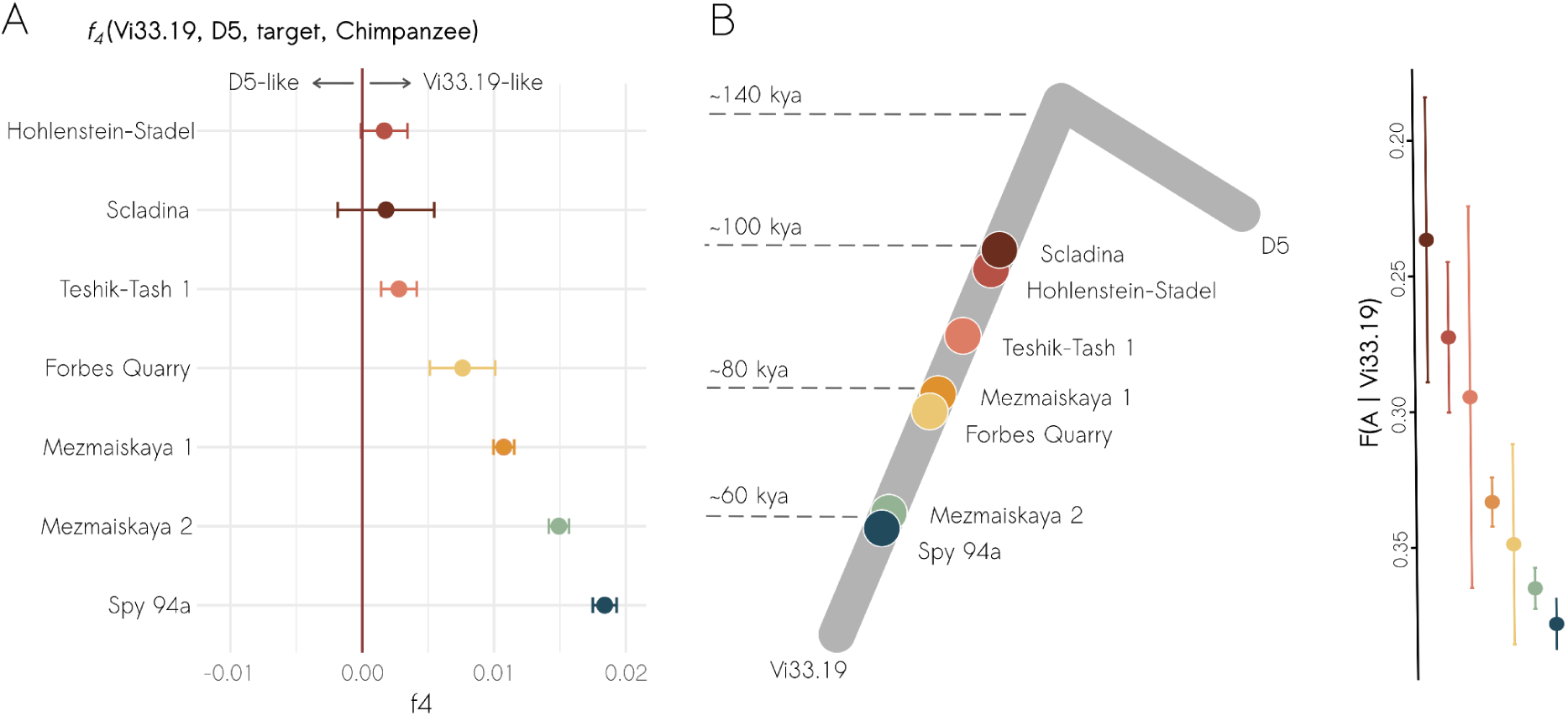
Population affinities and split times estimated using *admixslug*. **A.** *f_4_(*Vi33.19, D5, target, Chimpanzee) where the target is the low-coverage Neandertal genome listed on the y-axis. The error bars represent the 95% CI (1.5 standard errors). **B.** F(A|Vi33.19) statistics for the low-coverage Neandertals and schematic split time estimates inferred from this statistics. The error bars on the F(A|B) values correspond to the 95% CI (2 standard errors). Different colours represent the Neandertal genome used, and are consistent in the two panels.

For the three Neandertals with the highest quality data (Spy 94a, Mezmaiskaya 1 and 2), our analysis supports previous results indicating that they are genetically more similar to Vindija 33.19 than to D5, placing them on the Western Eurasian lineage. Spy 94a and Mezmaiskaya 2 have a shorter split time from the Vindija 33.19 lineage at ∼60 ka years ago^8,9^ than Mezmaiskaya 1 (∼80 ka years ago, **Figure 4**)^8,9^. We have less data from the Forbes Quarry Neandertal from Gibraltar, but overall our results are consistent with it also being on the Western Eurasian lineage. Previously it was not possible to perform *f*-statistics with the older and lower quality genomes of Scladina and Hohlenstein-Stadel and their closer genetic affinity to Vindija 33.19 compared to the D5 Neandertal was demonstrated using an indirect approach^10^. Here we replicate these results using *f_4_*-statistics calculated using *admixslug*, which similarly places them on the Western Eurasian lineage (**Figure 4.A**).

Our results indicate that Teshik-Tash 1 is genetically closer to the Vindija 33.19 Neandertal than to the D5 Neandertal, showing that he belonged to the Western Eurasian Neandertal lineage (**Figure 4**). This is in contrast to his mtDNA, which is closer to the mtDNA of the D5 Neandertal than to the Vindija 33.19 mtDNA (**Extended Figure 1**). A similar discrepancy between the relationships of the nuclear and mtDNA genomes was observed for the Scladina Neandertal^10^. Even though we cannot directly estimate how close Teshik-Tash 1 and Scladina are to each other as both have very little data, the *f_4_(*Vi33.19, D5, target, Chimpanzee) for Teshik-Tash 1 and Scladina respectively are 0.0028 (95% CI: 0.0014 - 0.0041) and 0.0018 (95% CI: -0.0019 - 0.0055), indicating that they have similar affinity to the Vindija 33.19 Neandertal (**Figure 4.A**). Furthermore, the *f_4_*(Vi33.19, Chagyrskaya 8, Teshik-Tash 1, Chimpanzee) is slightly positive, but not significantly so (**Figure S3.8**), suggesting that Teshik-Tash 1 is not on the Chagyskaya 8 lineage. Using F(A|B) statistics, we find that Teshik-Tash 1 splits from the Vindija 33.19 lineage after Scladina and Hohlenstein-Stadel, but before the Mezmaiskaya 1 and Forbes Quarry Neandertals. Taken together with the published split time estimates^9,10^, this would suggest that the lineage leading to Teshik-Tash 1 branched off from the lineage that leads to the Vindija 33.19 Neandertal between 100 ka and 80 ka years ago, although confidence intervals are large (**Figure 4.B**).

### No evidence of Denisovan ancestry in Teshik-Tash 1

Present-day populations of Central Asia carry higher levels of Denisovan ancestry when compared to West Asian and European populations, and recent archaeological studies have suggested that Denisovans might have been as far west as Tajikistan^30^, with this region acting as a corridor from Central Asia to the Altai Mountains, where Denisovans were present between 90 ka and 60 ka^31–34^. It is therefore possible that Teshik-Tash 1 could have carried Denisovan ancestry. Due to the low quality of the genome, it was not possible to directly identify genomic segments of Denisovan ancestry^35^. We instead used *f_4_-*statistics to compare the affinity of the high-coverage Denisovan genome to Teshik-Tash 1, as well as to three Neandertals that lack Denisovan ancestry^2,3^. More specifically, we computed the statistics *f_4_*(target, B, Denisovan D3, Chimpanzee), where B was either the high-coverage Vindija 33.19 or Chagyrskaya 8 Neandertal genome, and target was the low-coverage Teshik-Tash 1, Mezmaiskaya 1, Mezmaiskaya 2 or Spy 94a Neandertal genomes. If Teshik-Tash 1 carried Denisovan ancestry, we would expect these statistics to be positive. However, we do not see any difference in the estimates obtained for Teshik-Tash 1 and the other low-coverage target Neandertal genomes. Thus like other Neandertals on the Western Eurasian lineage, Teshik-Tash 1 did not carry any Denisovan ancestry (**Extended Figure 5**).

## Discussion

Until now, very little is known about genetic diversity of Neandertals between the Caucasus and the Altai Mountains (**Figure 1.A**)^2–4,6,8–12^. Here we present nuclear DNA from Teshik-Tash 1, to date the southeastern-most Neandertal to be genetically investigated. We develop a method that can be used to analyze archaic human data that is low-coverage and highly contaminated with present-day human DNA, such as we found in Teshik-Tash 1. Using our method, we show that Teshik-Tash 1 was on the Western Eurasian Neandertal lineage and likely split from the Vindija 33.19 branch after the split of Scladina and Hohlenstein-Stadel. This suggests that Teshik-Tash 1 was part of, or descended from, a widely distributed population of early Neandertals, which later gave rise to, or contributed to, the late Neandertals of Europe.

Spatially and perhaps also temporally, the closest high-coverage yielding Neandertal to Teshik-Tash 1 is the ∼80 ka years old Chagyrskaya 8^5,8^. To understand if Teshik-Tash 1 could be part of the Western Eurasian Neandertal population in Chagyrskaya, we directly tested if Teshik-Tash 1 is closer to Vindija 33.19 or Chagyrskaya 8. We observe slight but not significant affinity to the Vindija 33.19 Neandertal, suggesting that Teshik-Tash 1 was part of a population that is genetically distinct from the Chagyrskaya Neandertals (**Figure S3.8**).

There is direct evidence for Denisovan existence in the Altai Mountains between 90 ka to 60 ka years ago^32–34^, and their presence in the more southern regions of central Asia has been suggested^31^. Given these, it is interesting that we do not detect any Denisovan ancestry in the Teshik-Tash 1 genome. This could mean that the Denisovan range did not extend to the Teshik-Tash Cave, or that Teshik-Tash 1 individual and his ancestors did not overlap in time with the Denisovans in the region.

With the increased availability of sedimentary ancient DNA data, methods that utilize low-coverage data by comparing them to high-coverage references are becoming increasingly common^3,20,36,37^. With *admixslug*, we extend this to low-coverage, highly contaminated nuclear data from skeletal remains, allowing us to study genomes that have previously been deemed to be of too low-quality to be useful. By modelling contamination for different groups of molecules based on features such as deamination, sequence lengths, extraction methods or libraries, *admixslug* reveals how exactly contamination rates vary within a data set, and provides information on how best to filter data. We find that the length of molecules can be an important covariate associated with different contamination rates, although may be highly specific to each DNA library. Removing sequences above a length threshold could have a big impact on lowering contamination rates, and should be carefully considered before further analyses.

Compared to similar approaches such as DICE^38^, our method is more flexible because it allows contamination to vary between groups of sequences, and estimates summary statistics instead of a full demographic history. A drawback of this model-free approach is that it will perform less well if the endogenous and contaminant populations are very similar because the endogenous and contaminant models have very similar likelihoods. Another limitation is that we cannot estimate within-sample diversity of the low-coverage sample, preventing us from calculating F_ST_ and other summary statistics for which this is important. For this, more sophisticated genotype likelihood models would need to be extended to low-coverage, contaminated data^39^.

In summary, our results show that careful modeling of contamination by *admixslug* allows the genetic analysis of badly preserved DNA from archaic human remains. This opens the possibility to extend genetic analyses to many specimens where contamination by present-day human DNA has been a limiting factor.

## Extended Data

**Extended Table 1:**
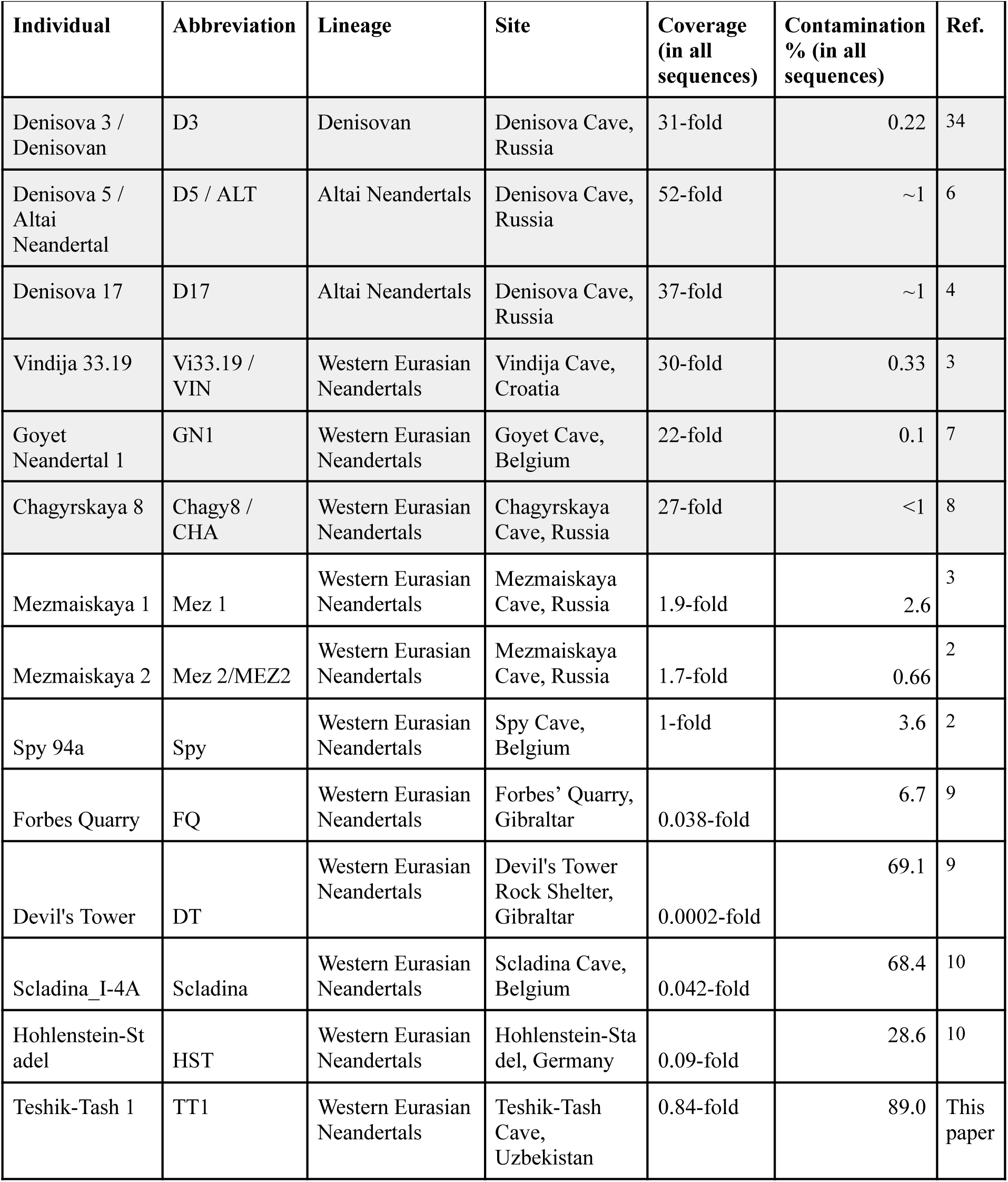
Archaic human genomes used or mentioned in this study. The average fold of coverage and present-day human DNA contamination reported for all sequences. See **Table S3.7** for contamination and coverage in the deaminated sequences. For low coverage genomes *admixslug* contamination estimates are reported. For high-coverage genomes (shaded in gray), contamination estimates are taken from the respective publications noted in the last column of the table.

| Individual | Abbreviation | Lineage | Site | Coverage<br>(in all<br>sequences) | Contamination<br>% (in all<br>sequences) | Ref. |
| --- | --- | --- | --- | --- | --- | --- |
| Denisova 3 /<br>Denisovan | D3 | Denisovan | Denisova Cave,<br>Russia | 31-fold | 0.22 | 34 |
| Denisova 5 /<br>Altai<br>Neandertal | D5 / ALT | Altai Neandertals | Denisova Cave,<br>Russia | 52-fold | ~1 | 6 |
| Denisova 17 | D17 | Altai Neandertals | Denisova Cave,<br>Russia | 37-fold | ~1 | 4 |
| Vindija 33.19 | Vi33.19 /<br>VIN | Western Eurasian<br>Neandertals | Vindija Cave,<br>Croatia | 30-fold | 0.33 | 3 |
| Goyet<br>Neandertal 1 | GN1 | Western Eurasian<br>Neandertals | Goyet Cave,<br>Belgium | 22-fold | 0.1 | 7 |
| Chagyrskaya 8 | Chagy8 /<br>CHA | Western Eurasian<br>Neandertals | Chagyrskaya<br>Cave, Russia | 27-fold | <1 | 8 |
| Mezmaiskaya 1 | Mez 1 | Western Eurasian<br>Neandertals | Mezmaiskaya<br>Cave, Russia | 1.9-fold | 2.6 | 3 |
| Mezmaiskaya 2 | Mez 2/MEZ2 | Western Eurasian<br>Neandertals | Mezmaiskaya<br>Cave, Russia | 1.7-fold | 0.66 | 2 |
| Spy 94a | Spy | Western Eurasian<br>Neandertals | Spy Cave,<br>Belgium | 1-fold | 3.6 | 2 |
| Forbes Quarry | FQ | Western Eurasian<br>Neandertals | Forbes' Quarry,<br>Gibraltar | 0.038-fold | 6.7 | 9 |
| Devil's Tower | DT | Western Eurasian<br>Neandertals | Devil's Tower<br>Rock Shelter,<br>Gibraltar | 0.0002-fold | 69.1 | 9 |
| Scladina_I-4A | Scladina | Western Eurasian<br>Neandertals | Scladina Cave,<br>Belgium | 0.042-fold | 68.4 | 10 |
| Hohlenstein-St<br>adel | HST | Western Eurasian<br>Neandertals | Hohlenstein-Sta<br>del, Germany | 0.09-fold | 28.6 | 10 |
| Teshik-Tash 1 | TT1 | Western Eurasian<br>Neandertals | Teshik-Tash<br>Cave,<br>Uzbekistan | 0.84-fold | 89.0 | This<br>paper |

**Extended Figure 1:**
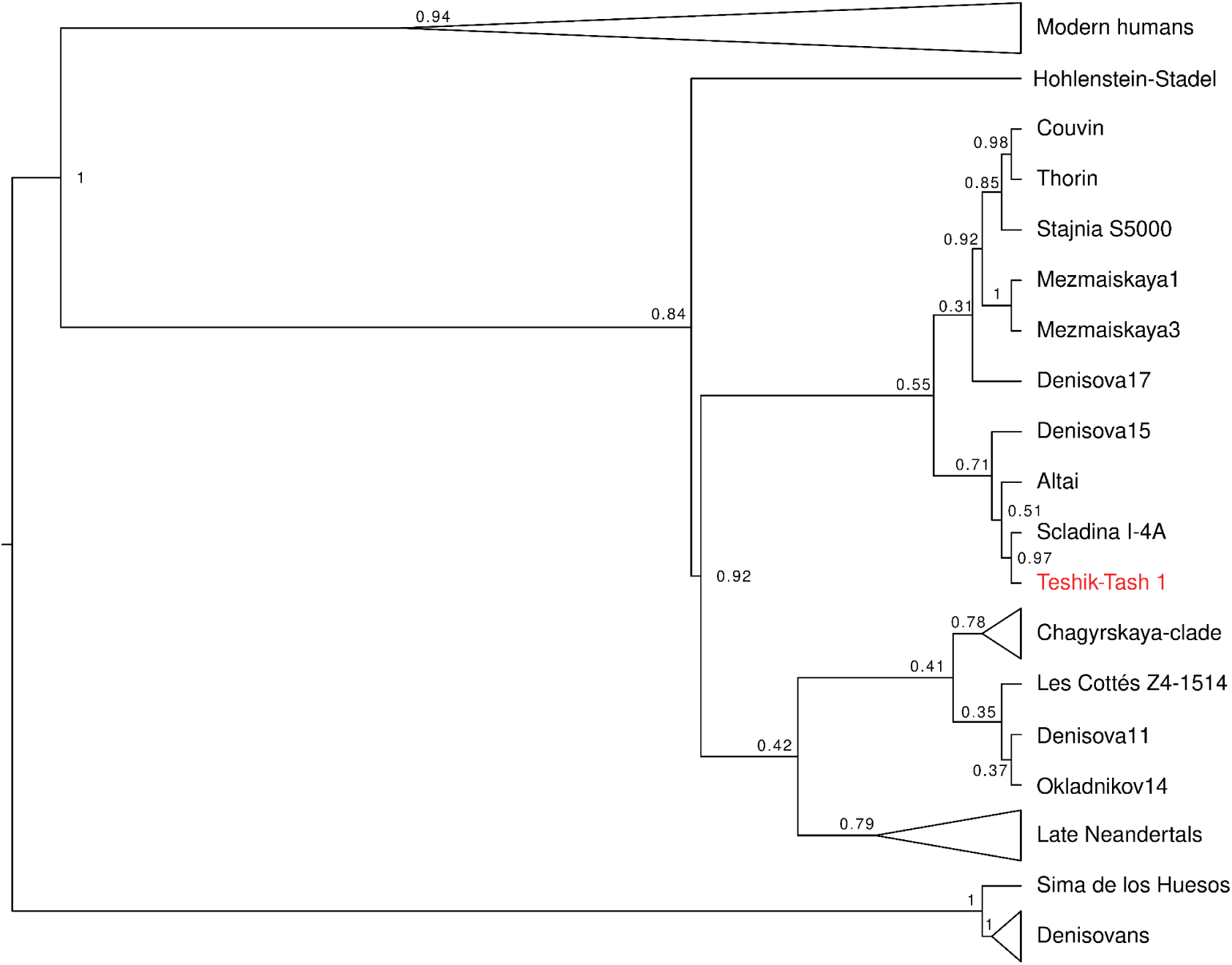
Mitochondrial DNA phylogeny. Maximum likelihood tree obtained using MEGA^40^, including the newly generated consensus for Teshik-Tash 1. The values on branches represent the bootstrap support for each branching, as calculated from 100 replicates. Clades for Denisovans, Late Neandertals, Chagyrskaya Neandertals and modern humans are collapsed for simplicity. A more detailed phylogeny can be found in the supplementary, **Figure S3.3.**

**Extended Figure 2:**
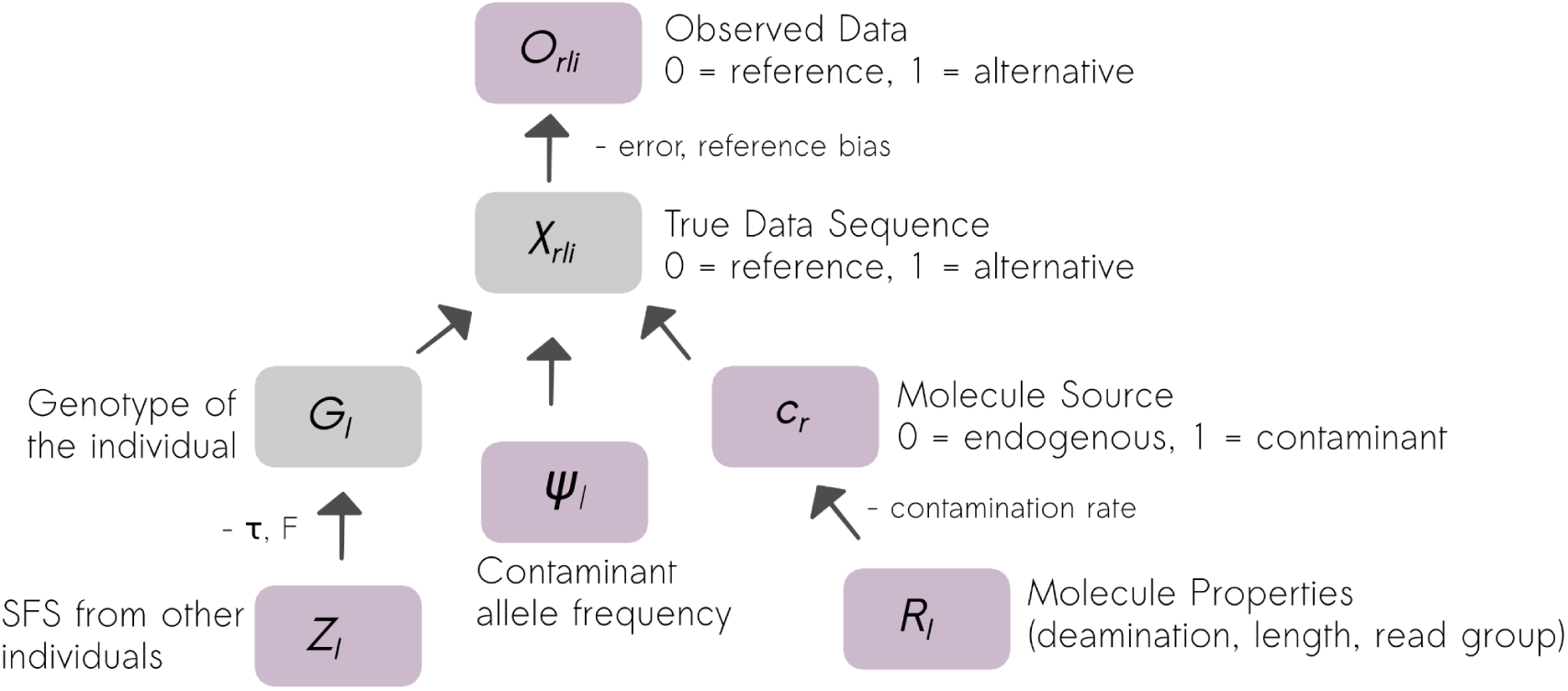
Schematic representation of the method. The parameters in the violet boxes are known, or obtained from the reference files while the ones in gray boxes are the latent states. The parameters next to the arrows between boxes are estimated by *admixslug* and used for inference.

**Extended Figure 3:**
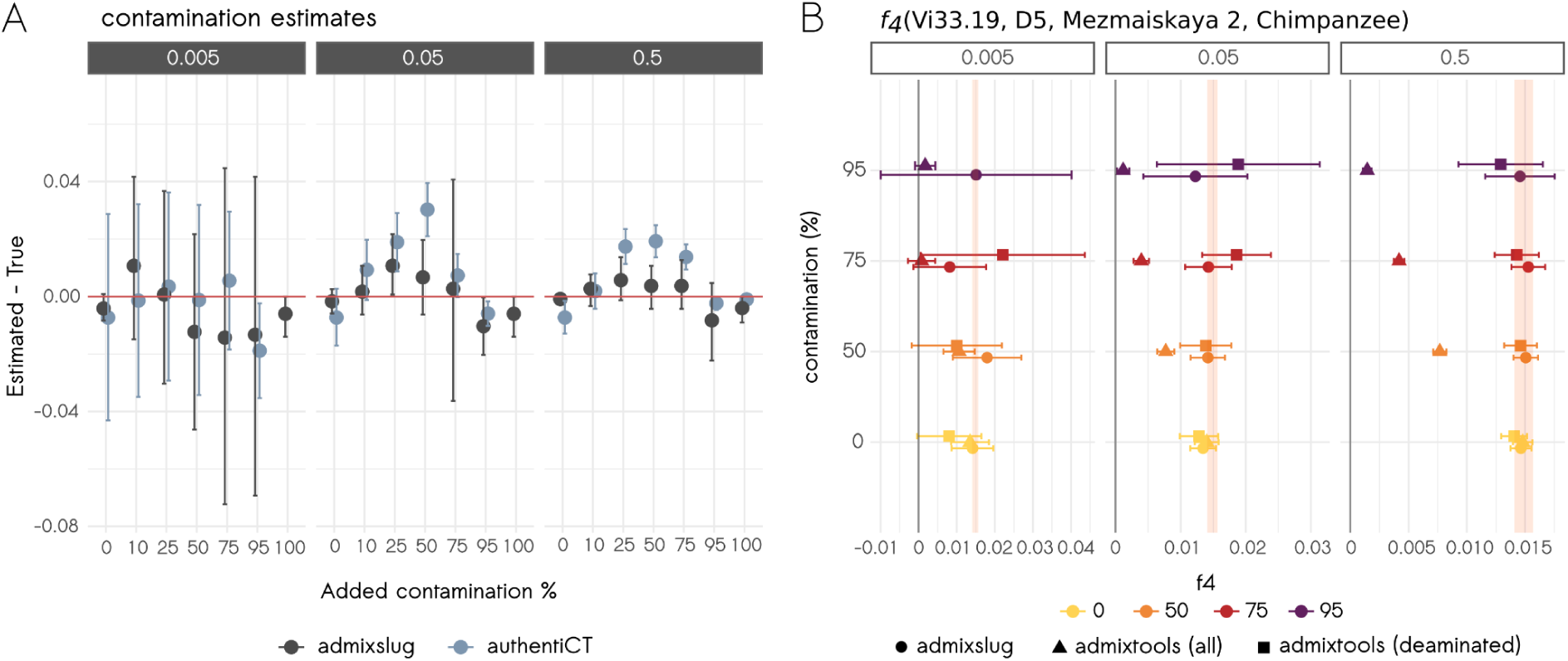
Testing *admixslug* on real data. We use downsampled and present-day human DNA contamination added versions of the Mezmaiskaya 2 Neandertal genome^2^ to test **A.** Contamination estimates of *admixslug*, in comparison to AuthentiCT^41^. The red horizontal line shows the level that estimated contamination is equal to the correct contamination. The y-axis measures the proportion of deviation from the correct values, while the x-axis shows the percentage of the added contamination. Column grids on the top indicated the average downsampled coverage, and colour of the points stand for the method used. Error bars correspond to the 2 standard errors. **B.** *f_4_*(Vi33.19, D5, target, Chimpanzee) where the target is downsampled and contaminated versions of the Mezmaiskaya 2 genome. The colours and shapes of the points correspond to added contamination and method used for estimation, respectively. Error bars represent the 95% CI (1.5 standard errors), and the orange area corresponds to the true value of the statistics estimated from the original Mezmaiskaya 2 genome that has no contamination.

**Extended Figure 4:**
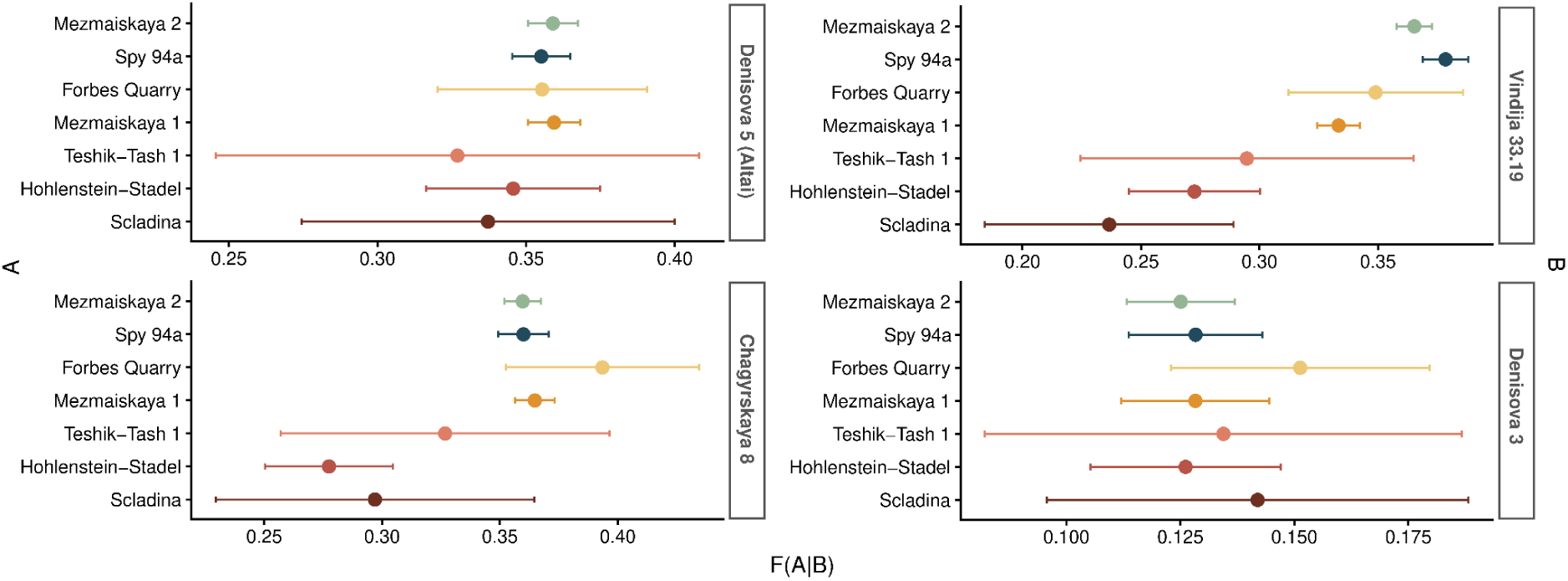
F(A|B) statistics. The individual whose genome used in position A is shown on the y-axis while in the high-coverage genomes in position B are indicated as the row grids on the right side of each plot. The values of F(A|B) statistics are shown on the x-axis. The colours are the same as those used in the main figures and correspond to different individuals. Error bars represent the 95% CI, and correspond to two standard errors.

**Extended Figure 5:**
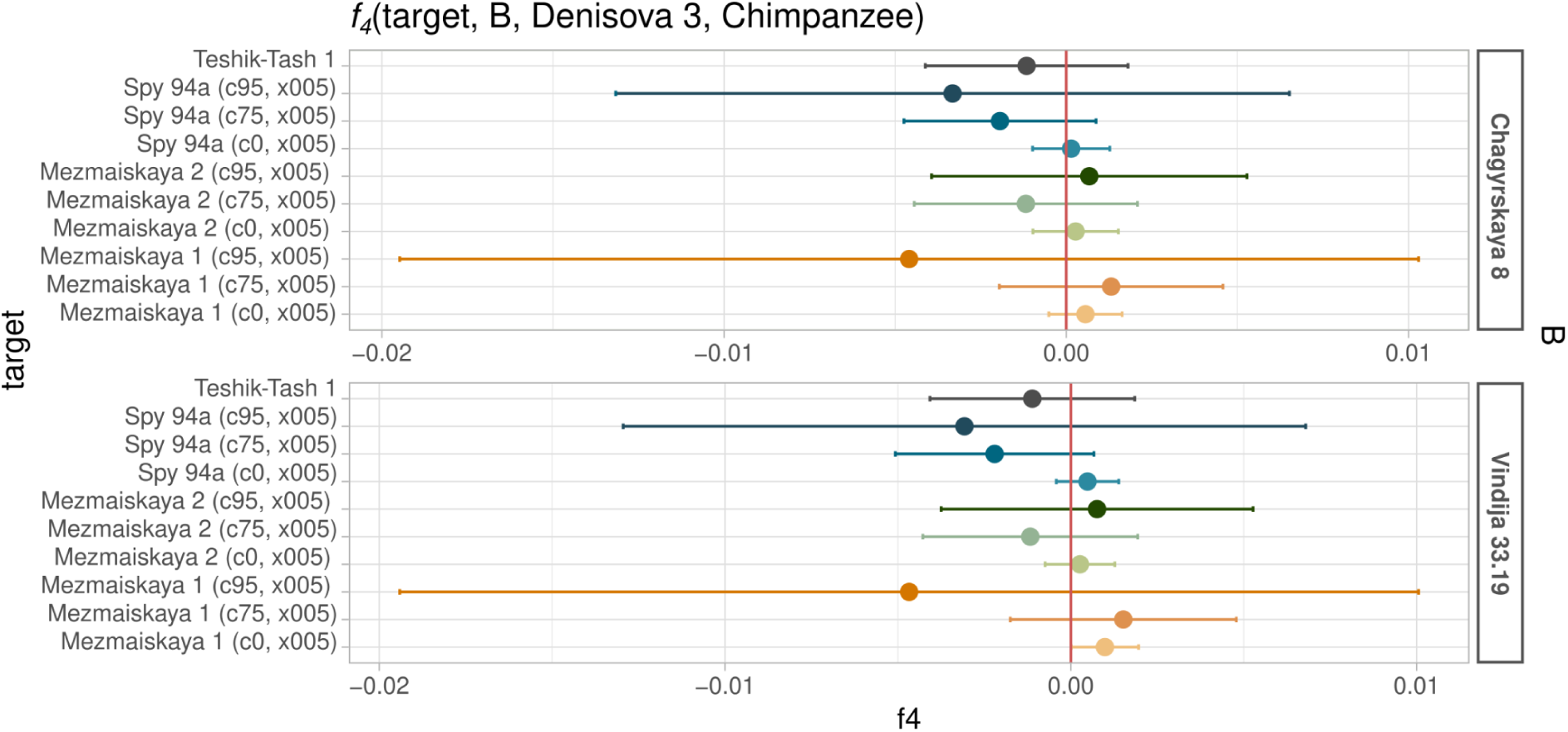
*f_4_*-statistics to investigate Denisovan ancestry in Teshik-Tash 1. The statistics *f_4_*(target, B, Denisova 3, Chimpanzee) measures the affinity of the low-coverage target genome to the Denisovan genome when compared to B, which is one of the two high-coverage Neandertal genomes indicated on the right side of the plot, as row grids. The error bars represent the 95% CI, corresponding to 1.5 standard errors. The genome in position target is shown on the y-axis, and includes the downsampled and artificially contaminated genomes from three late Neandertals in comparison to the Teshik-Tash 1 genome.

## Methods

### admixslug

To be able to make population genetics inference from very low coverage and highly contaminated archaic human genomes such as the genome of Teshik-Tash 1, we developed a new method, which we implemented in a python program and called *admixslug*. It jointly estimates present-day human DNA contamination and the conditional SFS of a target genome by combining information from both sequenced molecules (i.e. read groups, read lengths, deamination status etc.) and relatedness to other populations (genomes provided in a reference file) (see supplementary section **SI1.1**). For complete mathematical details of the admixslug model, its implementation and how to calculate downstream statistics, see supplementary sections **SI1.2** and **SI1.3**.

#### Testing admixslug

We tested our new method both on coalescent simulations and real data where we introduce present-day human DNA contamination. For both cases, we generated test scenarios with varying depth of coverage (1x, 0.5x, 0.1x, 0.05x, 0.01x and 0.005x fold) and DNA contamination levels (10%, 25%, 50%, 75%, 95%). For our tests on simulations, we further included a complex case where there was up to 30% contamination in the deaminated sequences.

To test *admixslug* on simulated Neandertal genomes, we used *msprime* (version 1.2.0)^28^. We used the demography detailed in the supplementary section **SI2.1** to sample two target individuals: one from the lineage that leads to the Neandertal D5 at 70 ka years ago (D5-like), and one from the Mezmaiskaya 2 lineage 40 ka years ago (Vi33.19-like). We simulated diploid samples with 20 chromosomes per individual, where each chromosome had a length of 50 Mb. Recombination rate was 10^-8^ per base pair per generation, and a per base pair per generation mutation rate was 10^-8^, with a generation time of 25 years. We mimicked the ascertainment scheme used for capturing data from the Teshik-Tash 1 libraries, which was introduced in Skov et al., 2022^5^. All simulated reads had a minimum read length of 30bp. We simulated two main scenarios for both the D5-like and Vi33.19-like Neandertal genomes: (1) where contamination was only found in non-deaminated sequences, and (2) contamination also in the deaminated sequences. We ran 50 simulations with different levels of contamination and coverage, and tested the performance of *admixslug*, as well as ADMIXTOOLS^25^. Furthermore, we calibrated the 95% confidence interval in our *f_4_*-statistics, using all different scenarios we simulated.

We tested *admixslug* on real data using the shotgun sequenced low-coverage genome of Mezmaiskaya 2^2^ as the base line, and adding various levels of contamination and subsequently downsampling the coverage, as detailed in supplementary section **SI2.2**. In these analyses we also use the sites ascertained for capture^5^, for comparability with the Teshik-Tash 1 genome and simulations. While testing *admixslug’s* contamination model, we used AuthentiCT^41^ for comparison. Similarly, ADMIXTOOLS^25^ was used for comparison in the tests for *f*-statistics.

### Sampling and DNA extraction

Two skeletal remains from the Teshik-Tash 1 individual, a petrous bone and a long bone fragment, were sampled for ancient DNA analyses (**Table SI3.1**). After removing a thin layer of surface material, we collected seven subsamples weighted between 5.9mg and 31.2mg. The petrous part of the left temporal bone was sampled in Moscow in 2015 using a sterile dentistry drill. An aliquot of 10.7mg was washed with a sodium phosphate buffer in order to remove contamination^23^ and another of 10mg was not treated before extraction. DNA was then extracted using a silica-based method optimized for the retrieval of short DNA molecules^42^. An aliquot of this extract was converted into a single-stranded DNA library^43^, and barcoded with a pair of unique indexes^44^. For the remaining five subsamples, we followed the protocol described in Rohland et al., 2018^45^ on an automated handling platform and using binding buffer “D”, without any treatment. We then used an automated version of the single-stranded DNA library preparation protocol^46^. Three types of data were generated from these libraries: shallow shotgun screening data, mitochondrial DNA (mtDNA) capture data and nuclear DNA capture data.

### Sequencing and filtering

Shallow shotgun sequencing data was obtained after library amplification by using a 75bp paired-end read configuration, at the MPI-EVA on HiSeq4000 and MiSeq platforms, through sequencing to a depth of 3-5 million reads. We demultiplexed the resulting sequences based on perfect matching of the expected index combinations, and aligned the sequences to hg19 human reference genome using BWA (version 0.5.10-evan.9-1-g44db244, https://github.com/mpieva/network-aware-bwa) with the ancient DNA parameters (“-n 0.01 –o 2 –l 16500”)^34^. We removed PCR duplicates using bam-rmdup (https://github.com/mpieva/biohazard-tools/), and filtered the sequences for mapping quality of 25 and minimum length cutoff of 30bp.

mtDNA data was obtained following protocols optimized for ancient DNA^47^. We used two rounds of automated in-solution hybridisation capture and a probe-set that tiled the revised Cambridge Reference Sequence (rCRS,NC_01290)^48^. Subsequently, capture libraries were sequenced at the Core Unit facilities on MiSeq and HiSeq platforms. We merged and trimmed the raw reads using (v.1.2.18, https://bioinf.eva.mpg.de/leehom/)^49^, and mapped the resulting sequences to rCRS reference using BWA version 0.5.10^50^ with ancient DNA parameters^34^. We demultiplex the sequences, removed PCR duplicates, and filtered for length and mapping quality the same way we did for the screening data.

To obtain nuclear DNA from Teshik-Tash 1, we enriched all eight libraries using an array designed specifically to capture nuclear variation in archaic populations^5^. The details of this ascertainment can be found in supplementary section **SI3.1**. We sequenced the captured libraries at the Core Unit facilities, on HiSeq and MiSeq platforms. Demultiplexing and filtering were performed exactly as described for the shallow shotgun data obtained for screening. To minimize reference bias towards modern human alleles and to keep our data comparable with the previously published low-coverage capture data from Neandertals^5,9,10^, we followed the alignment strategy introduced in Peyrégne et al., 2019^10^. Specifically, we aligned our sequences to both the hg19 human reference genome and to a modified reference genome that carries an alternative archaic allele for each of the variants on the array. We merged both alignments per library, and kept all sequences with a mapping quality of 25 or higher, and a length of 30bp or longer (**Table SI3.4**).

### Mitochondrial DNA analysis

We estimated the levels of present-day human DNA contamination in each library using lineage-diagnostic positions^20^ and AuthentiCT^41^. In both cases, contamination estimates exceeded 30% for all libraries, hence we decided to use only deaminated sequences. Summary statistics and contamination estimates can be found in **ST.2.** We called the mtDNA consensus for Teshik-Tash 1 using the four libraries that had a minimum of 10% terminal deamination, and less than 15% contamination. We used only the deaminated sequences, and required minimum 4-fold coverage and 75% consensus support for calling the consensus, with an in-house perl script. We realigned our consensus sequences with a set of other reference mitochondrial genomes using *mafft* with 1,000 iterations (version 7.453^51^). We then investigated phylogenetic relationships by building a Maximum Likelihood (ML) tree using 100 bootstraps, with MEGA^40^ (version 10.1.7). In addition, we used Kallisto^52^, an alternative method often used for sedimentary ancient DNA, to further explore the phylogenetic affinities of Teshik-Tash 1 to other mtDNA genomes (**SI3.1**).

### Sex determination for Teshik-Tash 1

We investigated the biological sex of Teshik-Tash 1 child using genetics (**SI3.2**). For this, we subsetted all nuclear capture data to sequences with deamination and estimated the ratio of sequences on the X-chromosome and autosomes (**SI3.2** and **Table ST.4**). Because males and females carry different numbers of X-chromosomes, this ratio can be used to determine the sex of the individual^53^. Because the capture ascertainment did not include any sites on the Y-chromosome, we used the shallow-shotgun screening data to check sequences that aligned to the Y-chromosome (**Table ST.5**). We provide additional support based on proteomics, summarizing results from an upcoming study (**SI4**) (Buzhilova and Ziganshin, in review).

#### Nuclear DNA analysis using *admixslug*

We compiled a set of published Neandertal genomes to compare with Teshik-Tash 1, including low-coverage genome-wide data from two European Neandertals that lived ∼100 ka years ago (from Hohlenstein-Stadel and Scladina^10^), two Neandertals from Gibraltar (Forbes Quarry and Devil’s Tower^9^), the ∼65 ka years old Mezmaiskaya 1 Neandertal^3^, and two late Neandertals from Spy and Mezmaiskaya^2^ (**SI3.3**). We subsetted these published low-coverage genomes to the positions targeted by the capture array we used for Teshik-Tash 1. We ran *admixslug* on all genomes with parameters:

admixslug --in input_file --states VIN CHA ALT DEN --ref reference_file --ancestral PAN --cont-id EUR -o output_name --ptol 0.001 --ll-tol 0.01 --max-iter 100 --filter-ancestral --len-bin-size 2000 --jk-resamples 500 --output-jk-sfs --output-fstats

To understand if Teshik-Tash 1 was closer to the Vindija Neandertal (Vi33.19) or to the Altai Neandertal (D5), we used *f_4_*(Vi33.19, D5, Teshik-Tash 1, Chimpanzee) as calculated by *admixslug* (see **SI3.4)**. To investigate if Teshik-Tash 1 carries ancestry from Denisovans, we removed ALT (Altai Neandertal, D5) from the `--states`. This was done to avoid potential biases because it is known that the Altai Neandertal carries Denisovan ancestry^4,35^. We were then interested in two statistics, *f_4_*(Denisova 3, B, target, Chimpanzee) to see if Denisova 3^34^ or the other high-coverage in position B is closer to the target (Teshik-Tash 1 and the other Neandertal genomes in the comparative set) and *f_4_*(target, B, Denisova 3, Chimpanzee) to check if target or B is closer to the Denisova 3 genome. We use the Chimpanzee reference genome panTro6 for both analyses.

We estimated the F(A|B) statistics directly from the output of *admixslug*, following **SI1.3.** F(A|B) statistics measures the proportion of heterozygous sites in a diploid genome *B* that share a derived allele with a pseudohaploid genome *A*^3,6,29^. We use F(target|Vi33.19) to approximate the split order of Teshik-Tash 1 and of other Neandertals in the comparative set from the high-coverage Vindija Neandertal (Vi33.19) genome (see **SI3.4**). The genomic coverage after subsetting for the capture ascertainment was too low for Devil’s Tower Neandertal (**SI3.4**), causing us to exclude this individual from further analyses.

## Data Availability

All newly reported ancient nuclear and mitochondrial DNA data is archived in the European Nucleotide Archive (accession no. PRJXXX).

## Code Availability

We provide the code for the coalescent simulations ran for testing *admixslug* in the GitHub repository https://github.com/LeonardoIasi/admixslug_simulations and *admixslug* is available in the GitHub repository https://github.com/BenjaminPeter/admixslug/.

## Acknowledgments

We thank MPI-EVA Core Unit, Johann Visagie and Rigo Schultz for their assistance with data production, recovery and processing. We thank Janet Kelso, Matthias Meyer and Stéphane Peyrégne for helpful discussions, and Janet Kelso also for her comments on the manuscript. The specimens were curated by Vitaliy M. Kharitonov, who recently passed away.

## Author Contributions

This study was designed by A.P.S., S.P. and B.M.P. Computational methodology was developed or tested by A.P.S., L.N.M.I. and B.M.P. Anthropological assessment or curation of skeletal material was performed by B.V., A.B. and A.D. Sample preparations and laboratory work were conducted by V.S., E.E., M.H., J.Z., A.S., S.N., B.N., and subsequent analyses were performed by A.P.S. and A.B.M. With input from all co-authors, A.P.S. and B.M.P. wrote the manuscript.

## Additional Information

Supplementary Information is available for this paper.

Correspondence and requests for materials should be addressed to Arev Pelin Sümer and/or Benjamin Marco Peter.

