## Supplementary Information for "A Method to Analyze Low-Quality Archaic Human Genomes and its Application to the Teshik-Tash 1 Neandertal"


**SI1: Admixslug Model** **2**
SI1.1 - Model overview 2
SI1.2 - Model details 4
SI1.3 - Calculating statistics 7
*f*-statistics 8
F(A|B) statistics 9
**SI2: Testing admixslug** **10**
SI2.1 - Model testing on simulations 10
Simulated demography and scenarios 10
Calculation of *f*-statistics 12
Testing the influence of the ascertainment 13
Testing the estimation of contamination 14
Validation of the *f*-statistics on simulations 15
$f_3$-statistics *15*
$f_4$-statistics *16*
Validation of the F(A|B) statistics on simulations 19
Single-estimate performance and confidence interval calibration 22
SI2.2 - Model testing on real data 24
Contamination 25
*f*-statistics *26*
$f_3$-statistics *26*
$f_4$-statistics *27*
Comparison with ADMIXTOOLS 28
Ascertainment used for capture 30
**SI3: Application on Teshik-Tash 1** **33**
SI3.1 - Data generation 37
Screening 38
Mitochondrial Capture 40
Nuclear Capture 43
SI3.2 - Genetic sex of Teshik-Tash 1 47
SI3.3 - Comparative data set for nuclear DNA analyses 47
SI3.4 - Admixslug results 48
Present-day human DNA contamination 48
Unbiased *f*-statistics and F(A|B) 51
**SI4: Sex determination based on proteomics** **56**
**References** **57**

### SI1: Admixslug Model

#### SI1.1 - Model overview

Here, we present a graphical model for the joint estimation of contamination and the conditional site-frequency spectrum, implemented in a program called *admixslug*. The model aims to combine information from both sequenced molecules from the target genome (i.e. read groups, read lengths, deamination status etc.) and relatedness to other populations (genomes provided in a reference file).

We assume we have aligned data produced by next-generation sequencing, from $L$ SNPs of a target individual, possibly low-coverage. Further, we assume a set of one or more of high-quality reference populations with genotypes at these loci, we also assume we have an ancestral allele. Inference is done in the context of the *m*-dimensional reference site-frequency spectrum (SFS), where *m* is the number of reference populations.

For example, if we have two reference populations, one represented by one, and the other by two diploid individuals, then the possible genotypes for population 1 would be 0, 1, 2 derived alleles, and for the second one 0, 1, 2, 3, 4 derived alleles. The resulting SFS is the combination of all possible combinations of alleles from the two populations, in this case there would be 3 x 5 = 15 entries (possibly -2, if we remove monomorphic entries). Inference is done using a conditional SFS, where we calculate the SFS of the target individual, conditional on the states of the reference SFS. Conditional SFS have been used elsewhere[1,2] but often just with one reference population, and a large target population. Here, we assume the target population is represented by just one individual, and for each entry in the reference the target individual at these loci.

Specifically, we estimate the proportion of derived alleles of the target individual for each entry of the conditional SFS. As we will show, this summary can be used to calculate several statistics of interest, including *f*-statistics[3,4], pairwise differences and F(A|B) statistics.

For inference, we assume that for each SNP we have zero or more reads, some come from the individual of interest, and some from a contaminant population. Reads are subdivided into $R$ disjoint read groups that have distinct contamination rates. Read groups may represent different libraries or other molecular properties such as molecule length or deamination patterns.

Given this input data, we first model three sets latent states:

1. The genotype of the target individual at a particular SNP $G_l$)
2. The source of a sequenced molecule (endogenous vs contaminant, $C_rli$)
3. The base of the sequenced molecule before sequencing ($X_rli$)

And we have the following known observations:

1. $Z_l$: The SFS-entry SNP l corresponds to
2. $Psi_l$: The derived allele frequency in a contamination panel
3. $R_li$: The read group of the i-th read at SNP l
4. $O_rli$: The observed base at the i-th read from read group r at SNP l

Note that some observations and variables are joint for each SNP ($G_l$, $Z_l$, $Psi_l$), while others are defined for each read ($C_rli$, $X_rli$, $O_rli$, $R_li$).

We are primarily interested in estimating the latent state $G$, the contamination rate for each read group, and the proportion of derived allele for all SFS entries $\tau_k$.

In this section, we give the full mathematical details of the admixslug model and its implementation.

#### Notation overview

Throughout, the notation is as follows:

- $R$ is the number of read groups
- $L$ is the number of biallelic SNP loci
- $K$ is the number of SFS-bins, which are entries in the reference SFS.
- $n_{rl}$ the number of reads of read group $r$ at SNP $l$.
- $O_{rli}$ is the $i$-th read from read group $r$ at SNP $l$
- $G_l$ is the genotype at SNP $l$, codified by the number of derived alleles
- $\psi_l$ is the derived allele frequency in a contaminant panel at locus $l$
- $Z_l$ is the set of all SFS entries, and the SFS-entry corresponding to SNP $l$
- $X_{rli}$ is an indicator that is one if the true allele corresponding to $O_{rli}$ is derived, zero if it is ancestral
- $C_{rli}$ an indicator that is 1 if the read corresponding to $X_{rli}$ is of contaminant origin, and 0 if it is endogenous
- $F_k$ a parameter estimating coalescence since gene flow for SNP in SFS-entry $k$.
- $\tau_k$ the proportion of derived alleles in SFS-entry $k$.
- $c_r$ proportion of contaminant reads in read group $r$
- $e, b$ error rate, and reference bias
- $W_l$
- $\theta = (c_r, e, b, \tau_k, F_k)$, the set of all parameters to be estimated

In all cases, we use a bold capital letter to denote the set of all parameters, i.e. $\mathbf{G} = (G_l)$ is the set of all genotypes.

### SI1.2 - Model details

#### Error model

The error model relates the (unobserved) true state of the sequenced molecule $X_{lri}$ to the observed sequencing data $O_{lri}$, which denotes the $i$-th read of read group $r$ at locus $l$. We consider sequencing error, and reference bias.

$X$ is based on ancestral/derived alleles, whereas $O$ is codified by reference/alternative alleles. We further have a variable $W_l$ that is 1 if the reference allele is flipped, i.e.

$$W_l = \begin{cases} 0 & \text{if REF} = \text{Ancestral allele} \\ 1 & \text{if REF} = \text{Derived allele} \end{cases}$$

The eight possible cases are then:

We can think of $e$ as the sequencing error, and $b$ as the reference bias + sequencing error.

This is implemented in `bwd_p_o_given_x`, which calculates a matrix of size $R \times 2$ containing the entries $P(O_i|X_i = j)$, flattened over $r$ and $l$.

| Observed base $O_{rli}$ | base on seq $X_{rli}$ | flipped $W_l$ | alleles | probability |
|---:|---:|---:|---|---:|
| 0 (ref) | 0 (anc) | 0 | ref = anc, alt = der | 1 - e |
| 1 (alt) | 0 (anc) | 0 | ref = anc, alt = der | e |
| 0 (ref) | 1 (der) | 0 | ref = anc, alt = der | b |
| 1 (alt) | 1 (der) | 0 | ref = anc, alt = der | 1 - b |
| 0 (ref) | 0 (anc) | 1 | ref = der, alt = anc | b |
| 1 (alt) | 0 (anc) | 1 | ref = der, alt = anc | 1 - b |
| 0 (ref) | 1 (der) | 1 | ref = der, alt = anc | 1 - e |
| 1 (alt) | 1 (der) | 1 | ref = der, alt = anc | e |

#### Sequence model

There are two possible origins for each sequence $X_{lri}$, it is either a contaminant, or endogenous. Let $C_{lri} = 1$ mean that $X_{lri}$ is contaminant, and $C_{lri} = 0$ mean that it is not. Furthermore, let $\psi_l$ be the **derived** allele frequency in a reference contamination panel, which we assume to be known. Let $G_l$ be the genotype of the target individual at SNP $l$, counted as the number of derived alleles (which is either 0, 1 or 2 in diploid individuals).

$$
\begin{aligned}
P(X_{lri} = 0|C_{lri} = 1, \psi_l) &= 1 - \psi_l \\
P(X_{lri} = 1|C_{lri} = 1, \psi_l) &= \psi_l \\
P(X_{lri} = 0|C_{lri} = 0, G_l) &= 1 - \frac{G_l}{2} \\
P(X_{lri} = 1|C_{lri} = 0, G_l) &= \frac{G_l}{2}
\end{aligned} \tag{1}
$$

In haploid regions (or more generally, in regions with different ploidy) we divide $G_l$ by the ploidy instead.

#### Contamination model

For read-group $r$, the probability that a read from that read group is a contaminant is

$$
\begin{aligned}
P(C_{lri} = 0|c_r) &= 1 - c_r \\
P(C_{lri} = 1|c_r) &= c_r
\end{aligned} \tag{2}
$$

independent of the locus.

#### Genotype model

We estimate the genotype of a locus given the conditional-SFS entry $Z_l = k$, $F_k$ is the probability that both alleles are IBD, and $\tau_k$ is the probability that the individual has a derived allele at position $k$ Thus

$$
\begin{aligned}
P(G_l = 0|Z_l = k, \tau_k, F_k) &= F_k(1 - \tau_k) + (1 - F_k)(1 - \tau_k)^2 \\
P(G_l = 1|Z_l = k, \tau_k, F_k) &= 2(1 - F_k)\tau_k(1 - \tau_k) \\
P(G_l = 2|Z_l = k, \tau_k, F_k) &= F_k\tau_k + (1 - F_k)\tau_k^2
\end{aligned} \tag{3}
$$

Estimating all the $\tau_k$ is one of the main goals of `admixslug`, as they can be used to calculate $F$-statistics and other quantities of interest.

For example, if we compare with the Altai Neandertal and Denisova 3 genomes, we would have the following $Z$ states:

This is implemented in `p_gt_diploid` and tested in `tests/test_slug.py:test_slug_p_gt_diploid`. $\tau$ estimates can be found in the output files .sfs.xz and .jksfs.xz.

For comparison with genotype-based methods, we also implement a `--gtmode` flag that allows the user to disable the genotype likelihood and contamination models and directly supply the

| $Z_l = k$ | Altai(D5) | Denisovan (D3) |
|---|---|---|
| 0 | 0 | 0 |
| 1 | 0 | 1 |
| 2 | 0 | 2 |
| 3 | 1 | 0 |
| 4 | 1 | 1 |
| 5 | 1 | 2 |
| 6 | 2 | 0 |
| 7 | 2 | 1 |
| 8 | 2 | 2 |

genotypes, in which case this is the only part of the model that is retained. To run this mode, in addition to `--gtmode`, `--rrgt`, `--dont-est-error` and `--e0 0` flags are used.

#### Likelihood and parameter estimation

##### Complete Data Likelihood

As in most graphical models, optimizing the exact likelihood is intractable because we would need to jointly optimize all parameters.

We therefore use an EM-algorithm to estimate parameters. We proceed by calculating the complete-data likelihood (where we assume the latent variables) are known, and then derived the $Q$-function for estimating parameters.

The complete data likelihood can be written as

$$\begin{aligned}\log P(\mathbf{O}, \mathbf{X}, \mathbf{C}, \mathbf{G}|\theta, \psi, \mathbf{Z}) = &\sum_{lri} \log P(O_{lri}|X_{lri}, e, b)\\ &+ \sum_{lri} \log P(C_{lri}|c_r)\\ &+ \sum_{lri} \log P(X_{lri}|C_{lri}, \psi_l, G_l)\\ &+ \sum_{l} \log P(G_l|Z_l, \tau_k, F_k)\end{aligned}$$

The corresponding Q-function is

$$\begin{aligned}Q(\theta|\theta') &= \mathbb{E}\big[\log P(\mathbf{O}, \mathbf{X}, \mathbf{C}, \mathbf{G}|\theta, \psi, \mathbf{Z})|P(\mathbf{O}, \mathbf{X}, \mathbf{C}, \mathbf{G})|\theta', \mathbf{Z}\big]\\ &= \sum_{lri} \log P(O_{lri}|X_{lri}, e, b)P(\mathbf{X}|\theta')\\ &+ \sum_{lri} \log P(C_{lri}|c_r)P(\mathbf{C}|\theta')\\ &+ \sum_{lri} \log P(X_{lri}|C_{lri}, \psi_l, G_l)P(\mathbf{C}, \mathbf{G}|\theta')\\ &+ \sum_{l} \log P(G_l|Z_l, \tau_k, F_k)P(\mathbf{G}|\theta')\end{aligned}$$

The probability terms are given in the previous section. What is missing is the expectation of the latent variables after each iteration. We calculate these recursively, using "forward"-probabilities to calculate the probabilities of states given the parameters, and the "backwards"-probabilities to calculate the probability of the observation given the latent states.

#### Forward Probabilities

We can make use of the recursive structure of the problem to calculate the marginal probability of $X_{lrj}$ by summing over the possible states in other latent variables. The probabilities $P(G_l|Z_l)$ and $P(C_{rli}|c_r)$ can be calculated without knowing the previous states.

**Read probabilities** For the read probabilities, we condition on $C$ and $G$:

$$
\begin{aligned}
Pr(X_{lrj}|C_{lrj}=0) =& \sum_g Pr(X_{lrj}|C_{lrj}=0, G_l=g)Pr(G_l=g|Z_l)\\
P(X_{lrj}|Z_l,\psi_l) =& Pr(X_{lrj}|C_{lrj}=0)Pr(C_{lrj}=0|c_r)Pr(G_l|Z_l)\\
&+ P(X_{lrj}|C_{lrj}=1)Pr(C_{lrj}=1|c_r)
\end{aligned}
$$

#### Backward Probabilities

The backward probabilities have form $Pr(\mathbf{O}|\cdot)$, where the $\cdot$ represents a latent variable. The probabilities $Pr(\mathbf{O}|\mathbf{X})$ can be calculated directly (see section Error Model). Given those, we can calculate the further probabilities for the other latent variables $\mathbf{C}$ and $\mathbf{G}$:

$$
\begin{aligned}
P(O_l|C_l) &= \prod_{rj} P(O_{lrj}|C_l)\\
&= \prod_{rj} P(O_{lrj}|X_{lrj})P(X_{lrj}|C_l)
\end{aligned}
$$

$$
\begin{aligned}
P(O_{lrj}|G_l) &= P(O_{lrj}, C_{lrj}=0) + P(O_{lrj}, C_{lrj}=1, G_l)\\
&= P(O_{lrj}|C_{rj}=0)Pr(C_{rj}=0) + \sum_x P(O_{lrj}|X_{lrj}=x)Pr(X_{lrj}=x|G_l, C_{rj}=1)
\end{aligned}
$$

#### Posteriors

Using the forward and backward probabilities, we can calculate the posterior expectations for the latent variable in each E-step.

**Posterior Genotypes** The probability that genotype $G_l$ is 0, 1, 2 is

$$
\begin{aligned}
P(G_l=g|\mathbf{O}) &= \frac{P(\mathbf{O}|G_l=g)P(G_l=g|Z_l)}{P(G_l=g|Z_l)} && (4)\\
&\propto P(G_l|Z_l) \times \prod_{rj} P(O_{lrj}|G_l) && (5)
\end{aligned}
$$

We ignore the denominator by calculating the probabilities for all three states and normalizing them to one, and all observation not associated with locus $l$ are the same for all genotypes, i.e. cancel out.

**Posterior Reads** The probability that read $X_{lrj}$ carries a derived allele

$$P(X_{lrj}|\mathbf{O}) \propto P(X_{lrj}|\mathbf{C},\mathbf{G})P(O_{lrj}|X_{lrj})$$

**Posterior Contamination** Calculate the posterior probability that read $rlj$ is contamination

$$P(C_{rlj}|\mathbf{O}) \propto P(O_{rlj}|C_{rli})c_r$$

#### M-step

Given the posterior expectations calculated in the E-step, we update the parameters for the next iteration in the $M$-step:

**Estimating $e$ and $b$**

To simplify notation, let $n_{a,b,c} = \sum P(X_{lrj} = b|O_{lrj} = a, W_l = c)$.

$$\hat{e} = \frac{n_{1,0,0} + n_{1,1,1}}{n_{1,0,0} + n_{1,1,1} + n_{0,0,0} + n_{0,1,1}}$$
$$\hat{b} = \frac{n_{0,1,0} + n_{0,0,1}}{n_{0,1,0} + n_{0,0,1} + n_{1,1,0} + n_{1,0,1}}$$

For example, $n_{1,1,1} = Pr(X_{lrj} = 1|O_{lrj} = 1, W_l = 1)$, i.e. a case where the read has a derived allele ($X_{lrj} = 1$), we observe an alternative allele ($O_{lrj} = 1$)) and the locus is defined as that the alternative allele is the ancestral allele ($W_l = 1$). This configuration occurs if there is an error away from the reference, which is why this term appears in the estimation of the new $e$.

**Estimating $c_k$**

$$c_r = \sum_{li} P(C_{lri}|O_{lrj})$$

i.e. we average over the posterior contamination estimates from all reads in the read group

**Estimating $\tau_k$ and $F_k$**

Done numerically by optimizing

$$\begin{aligned} Q(\tau_k, F_k|\tau_k', F_k') &= \sum_l I[Z_l = k] \log P(G_l|Z_l, \tau_k, F_k)P(\mathbf{G}|\tau_k', F_k') \\ &= \sum_l I[Z_l = k] \sum_{g=0}^{2} \log P(G_l = g|Z_l = k, \tau_k, F_k)P(G = g|\tau_k', F_k') \end{aligned}$$

where $I$ is an indicator function, $P(G = g|\tau_k', F_k')$ are the estimates from the previous iteration and $P(G_l = g|Z_l = k, \tau_k, F_k)$ are given by eq 3.

### SI1.3 - Calculating statistics

#### f-statistics

We can calculate *some* $F$-statistics using the $\tau$ estimates of `admixslug`. F2-, f3- and f4-statistics are outputted by `admixslug` when the `--output-fstats` flag is set. We use the estimates based on pairwise difference:

$$F_2(X,Y) = 2\pi_{xy} - \pi_{xx} - \pi_{yy} \tag{6}$$
$$F_3(X;Y,Z) = \pi_{xy} + \pi_{xz} - \pi_{yz} - \pi_{xx} \tag{7}$$
$$F_4(X,Y;Z,W) = \pi_{xz} + \pi_{yw} - \pi_{xw} - \pi_{yz}. \tag{8}$$

Assume we have $L$ loci, and population $a_{il}, d_{il}$ are the ancestral/derived counts in population $i$ at locus $l$, respectively, such that $n_{il} = a_{il} + d_{il}$. Then

$$\pi_{ij} = \begin{cases} \frac{1}{L}\sum_l \frac{a_{il}d_{jl} + d_{il}a_{jl}}{n_{il}n_{jl}}, & \text{if } i \neq j \\ \frac{1}{L}\sum_l \frac{a_{il}d_{il}}{n_{il}(n_{il}-1)}, & \text{if } i = j \end{cases} \tag{9}$$

using the conditonal SFS from `admixslug`, we can write equivalently

$$\pi_{ij} = \begin{cases} \sum_k s_k \frac{a_{ik}d_{jk} + d_{ik}a_{jk}}{n_{ik}n_{jk}}, & \text{if } i \neq j \\ \sum_k s_k \frac{a_{ik}d_{ik}}{n_{ik}(n_{ik}-1)}, & \text{if } i = j \end{cases}, \tag{10}$$

where $a_{ik}, d_{ik}, n_{ik}$ are now the counts of ancestral/derived/total alleles in population $i$ and SFS-category $k$, and $s_k$ is the (estimated) proportion of SNPs of this category.

These equations can directly be used to calculate $\pi$ within and between pairs of reference populations. To calculate $\pi$ between a reference population and the target individual, we use the estimator

$$\pi_{is} = \sum_k \frac{s_k}{n_{ik}} \left[ \tau_k a_{ik} + (1 - \tau_k) d_{ik} \right], \tag{11}$$

since $\tau_k$ is the expected proportion of SNPs carrying a derived allele in SFS-category $k$.

One caveat is that we cannot calculate $\pi ss$, the heterozygosity in the target individual. Also, for references without heterozygosity (e.g. the chimp-outgroup, or pseudo-haploid reference individuals), $\pi_{ii}$ cannot be calculated. By convention, we set these to zero. Thus, $F$-statistics involving these heterozygosities ($F_2$-statistics, and $F_3$-statistics using these individuals as samples) will be overestimated by a constant.

###### Standard error estimations for `f`-statistics

Similar to other methods, to account for linkage disequilibrium, admixslug uses block jackknives for standard error estimation. The size of the block that will be removed is controlled by the parameter `--jk-resamples`. This parameter accepts the number of blocks the genome will be divided into. If the value provided for this parameter is 10 for example, at each step 10% of the genome is removed. Let $f_{-i}$ denote the parameter estimate with the $i$-th block removed, $g$ denote the number of blocks and $\mu_{jk} = 1/g \sum_i f_{-i}$ be the jackknife mean. We estimate the jackknife variance as

$$\sigma^2_{jk} = \frac{g-1}{g} \sum_{i=1}^{g} (\mu_{jk} - f_{-i})^2 \tag{12}$$

#### $F(A|B)$ statistics

The $F(A|B)$ statistics where $B$ is a single diploid genome are the proportion of derived alleles at sites where $B$ is heterozygous. This corresponds exactly to the $\tau_k$ for categories where $k$ is heterozygous.

Thus,

$$F(A|B) = \frac{\sum_k s_k \tau_k}{\sum_k s_k}, \tag{13}$$

where the sum is only over entries where $B$ is heterozygous.

### SI2: Testing *admixslug*

#### SI2.1 - Model testing on simulations

In this section we evaluate the performance of *admixslug* on simulated data. We compared different parameters: coverage, proportion of contamination, proportion of deaminated reads being contaminants. The code used for running the simulations detailed below are publicly available on GitHub (https://github.com/LeonardoIasi/admixslug_simulations).

##### Simulated demography and scenarios

We assessed the performance using coalescent simulations from which we generate input data for *admixslug*. For this we used *msprime* (version 1.2.0)[5] to generate a demography of four populations. Chimpanzee (PAN), modern humans with sub-Saharan-African ancestry (AFR), non-African modern humans (non-AFR), Denisovans (DEN) and four Neandertal populations. The four Neandertal populations represent the Vindija 33.19 Neandertal lineage (Vi33.19 or VIN)[6], the Altai Neandertal lineage (D5 or ALT)[7], the Chagyrskaya 8 Neandertal lineage (Chag8 or CHA)[8] and the Mezmaiskaya 2 lineage (MEZ2)[9]. The chimpanzee lineage split from all others 6 million years ago (mya) and modern humans and the archaic humans split 600 thousand years ago (kya). The Denisovans split from the Neandertals 400 kya[7]. The modern humans split 70 kya with a subsequent 2,500 years long bottleneck in non-African modern humans, reducing their $N_e$ temporarily to 500. The Vindija lineage is chosen to be the 'main' lineage from which all others split as follows: Altai-Vindija 137 kya[6], Chagyrskaya-Vindija 100 kya and Mezmaiskaya 2-Vindija 61 kya[8]. We simulated a one generation long asymmetric gene flow 47 kya from the Vindija lineage into non-African modern humans contributing 3% of the genome. We sampled one diploid chimpanzee genome, 100 Africans, 50 non-Africans, one Denisovan sampled 70 kya[10], one Vindija Neandertal sampled 50 kya[6], one Mezmaiskaya 2 Neandertal sampled 45 kya[8], one Altai Neandertal sampled 125 kya[7] and one Chagyrskaya Neandertal sampled 80 kya[8]. We sampled two target individuals, one from the Altai lineage at 70 kya and one from the Mezmaiskaya 2 lineage 40 kya. Effective population sizes ($N_e$) were set to 10,000 for chimpanzees, Africans and non-Africans (except for the time of the bottleneck) and 2,000 for all archaic humans (**Figure SI2.1**).

We simulated diploid samples with 20 chromosomes per individual of 50 Mb length with a recombination rate of $10^{-8}$ per base pair per generation and a per base pair per generation mutation rate of $10^{-8}$, with a generation of 25 years.
We mimicked the ascertainment scheme used for the capturing of the data introduced in Skov et al., 2022[11]. In this case we used the sampled Vindija, Altai and Chagyrskaya Neandertals together with the

sampled Denisovan (all archaic humans: ARC) and the Africans to ascertain the data in the following way:

- AFR SNPs with derived allele frequency > 10%
- SNPs varying among the four ARC
- fixed differences between ARC and AFR
- sites fixed derived in ARC, while not fixed derived in AFR

For the target individuals we either used the true genotypes of all positions or ascertained positions directly to calculate *f*-statistics (ground truth), or we simulated sequencing data. For each SNP, the number of reads was drawn from a Poisson distribution with mean equal to the specified coverage. The total coverage was partitioned into endogenous and contaminant components according to the contamination proportion, such that endogenous reads were sampled as $Poisson(cov \cdot (1-cont))$ and contaminant reads as $Poisson(cov \cdot cont)$.

Given these read counts, alleles were sampled using a Binomial distribution. We sampled endogenous reads using the true derived allele frequency at the site in the target individual as the probability of sampling the derived allele. For contaminant reads, the derived allele probability was given by the allele frequency in the contaminant population. The observed allele counts were obtained by summing endogenous and contaminant derived and ancestral reads.

All reads have a minimum length of 30 bp. Contaminant reads were simulated given a gamma distribution with shape parameter 8 and scale parameter 7 resulting in an average read length of 86 bp (minimum read length + shape * scale) and varying amounts of being deaminated, whereas endogenous reads were simulated with shape parameter = 2 and scale parameter = 10 resulting in an average read length of 50 bp with a 40% chance of being deaminated.

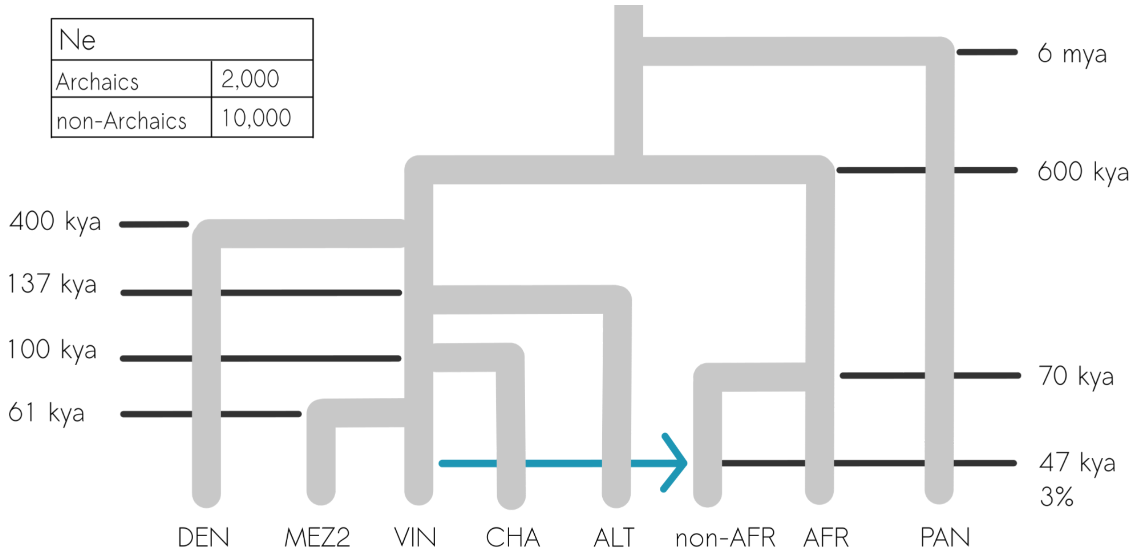

**Figure SI2.1:** Schematic representation of the simulated demography. Blue arrows represent the geneflow from Neandertals to the ancestors of modern humans outside of Africa ~47,000 years ago. Dark gray lines point out the dates of the split time and geneflow events. The table on the left corner presents the simulated effective population sizes for archaic and non-archaic human populations. The blue arrow represents gene-flow from a Vindija-like Neandertal population to all non-African modern humans.

#### Calculation of *f*-statistics

We used the following command to calculate *f-statistics* using *admixslug*:

```
admixslug --infile input_file --ref ref_file -o output_name \
--states VIN CHA ALT DEN --cont-id AFR –ancestral PAN \
--ll-tol 1e-2 –p-tol 1e-2 \
--max-iter 250 --filter-pos 80 –filter-ancestral \
–jk-resamples 10 –output-jk-sfs –output-fstats –no-snp –no-pars
```

For the ground truth we used the true simulated genotypes, for comparison we also computed *f-statistics* by simply randomly sampling one read per position from the *admixslug* input file mimicking random read sampling. We either did a random read sampling from all sites or only deaminated. The unbiased *f-statistic* was calculated as:

$$f_3 = \frac{1}{N}\Sigma\left[(p_{Ci} - p_{Ai}) \times (p_{Ci} - p_{Bi}) - p_{Ci}(1 - p_{Ci})\right]$$

$$f_4 = \frac{1}{N}\Sigma(p_{Ai} - p_{Bi}) \times (p_{Ci} - p_{Di})$$

$$F(A|B) = \frac{\Sigma p_{Ai} 1[p_{Bi}=0.5]}{\Sigma 1[p_{Bi}=0.5]}$$

We assume all individuals are diploid, the sums are over all SNPs $i$, $p_{xi}$ is the allele frequency of individuals $x$ at position $i$ and $N$ corresponds to the number of sites where all individuals have allele frequency information. To calculate the F(A|B) statistic we use an indicator function $I[]$ that is one when the condition holds (i.e. the site is heterozygous), and zero otherwise.

For the comparison to ADMIXTOOLS[3] we converted our input reference file for *admixslug*, which contains the high-coverage archaic genomes, and our target genome, into EIGENSTRAT format. For our low-coverage target genomes which we simulated, we used random read sampling per position. We either did random read sampling from all observed reads or subset to deaminated reads only. From this we created the EIGENSTRAT files. We calculated $f_4$-statistics using the ADMIXTOOLS wrapper *admixr*[12].

#### Testing the influence of the ascertainment

First, using our simulation framework, we assess potential biases introduced by the capture process, which is ascertained to specific genomic sites. We mimic this ascertainment scheme in our simulations and compare results obtained from the ascertained sites to those based on all polymorphic sites. For this comparison, we use the true simulated genotypes.

We compute $f_3$*(Chimpanzee; X, target)* and $f_4$*(Neandertal D5, B, target, Chimpanzee)* for the two target Neandertals. All scenarios show differences between estimates based on ascertained sites and those based on all sites (**Figure S12.2**).

We therefore conclude that estimates obtained under different ascertainment schemes are not directly comparable. Consequently, all subsequent analyses of simulated data are evaluated against the true values under the same ascertainment scheme.

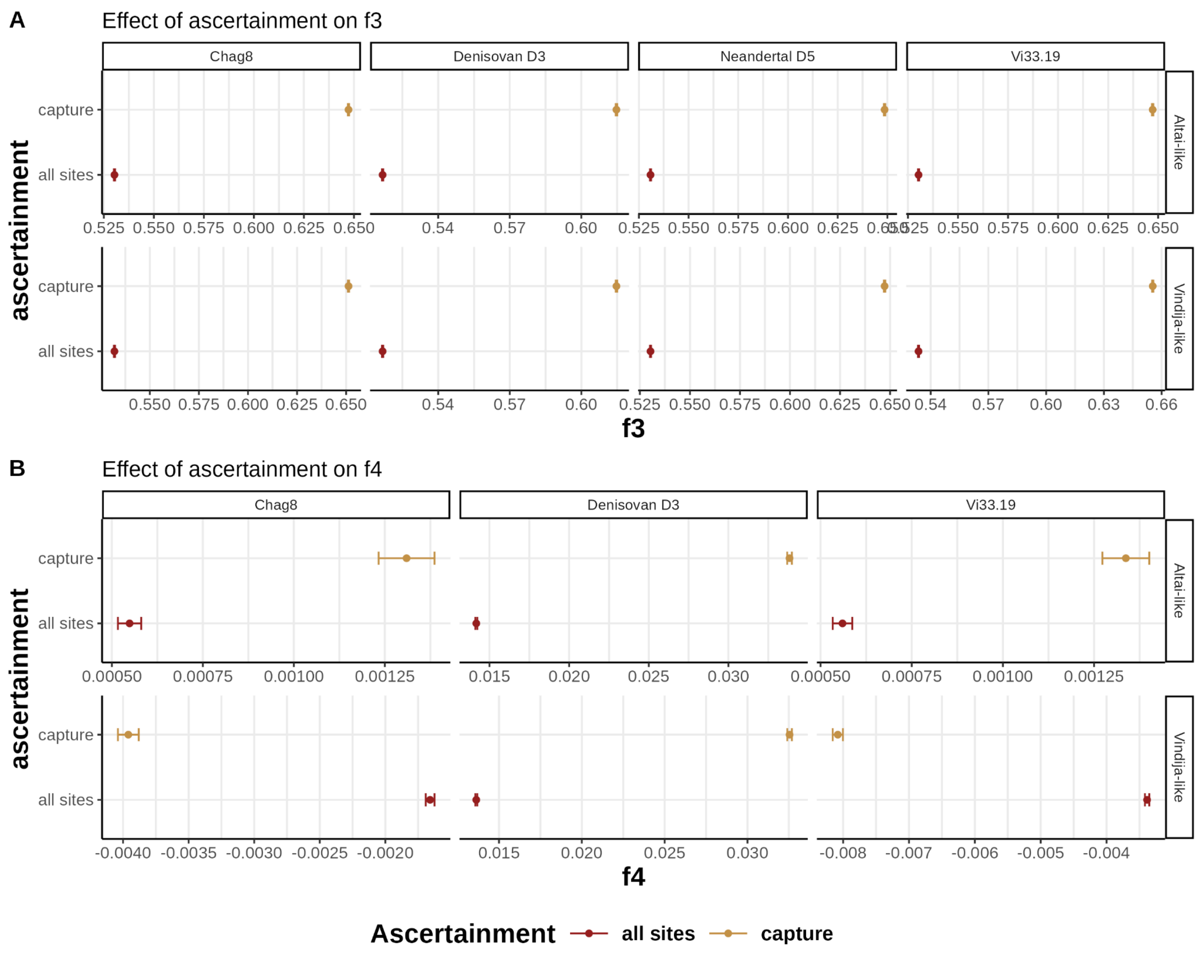


**Figure SI2.2:** Comparison between all sites and ascertainment mimicking the capture probe set from Skov et al., 2022[11]) for two simulated Neandertals across different *f-statistics*. *F-statistics* are directly estimated from simulated genotypes. Point indicates the mean estimate across 50 replicates and the error bars give one standard deviation.

#### Testing the estimation of contamination

**No contamination simulated in the deaminated sequences**

In the first simulations we have varying amounts of coverages for the same target genomes and varying amounts of contamination. All deaminated reads are coming from the target genome. **Figure SI2.3** shows that *admixslug* can accurately estimate the overall amount of contamination.

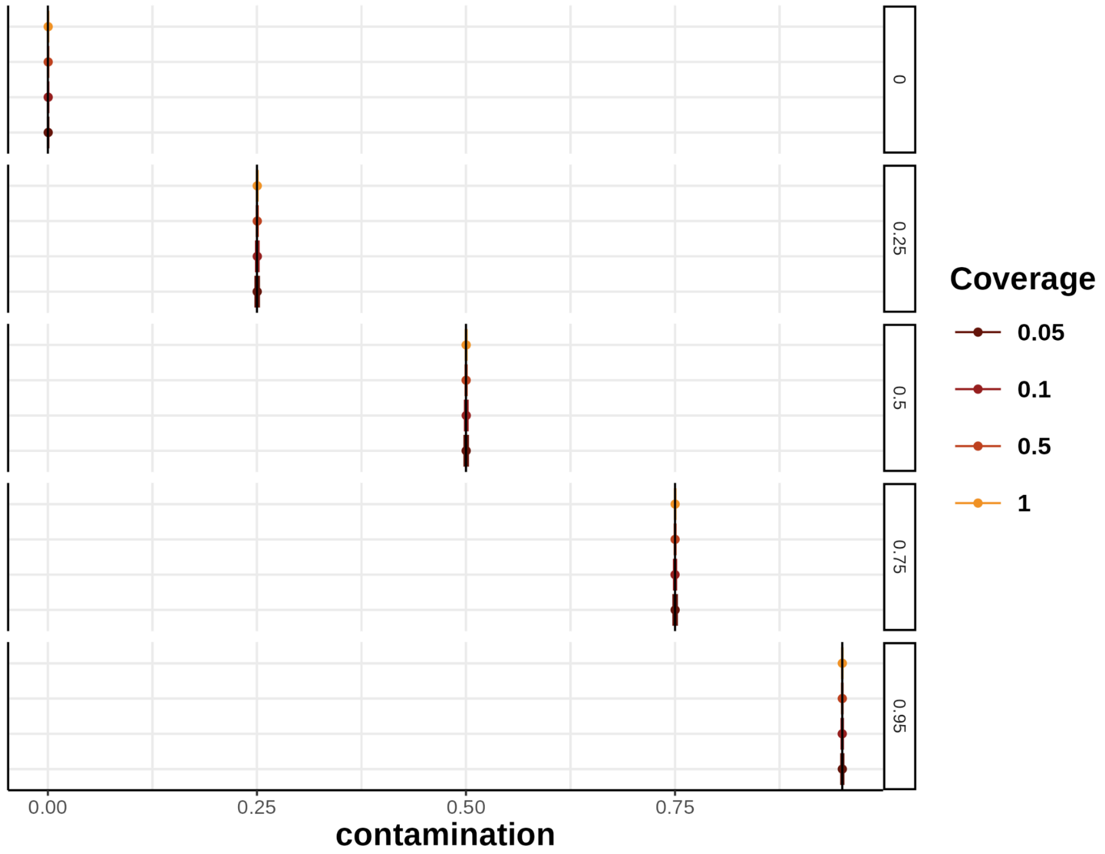


**Figure SI2.3:** Contamination estimated by *admixslug* across all simulations with contamination only in non-deaminated reads. Colors correspond to coverage while rows correspond to amount of contamination as a proportion of all reads. Black line indicates the true contamination proportion. The dots indicate mean across all *admixslug* estimates for this scenario and one standard deviation.

**Simulated contamination in the deaminated sequences**

It is a common practice to subsample the data set to reads that show signals of deamination when there is high levels of overall present-day human DNA contamination. This implicitly assumes that reads coming from a contaminant source do not contain deaminations, an assumption that is sometimes violated for low-coverage Neandertals[13–15]. Therefore, we test the ability of *admixslug* to estimate contamination when contaminant reads are also deaminated. We demonstrate in **Figure SI2.4** that *admixslug* can correctly estimate the proportion of contamination even in the most extreme scenario where more than 30% of all reads with terminal deaminations are contaminated.

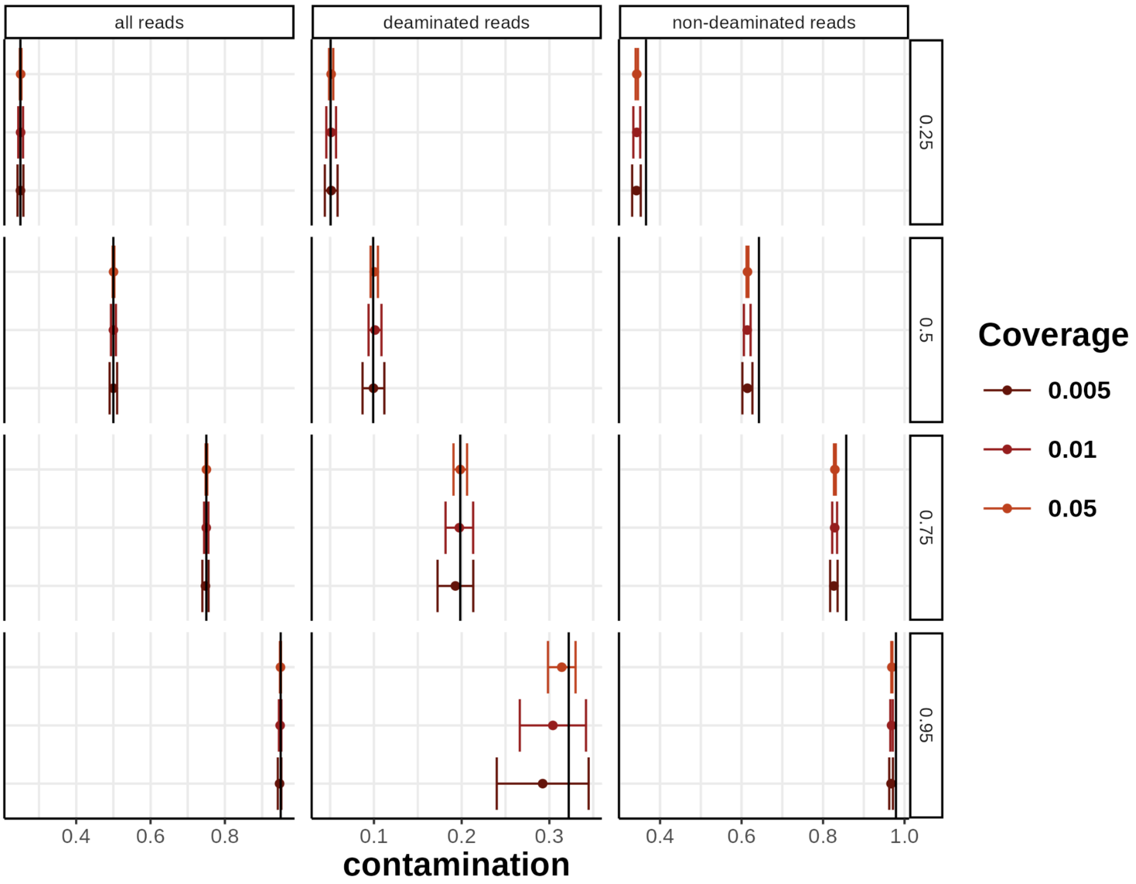


**Figure SI2.4:** Contamination estimated by *admixslug* across all simulations with contamination in both deaminated and non-deaminated reads. Colors correspond to average depth of coverage while rows correspond to the overall amount of contamination as a proportion of all reads. Columns indicate the estimate for all reads, the estimate for deaminated and non-deaminated reads. The black line indicates the true contamination proportion for all reads, deaminated reads and non-deaminated reads. The dots indicate mean across all *admixslug* estimates for this scenario and one standard deviation.

#### Validation of the *f*-statistics on simulations

We examine the performance of *admixslug* to estimate f-statistics in scenarios where the target Neandertal (either Vindija (Vi33.19) or Altai-like (D5)) has different coverages and different amounts of contamination. We compare the estimates to estimates calculated on the raw data either on all reads, or when reads are subsampled to those with deamination. We start by testing the simpler scenario where contamination is only in sequences without deamination. After showing that *admixslug* can estimate *f-statistics* accurately, we test the more complicated scenario where contamination is also found in the deaminated reads.

##### $f_3$*-statistics*

We obtain unbiased $f_3$ estimates using *admixslug* across all tested overages and contamination levels. The estimates show a smaller variation compared to the estimates using deaminated reads only (**Figure SI2.6**).

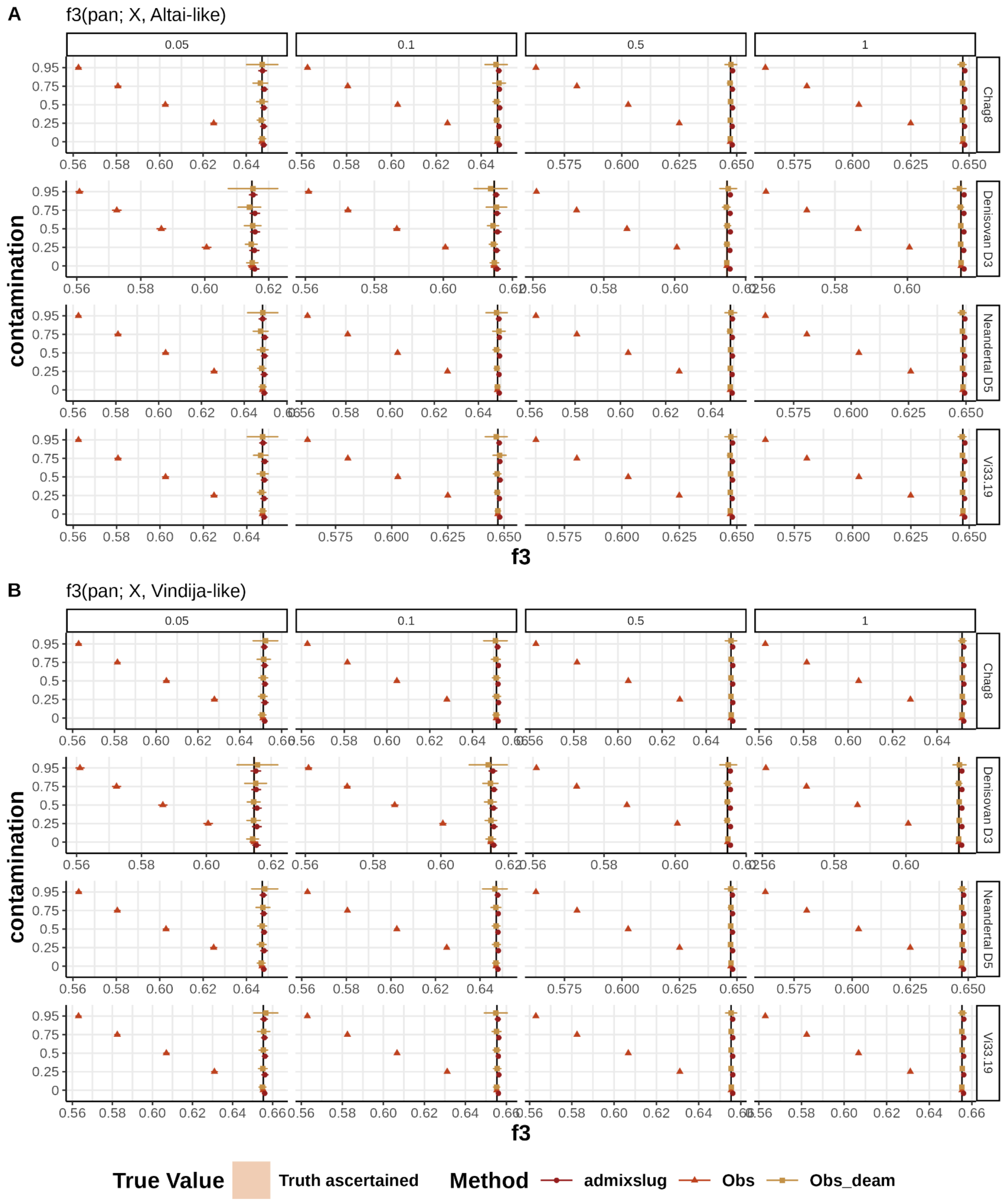


**Figure SI2.6:** Comparison of $f_3$ estimates obtained using *admixslug*, all reads (Obs), and deaminated reads only (Obs_deam) across different contamination levels. The points represent the mean estimate across 50 replicates, and error bars show ±1 standard deviation. The rows indicate the individual used as population X in the $f_3$-*statistic*, and columns indicate the average depth of coverage of the target genome. The colours denote the estimation method. The black line shows the mean true value, and the shaded area represents ±1 standard deviation of the true values across replicates.

##### $f_4$-*statistics*

We observe a similar behaviour for $f_4$-*statistics* (**Figure SI2.7**).

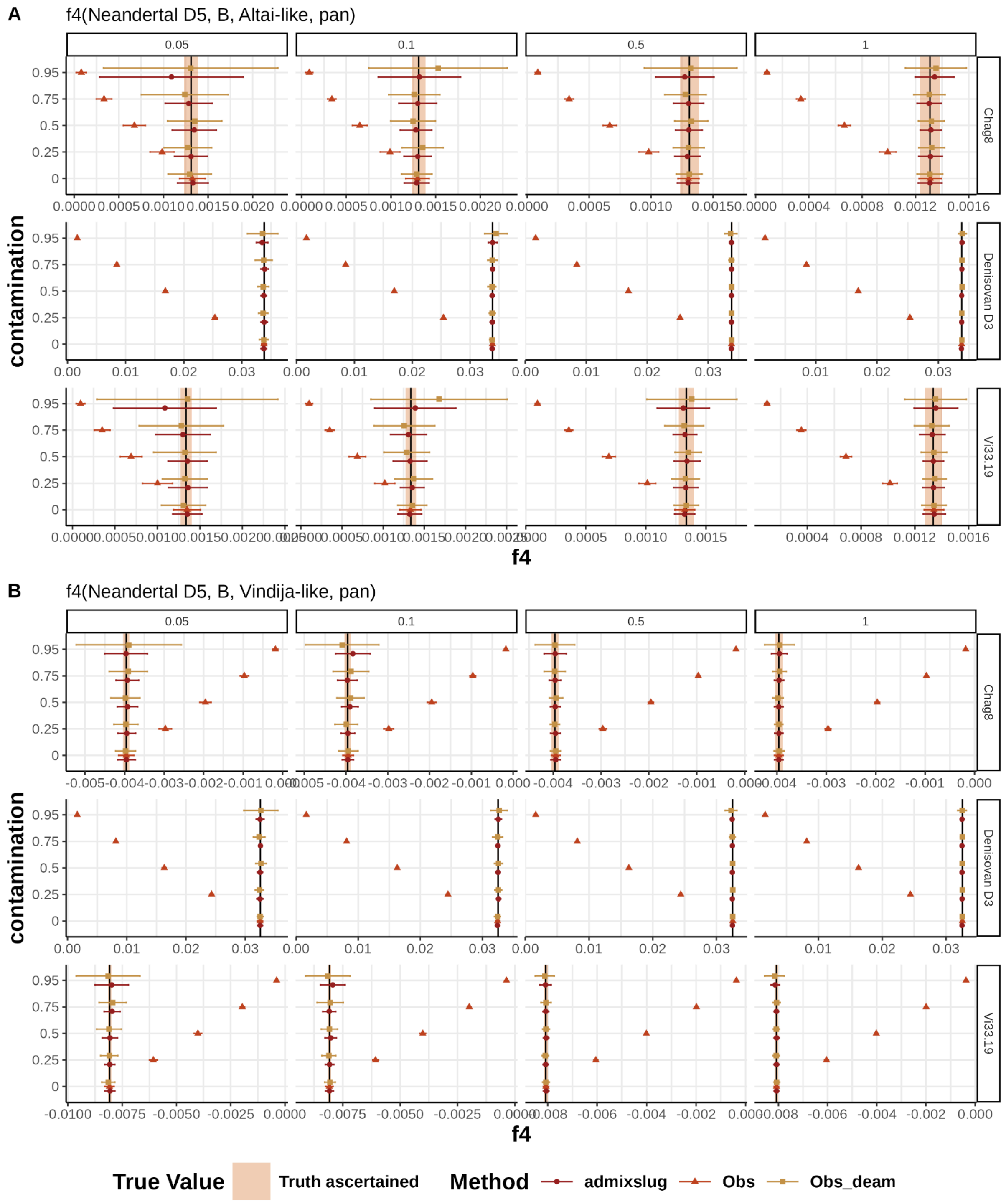


**Figure SI2.7:** Comparison of $f_4$ estimates obtained using *admixslug*, all reads (Obs), and deaminated reads only (Obs_deam) across different contamination levels. Points represent the mean estimate across 50 replicates, and error bars show ±1 standard deviation. The rows indicate the individual used as population B in the *$f_4$-statistic*, and columns indicate the depth of coverage of the target genome. The colours denote the estimation method. The black line shows the mean true value, and the shaded area represents ±1 standard deviation of the true values across replicates.

Next, we consider a more challenging scenario in which contaminated reads also exhibit deamination. We simulate contamination levels of 0%, 25%, 50%, 75%, and 95%, with corresponding proportions of deaminated reads originating from contaminants of 0%, 5%, 10%, 20%, and 30%, respectively. We further focus on very low coverages of 0.05×, 0.01×, and 0.0005×.

At low contamination levels, subsetting to deaminated reads results in only a small bias in the estimates, although with reduced power compared to *admixslug* (**Figure SI2.8**) However, as shown in **Figure SI2.8**, this approach fails to produce unbiased estimates at higher contamination levels. In contrast, *admixslug* reliably provides unbiased $f_4$ estimates across all contamination levels at a minimum coverage of 0.05×, and at lower coverages remains unbiased up to contamination levels of 75%.

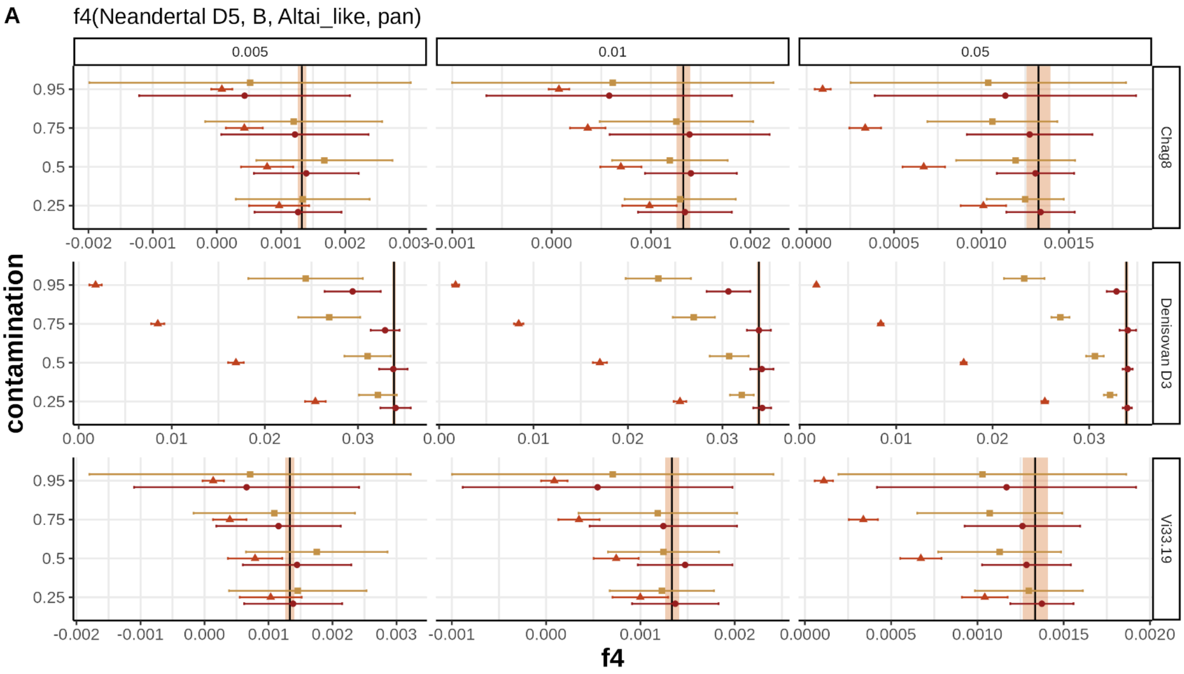

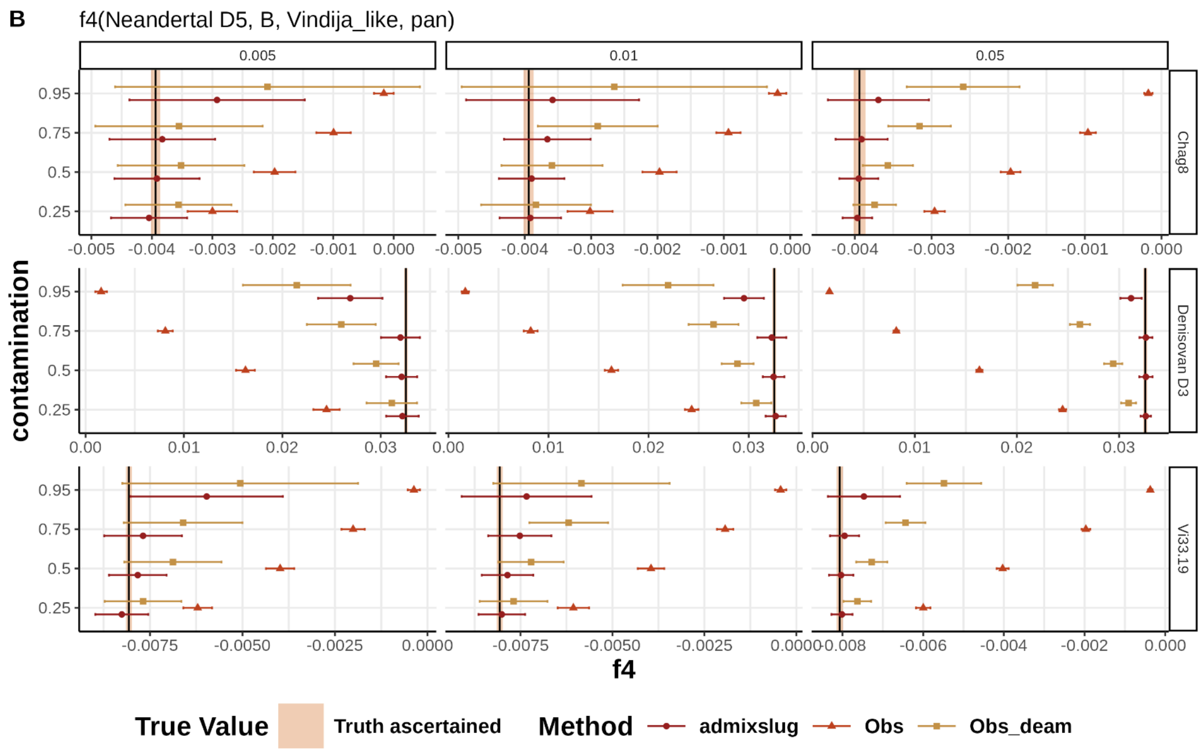


**Figure SI2.8:** Comparison of $f_4$ estimates when there is deamination in contaminated reads. Estimates are obtained using *admixslug*, all reads (Obs), and deaminated reads only (Obs_deam) across different contamination levels. The points represent the mean estimate across 50 replicates, and error bars show ±1 standard deviation. The rows indicate the individual used as population B in the *$f_4$-statistic*, and columns indicate the average depth of coverage of the target genome. The colours denote the estimation method. The black line shows the mean true value, and the shaded area represents ±1 standard deviation of the true values across replicates.

**Validation of the F(A|B) statistics on simulations**

Here we test if downsampling and adding contamination introduces bias to these estimates, using simulations. We again consider two scenarios of contamination I) where all the contamination is found in non-deaminated sequences, and II) some of the contaminant sequences have deamination. For scenario I we compared how F(A|B) values change when we have 0%, 50%, 75% and 95% contamination for varying coverage depth (**Figure SI2.9**). Our results indicate that even for the lowest coverage and highest contamination test cases, *admixslug* can estimate the correct F(A|B) statistics.

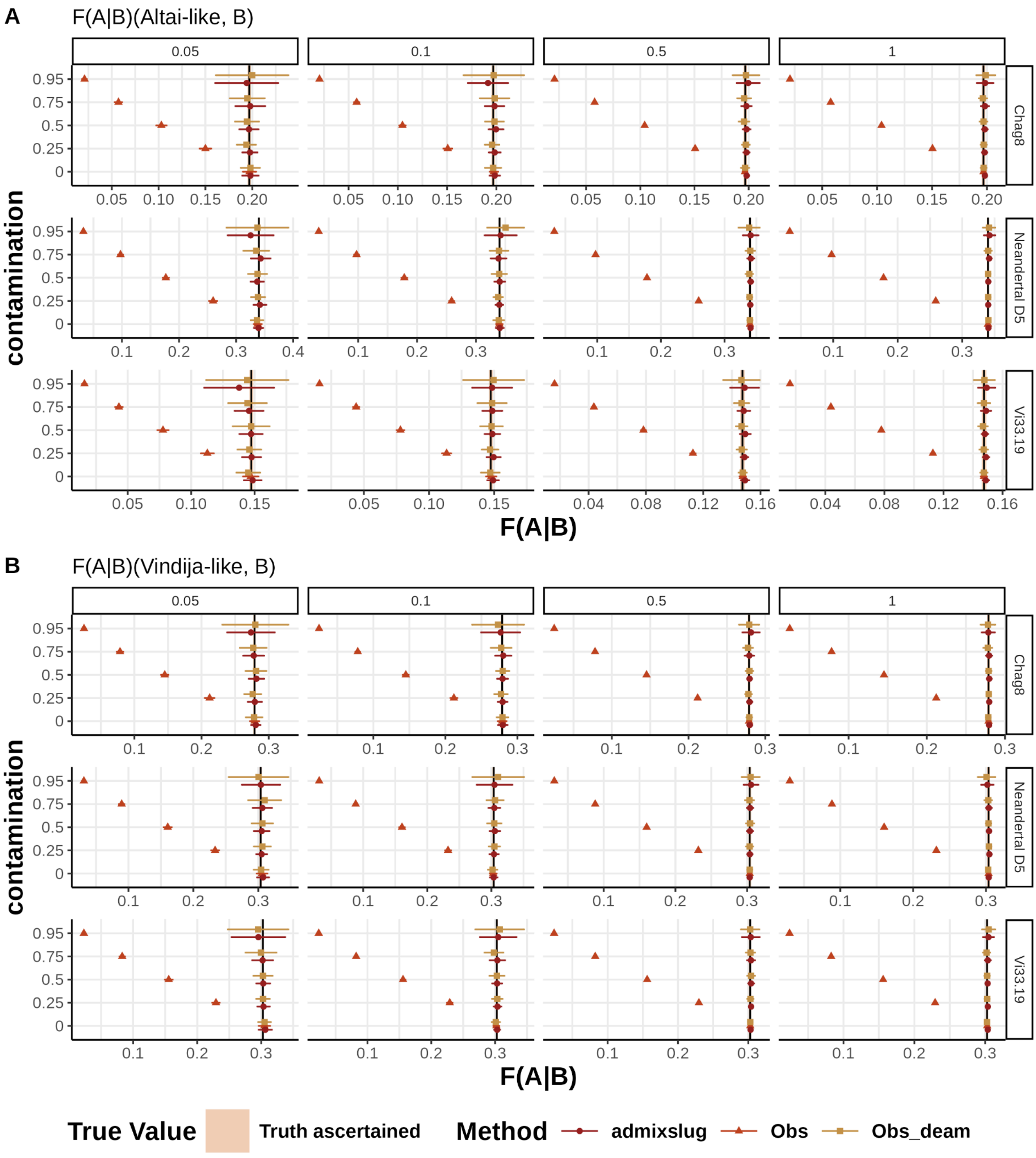


**Figure SI2.9:** Comparison of F(A|B) statistics calculated from simulations. Estimates are obtained using *admixslug*, all reads (Obs), and deaminated reads only (Obs_deam) across different contamination levels. The points represent the mean estimate across 50 replicates, and error bars show ±1 standard deviation. The rows indicate the individual used as population B in the F(A|B) statistic, and columns indicate the coverage of the target individual. The colours denote the estimation method. The black line shows the mean true value, and the shaded area represents ±1 standard deviation of the true values across replicates.

To test the more challenging scenario II where we have high levels of contamination also in the deaminated sequences, we explored *admixslug*s performance on even lower depth of coverages, 0.01x and 0.005x. We show in **Figure SI2.10** that *admixslug* provides mostly unbiased estimates except for high contamination scenarios with extremely low coverage. More specifically, *admixslug* estimates of

F(A|B) overlap with the true estimate 69 out of 72 of the simulated scenarios. The 3 cases where *admixslug* estimate does not overlap the truth are when simulated contamination is 95% and overall average depth of coverage is 0.005x, meaning only 0.00025x coverage for the endogenous DNA component.

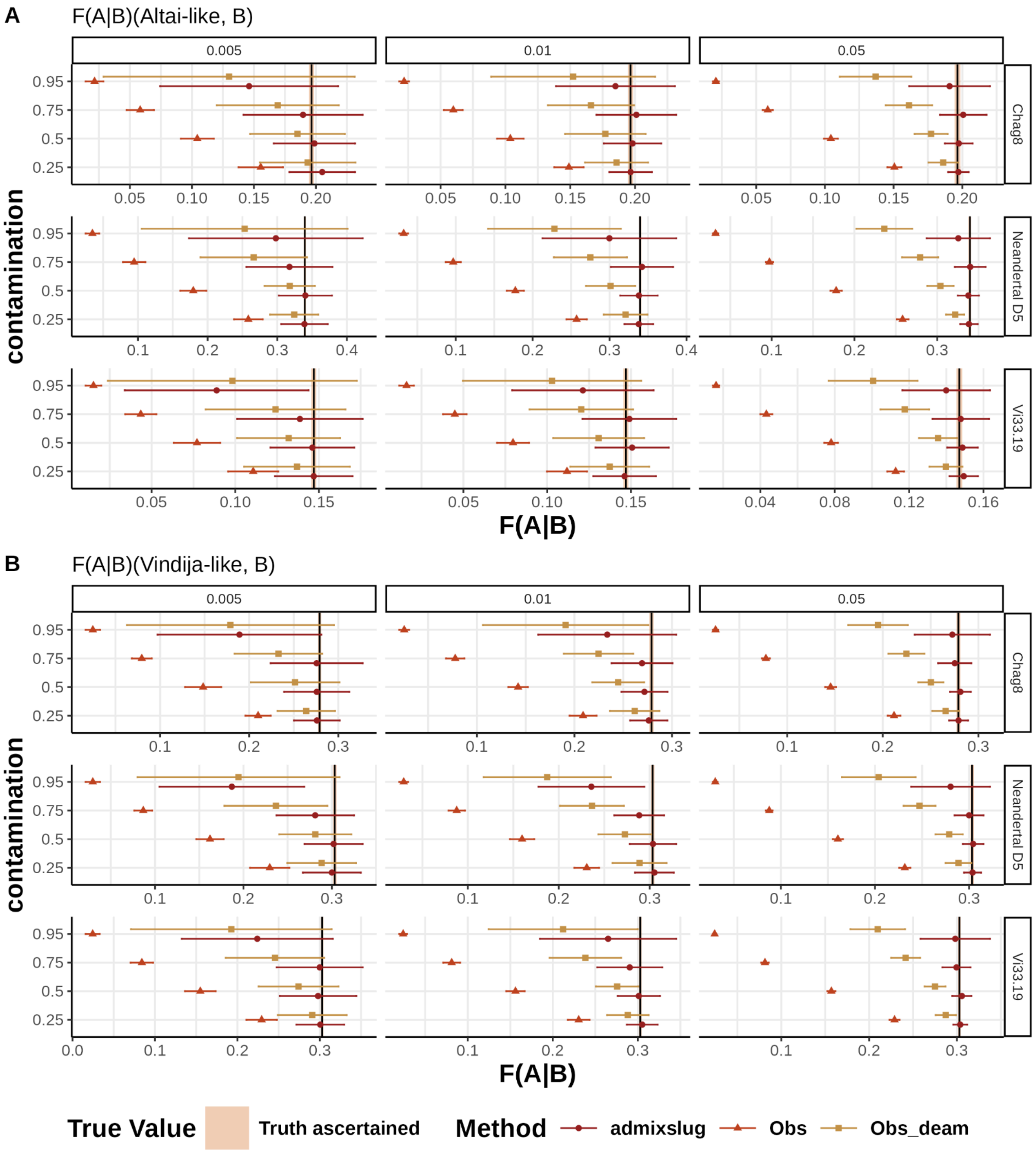


**Figure SI2.10:** Comparison of F(A|B) statistics calculated from simulations when there is deamination in contaminated reads. Estimates are obtained using *admixslug*, all reads (Obs), and deaminated reads only (Obs_deam) across different contamination levels. The points represent the mean estimate across 50 replicates, and error bars show ±1 standard deviation. The rows indicate the individual used as population B in the F(A|B) statistic, and columns indicate the coverage of the target genome. The colours denote the estimation method. The black line shows the mean true value, and the shaded area represents ±1 standard deviation of the true values across replicates.

#### Single-estimate performance and confidence interval calibration

Next, we compare single estimates of the *$f_4$-statistics* between *admixslug* and the widely used ADMIXTOOLS software[3]. We compared the two methods in the scenario with deamination in the contaminated reads and let ADMIXTOOLS run either on all data or only on the deaminated reads. While the confidence intervals of *admixslug* overlap the true value except in two instances, ADMIXTOOLS estimates on all data almost never overlap with the truth (**Figure SI2.11**). Also the estimate on deaminated reads only show increasingly biased estimates with higher contamination level or result in uninformative estimates with CI spanning from -1 to 1.

We observe that ADMIXTOOLS generally produces tighter confidence intervals when applied to deaminated reads only, compared to *admixslug*. Although *admixslug* incorporates uncertainty in genotype calls, which is expected to result in wider confidence intervals, its estimates are based on substantially more data.

We therefore calibrated the CI using all simulation scenarios (i.e. across all different contamination and coverage parameters). While *$f_3$(Chimpanzee; X, target)* and F(A|B) estimates include the true value 94% and 92% of the time using ±2 standard errors, respectively, *$f_4$(Neandertal D5, B, target, Chimpanzee)* estimates include the true value 99% of the time. We find that using ±1.5 standard errors already achieves approximately 95% coverage in $f_4$ estimates from *admixslug*. Because of this, we use 1.5 standard errors when plotting 95% CI in our analyses using real data and we point it out.

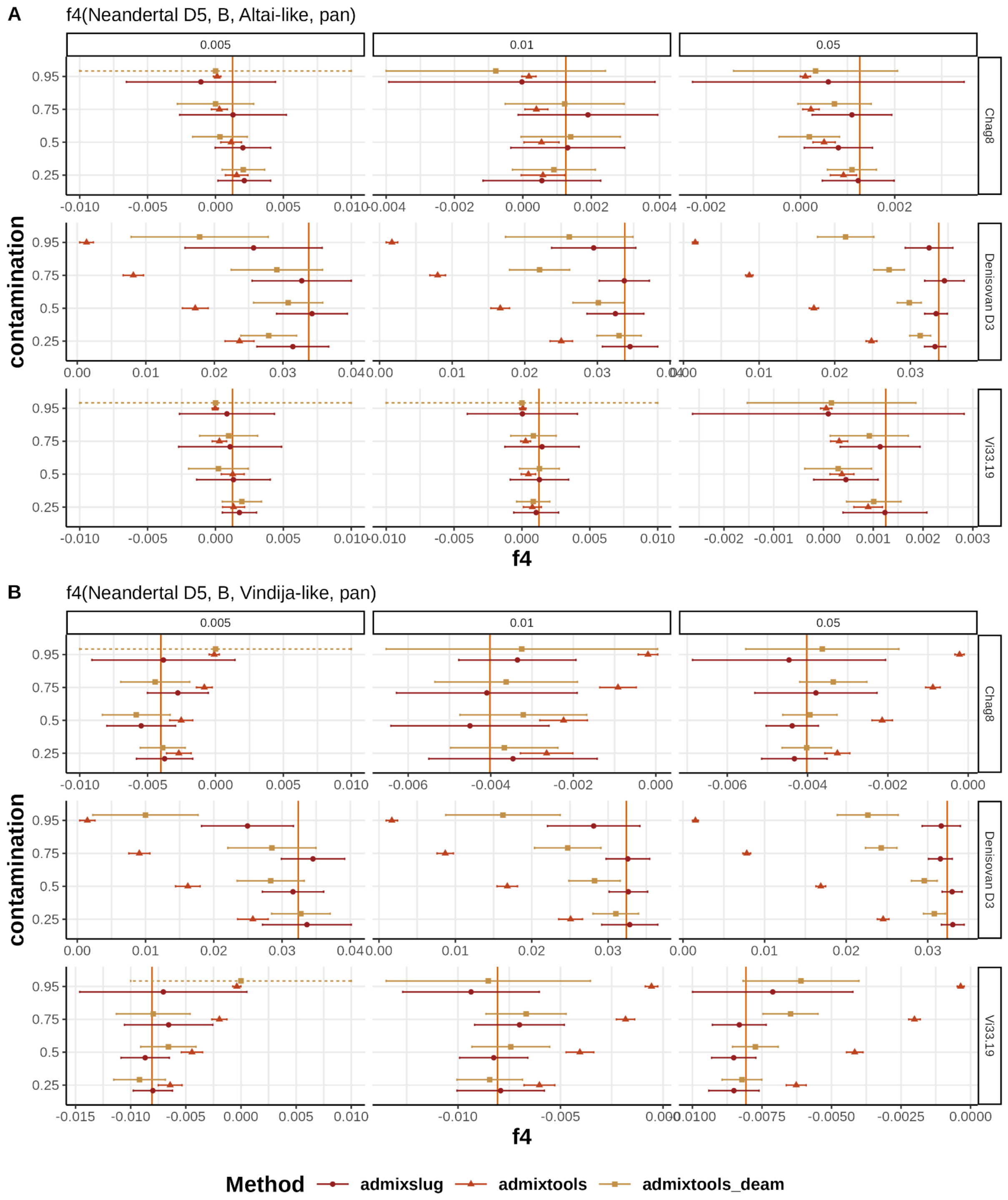


**Figure SI2.11:** Comparison of single $f_4$-*statistics* calculated from simulations when there is deamination in contaminated reads. Estimates are obtained using *admixslug*, ADMIXTOOLS on all reads (admixtools), and ADMIXTOOLS on deaminated reads only (admixtools_deam) across different contamination levels. The points represent the mean estimate, and error bars show ±2 standard errors. The dotted error bars indicate intervals that extend beyond the displayed axis limits. The rows indicate the individual used as population B in the $f_4$-*statistic*, and columns indicate the average depth of coverage of the target genome. The colors denote the estimation method. The orange vertical line shows the true value.

#### SI2.2 - Model testing on real data

To assess the performance of *admixslug* on real data, we artificially contaminated the low-coverage genome of Mezmaiskaya 2, a ~45,000 years old Neandertal[9], by mixing in present-day DNA sequences in varying proportions. Mezmaiskaya 2 was chosen for having minimal contamination (~0.87%) and a 1.7x coverage genome obtained through shotgun sequencing of four libraries. We filtered the Mezmaiskaya 2 genome for positions that overlap with a nuclear capture array containing 643,472 transversions across the genome[11], removed PCR duplicates and filtered for read length 35 bp and mapping quality 25.

We sampled reads from present-day human DNA fragments modified to mimic the contamination typically observed in real data[16], as the source of contamination mixed with the uncontaminated Neandertal genome. Before sampling reads proportionally to the number of reads in the Mezmaiskaya 2 genome, we filtered our contamination source file the same way, with minimum length of 35 base pairs and mapping quality of 25, on the target sites. Even though terminal C-to-T substitutions are not a characteristic of the present-day human contamination, some sequences carry these by chance. In our contaminant data, approximately 2.5% of the sequences carried this pattern, causing minimal contamination in the deaminated sequences.

By proportionally introducing reads from the contaminant, we created different versions of the Mezmaiskaya 2 genome with 10%, 25%, 50%, 75%, 95% contamination, as well as a 100% contamination dataset composed exclusively of contaminant sequences - to test *admixslug's* behaviour in the absence or endogenous DNA. Additionally, to test the effect of decreasing coverage depth on the accuracy of contamination estimates, we downsampled these contaminated versions and the unmodified Mezmaiskaya 2 genome to 1x, 0.5x, 0.1x, 0.05x, 0.01x and 0.005x average coverage, as we did in our tests on the simulations (see **SI2.1**).

As an example, we create the scenario with 95% present-day human contamination scenario, by randomly sampling reads from the contaminant source file and uncontaminated Mezmaiskaya2 file in a 19:1 ratio, respectively, and merging them. The resulting file is then downsampled to the desired coverage. Thus, in the most extreme case with 95% added contamination and 0.005x coverage, the endogenous Mezmaiskaya 2 component represented only 0.00025x coverage, or around 160 endogenous and 3,000 contaminant reads overlapping variable sites.

In this analysis our aim was to investigate the performance of *admixslug* in estimating overall contamination, and compare these with estimates from AuthentiCT[17]. AuthentiCT is a published method to infer present-day human DNA contamination in low coverage ancient genomes, but assumes that the deaminated sequences (sequences with terminal C to T substitutions) are uncontaminated. For comparability, here we did not test the effect of contamination in the deaminated sequences, although *admixslug* is capable of assessing this type of contamination in addition to the overall contamination (see **Section SI2.1**). Lastly, before running *admixslug* we modified the input file to hide the library IDs, to blind it to the readgroup.

We ran *admixslug* with the command :

```
admixslug --in input_file \
       --states VIN CHA ALT DEN \
       --ref ref_file \
       --ancestral PAN \
       --cont-id EUR \
       -o output_name --ptol 0.001 --ll-tol 0.01 \
       --max-iter 100 --filter-ancestral \
       --jk-resamples 500 --output-jk-sfs --output-fstats
```

#### Contamination

For the *admixslug* analyses, we split up sequences by sequencing library, length, and whether they start with a terminal deamination or not, here using the default setting of the first three and last three bases of the molecule. *Admixslug* first splits molecules by library and deamination status, and then bins the remaining reads so that each bin has 50,000 reads on average.

To make the results comparable with AuthentiCT[17], we estimated a weighted average of these to calculate the overall average present-day human DNA contamination in each test genome. Taking the previously reported amount of contamination of 0.87% as the truth[9], we calculated the difference estimated-true to assess the accuracy of contamination estimates. We observe that both methods perform very well, with *admixslug* providing estimates that overlap with the correct value more often (**Figure SI2.12**). AuthentiCT slightly overestimates the present-day human DNA contamination for the range of 10%-75% for most downsampled coverages, but by at most 3%. (**Figure SI2.12**). Overall, we find that *admixslug* is highly accurate and precise for estimating present-day human DNA contamination, even for very low coverage genomes.

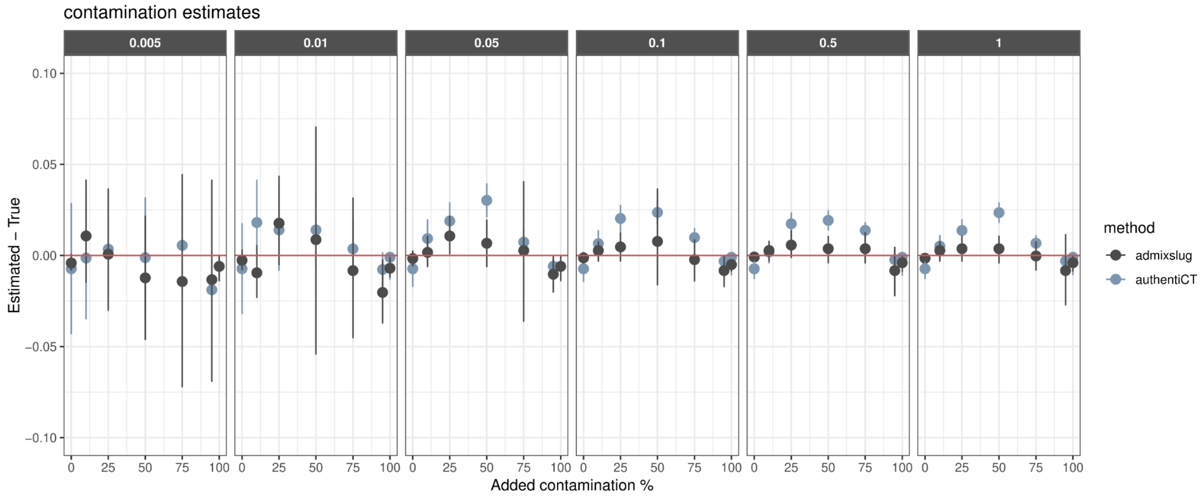


**Figure SI2.12:** Average present-day human contamination estimates from *admixslug* and AuthentiCT[17]. The difference between the estimated value and the true amount of contamination is plotted on the y-axis as proportions, while the added contamination amount is shown on the x-axis. Column grids represent the average depth of coverage, varying between 0.005x and 1x. In the scenario with 100% contamination, AuthentiCT could not produce an output and hence is missing from the plot for average coverages of 0.05x and 0.005x. The error bars on the estimates correspond to two standard errors of the mean for both *admixslug* and AuthentiCT. The red line is at zero, where the estimate is equal to the true value.

#### *f*-statistics

When the "--output-fstats" parameter is used, *admixslug* outputs various $f_2$-, $f_3$- and $f_4$-statistics with all combinations of genomes indicated with "--states" and "--ancestral" parameters, and the target. Using the original Mezmaiskaya 2 genome (1.7x and ~0% contamination) as the true value, we present the $f_3$- and $f_4$-statistics for downsampled and contaminated versions of Mezmaiskaya 2. As Mezmaiskaya 2 is a Late Neandertal, she is genetically most similar to the ~45,000 years old Neandertal from the Vindija Cave in Croatia[6]. Keeping this in mind, below we present results from various $f_3$- and $f_4$-statistics, and compare values estimated for the contaminated and downsampled Mezmaiskaya 2 genomes to the uncontaminated original for these statistics.

##### $f_3$-statistics

We calculated outgroup $f_3$-statistics using a chimpanzee genome (PAN) as the outgroup. The statistics $f_3$*(A, Mezmaiskaya 2; PAN)* measures the genetic similarity between genome "A" and Mezmaiskaya 2. In our analyses, the high-coverage archaics Vindija 33.19 (VIN)[6], Chagyrskaya 8 (CHA)[8], Altai (Denisova 5) (ALT)[7] and Denisova 3 (DEN)[10]) are in position "A". We confirm the expectation that Mezmaiskaya 2, a late Neandertal, is genetically most similar to the Vindija Neandertal, and we obtain the highest $f_3$ values among all genomes in "A" for Vindija. Also, the $f_3$-values obtained when the

Denisovan genome is “A” is lower than for all three high-coverage Neandertal genomes. When we have 100% present-day human DNA contamination in our sample Mezmaiskaya 2 test genome, we obtain the lowest $f_3$-values (**Figure SI2.13**). This makes sense as the statistics becomes *$f_3$(A, Modern human, Chimpanzee)*, and modern humans are an outgroup to Neandertals and Denisovans.

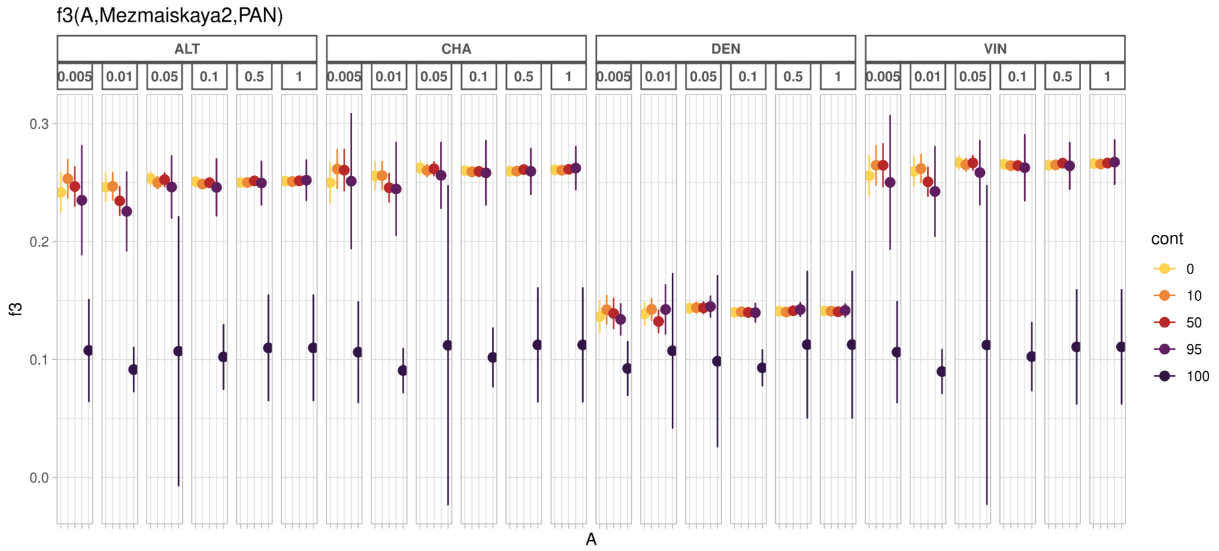


**Figure SI2.13:** $f_3$-outgroup statistics estimated by *admixslug* for different versions of contaminated and downsampled Mezmaiskaya 2 Neandertal genome. Column grids represent the high-coverage archaic genome used as “A” in the statistics *$f_3$(Mezmaiskaya 2, A, Chimpanzee)*. Different colours stand for the added percentage of present-day human DNA contamination in the samples, and the further column grids indicate the average depth of coverage of the different Mezmaiskaya 2 test genomes. Error bars are three standard errors of the mean.

For all high-coverage archaic genomes used as genome “A”, all downsampled and contaminated versions of the Mezmaiskaya 2 genome, *admixslug* is able to estimate *$f_3$-statistics* with high accuracy. We note that for the most extreme scenario of a 0.005x average coverage with 95% contamination confidence intervals of the $f_3$-value overlaps with the true value of the statistics (estimated from the original data with no contamination and 1x average depth of coverage, yellow points on the rightern most subpanels in **Figure SI2.13**) under the ascertainment used throughout this paper.

##### *$f_4$*-statistics

We calculated the $f_4$-statistics that measures if the Mezmaiskaya 2 genome shows more genetic affinity to the high-coverage Vindija genome or to another high-coverage archaic genome by computing *$f_4$(Vindija, B, Mezmaiskaya 2, Chimpanzee)* using *admixslug*. This statistics should be positive because Mezmaiskaya 2 shows higher similarity to Vindija compared to the individual in position “B”, as was previously shown[9]. We observe that the value of this $f_4$-statistics is always positive when genome in position B is the Altai Neandertal (D5) or the Denisovan (D3), except in cases where we introduce 100% contamination. When the Chagyrskaya Neandertal is in position B, statistics calculated here start to become not significant for coverages lower than 0.05x. This is because the Vindija and Chagyrskaya Neandertals are genetically more similar to each other, when compared to the Altai Neandertal and the Denisovan, and we start losing power for very low coverages.

Our results show that even with 95% contamination *admixslug* can estimate the correct $f_4$-value, but the larger error will lead to false negatives (error bars of three standard errors overlap with 0) when the coverage is very low (e.g. $f_4$*(Vindija, Altai, Mezmaiskaya 2, Chimpanzee)* when coverage is lower than 0.05x) (**Figure SI2.14**). For coverages as low as 0.005x, we can still find a statistically significant result for this statistics up to 50% contamination.

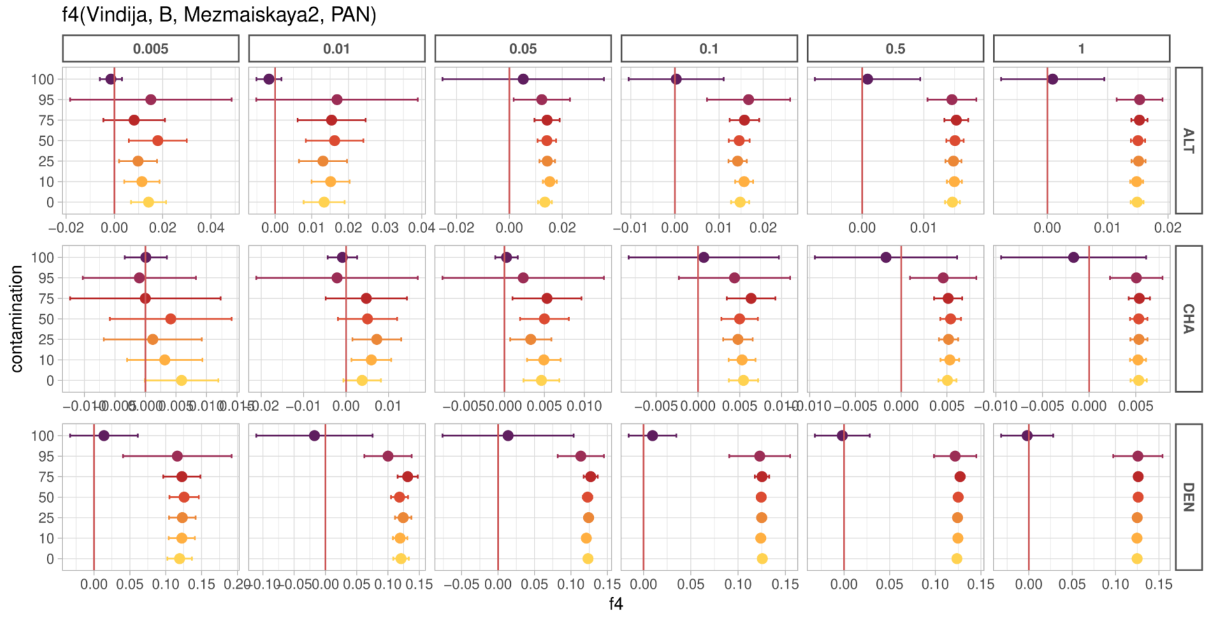


**Figure SI2.14:** $f_4$-statistics estimated by *admixslug* for different versions of contaminated and downsampled Mezmaiskaya 2 Neandertal genome. The row grids represent the high-coverage archaic genome used as "B" in the statistics $f_4$*(Vindija, B, Mezmaiskaya 2, Chimpanzee)*. Different colours stand for the added percentage of present-day human DNA contamination in the different versions of the Mezmaiskaya 2 genome, and the column grids indicate the average depth of coverage of these different versions. Error bars are two standard errors of the mean.

##### Comparison with ADMIXTOOLS

ADMIXTOOLS[3] is the most widely used software to calculate F-statistics. We compare the performance of ADMIXTOOLS and *admixslug* on the downsampled data set with and without added present-day human DNA contamination. Unlike *admixslug*, ADMIXTOOLS is naive to the present-day human contamination in the target samples and uses a single, random read per SNP instead of all the data. In ancient DNA studies, when present-day human DNA contamination is detected in the sample, usually the data is subsetted to deaminated sequences, with the implicit assumption that none of the deaminated sequences come from the contaminant. In our tests with real data, this is the case as the contaminant sequences we introduced were not deaminated. Hence, we expect ADMIXTOOLS to produce the correct $f_4$-statistics when the data is subsetted to the deaminated sequences. Due to not having enough data, ADMIXTOOLS did not run for the scenario where we had 95% contamination, 0.005x coverage and deaminated reads only.

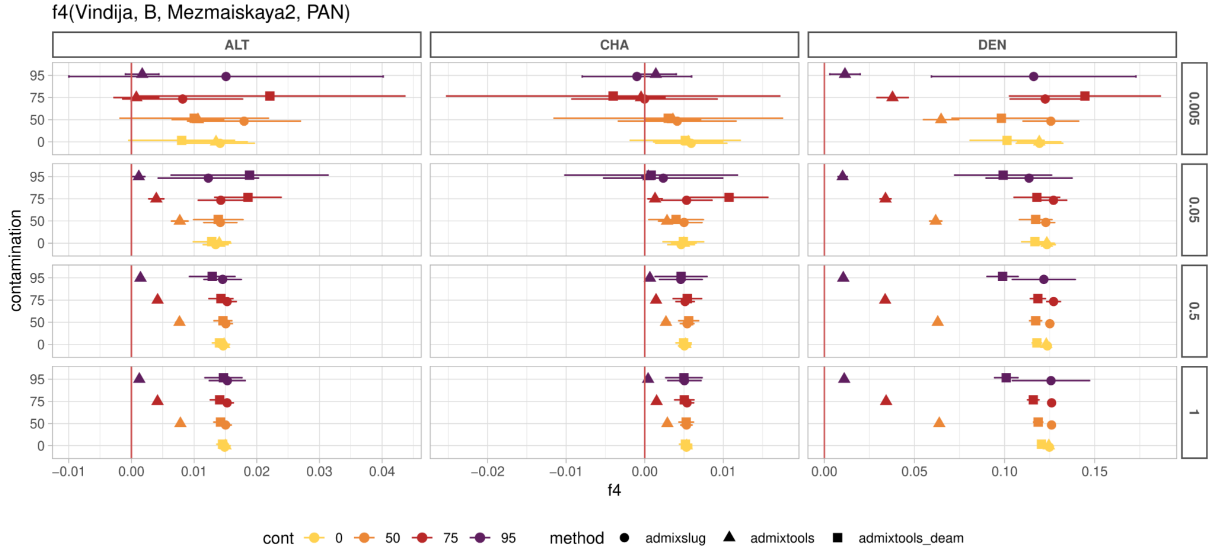


**Figure SI2.15:** $f_4$-statistics estimated by *admixslug* and ADMIXTOOLS for contaminated and downsampled Mezmaiskaya 2 genomes. The shapes of the points correspond to the method, and different colours stand for the added percentage of present-day human DNA contamination in the different versions of the Mezmaiskaya 2 genome. Column grids represent the high-coverage archaic genome used as "B" in the statistics $f_4$*(Vindija, B, Mezmaiskaya 2, Chimpanzee)*, and the row grids indicate the average depth of coverage of these different versions. Error bars represent the 95% CI (1.5 standard errors for *admixslug*, as recalibrated in **SI2.1**).

When all sequences are used in ADMIXTOOLS values of the $f_4$-statistics shift to 0 with increasing amounts of contamination. We note that ADMIXTOOLS estimates are highly biased with narrow confidence intervals. Depending on the amount of contamination, interpretation from $f_4$-statistics estimated can differ (**Figure SI2.15)**.

When only deaminated sequences are used, the shift is much weaker (since contamination rate in the retained data is much lower). In this case, the ADMIXTOOLS estimates sometimes overlap the correct value of the statistics and in other cases, such as for the statistic *f4(Vindija, Denisova, Mezmaiskaya 2, Chimpanzee)* the bias is strong enough to remain statistically significant **Figure SI2.15**.

In all test scenarios, *admixslug* estimates the correct value of the statistics and in most cases overlap with the ADMIXTOOLS estimates from the deaminated sequences. For low-coverage data, *admixslug* has larger confidence intervals than admixtools. This is because *admixslug* takes the uncertainty in the genotype into account, which random-read sampling ignores, resulting in overly confident estimates.

In cases with minimal present-day human contamination in the deaminated sequences, we find that *admixslug* and ADMIXTOOLS on the deaminated sequences only performs comparably for some cases but not all, depending on the genome in position B (see the lowest panel in **Figure SI2.15** where B is the Denisovan (D3)), likely due to traces of contamination in the deaminated sequences. Overall, our method is more reliable, and can run with very little data, where ADMIXTOOLS can not (e.g. 95% contamination, 0.005x subsetted to deaminated sequences. More importantly, here we test cases where there is no contamination in the deaminated sequences. Unlike *admixslug*, ADMIXTOOLS does not have a contamination model and produces biased estimates in the presence of deaminated present-day human contamination (**Figure SI2.11**).

#### Ascertainment used for capture

Genomic data from our focal individual, Teshik-Tash 1, was obtained using a capture array introduced in the Skov et al 2022[11] and described in **SI3**. Because of this, we subset all data to the sites on this array, even when genomes are sequenced through shotgun sequencing (e.g. for Mezmaiskaya 2, as well as all Neandertal genomes in the comparative set in **SI3**). For the simulations, we use an ascertainment scheme that mimics the capture array design (**SI2.1**).

Our tests on simulations indicated that the ascertainment changes the overall values of the *f*-statistics estimated, but not relative patterns. However, simulating an ascertainment perfectly is not possible. To see how subsetting genome-wide data to these sites influence the $f_3$- and $f_4$-statistics in real data, we compared the *admixslug* results obtained from low-coverage shotgun sequenced genomes of Mezmaiskaya1, Mezmaiskaya2 and Spy 94a **(I)** after filtering those for the capture site ascertainment, and using an *admixslug* reference panel specifically for those sites and **(II)** using the shotgun genomes without ascertainment, with an *admixslug* reference panel that includes all polymorphic sites in archaic humans.

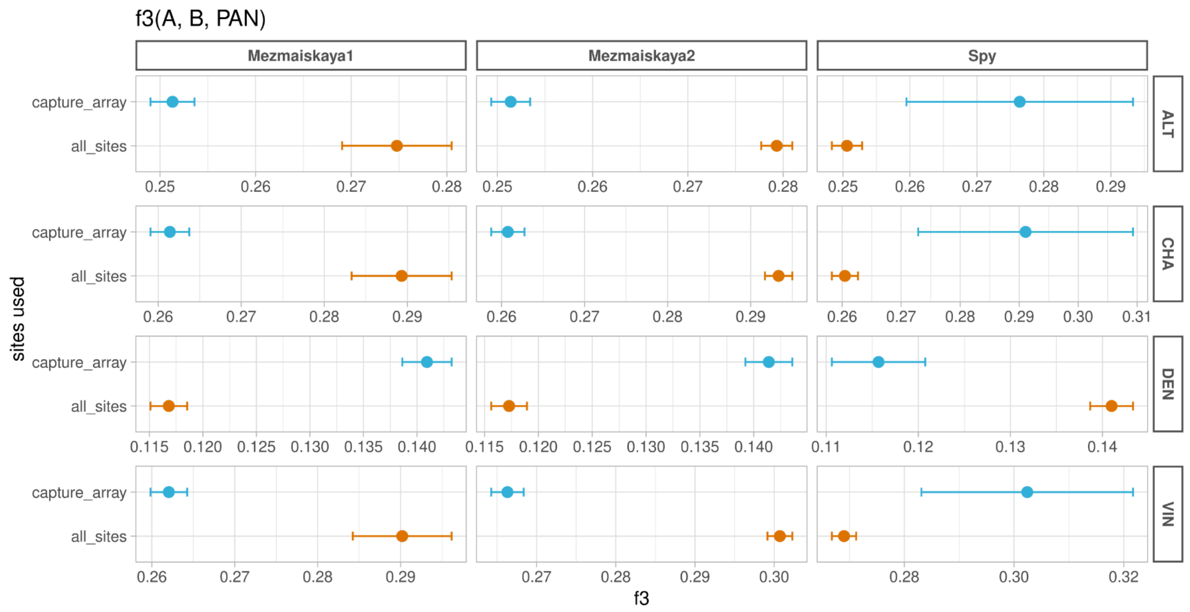


**Figure SI2.16:** The effect of ascertainment on the $f_3$-statistics for $f_3$(A, B, Chimpanzee) where the genome in A is shown on the row grids on the right y-axis, and genome in B is the low-coverage Neanderta genomes shown on the column grids on the top. The x-axis shows estimated values of the $f_3$-statistics. Blue shows the estimates obtained by using the capture array ascertainment that is used throughout the manuscript, while orange shows the estimates obtained by using all polymorphic sites in the archaic humans. Error bars represent 2 standard errors of the estimate.

Similar to what we found using simulations in **SI2.1**, here we show that estimates from both $f_3$- and $f_4$-statistics (**Figures SI2.16 and SI2.17**) are not comparable when different ascertainments are used. Although this is not an *admixslug* specific issue, it is something to be aware of when comparing results from differentially obtained data[3]. For comparable results, we suggest using the same reference file with the same ascertainment[18].

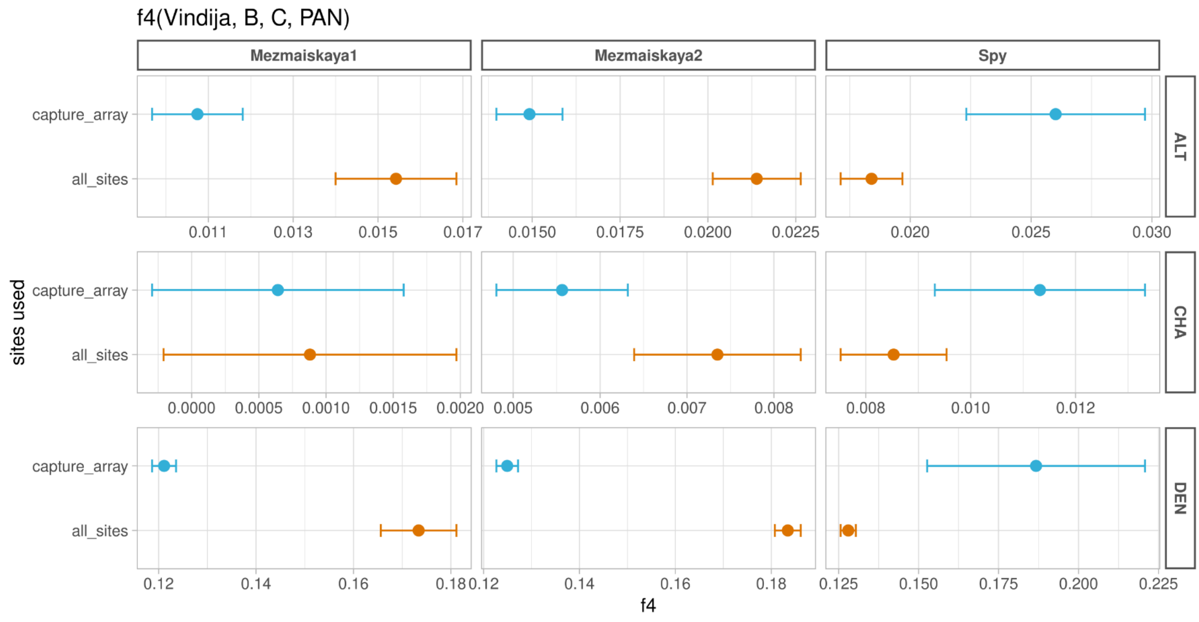

**Figure SI2.17:** The effect of ascertainment on the $f_4$-statistics estimates for $f_4$(Vindija, B, C, Chimpanzee). The column grids indicate the low-coverage Neandertal genome used in position C, and row grids are the high-coverage Neandertal genomes used in position B. Estimated values of the $f_4$-statistics are shown on the x-axis. Blue shows the estimates obtained by using the capture array ascertainment that is used throughout the manuscript, while orange shows the estimates obtained by using all polymorphic sites in the archaic humans. Error bars represent 2 standard errors of the estimate.

### SI3: Application on Teshik-Tash 1

#### Teshik-Tash Cave

One of the most famous Palaeolithic sites of Central Asia, Teshik-Tash Cave is situated in the Baisuntau Mountains, a southwestern extension of the Hissar mountain range, about 20 km north of the town of Baisun, Uzbekistan[19]. Here a deep gorge, known as Zautolosh cuts into the limestone massif, and the cave lies about 6 m above the bottom of the gorge. Access to the site is very hard, as the approximately 20 m wide gorge has almost vertical walls and is over 50 m deep. The cave opens to the northeast, with an about 20 m wide and 7 m high entrance; the depth is about 21 m (Movius, 1953, BVs observations). In 1938 and 1939, an expedition of the Marr Institute for the History of Material Culture of the USSR Academy of Sciences under the direction of A.P. Okladnikov and V.D. Zaporozhskaya excavated about 137 m2 of the site, removing all Palaeolithic sediments from the cave. They observed a succession of five thin cultural horizons separated by sterile layers of clay, silt and sand[20]. They interpreted the 1.3-1.5 m thick succession as several short-term visits interspersed with flooding of the cave[19]. In 2003, when researchers of the Institute for Archaeology, Siberian Branch Russian Academy of Sciences re-excavated the site, the gorge was completely dry during the summer, but it carries large amounts of water particularly in spring during snow melt. It can be assumed that at the time of occupation, the cave was closer to the bottom of the gorge, and thus could have been flooded periodically.

#### *The "Burial"*

The richest cultural horizon is the uppermost one, Cultural Layer I, and the child "burial" was discovered directly underneath this level. Three hearths with diameters ranging from 0.5 to 1 m were described in this level, one of them directly overlying the hominid remains. As the remains of the child were surrounded by five (or six) pairs of *Capra sibirica* horncores, Okladnikov assumed that the individual was buried in an elaborate rite[19]. According to his descriptions, the horns were originally arranged in a circle, with the child buried in the middle, parallel to the cave wall[20]. Later, the body was exhumed by carnivores, as shown by the gnaw marks on the femur. Gargett strongly criticized the burial hypothesis[21]. In his opinion, no clear signs of intentional burial are present, and carnivore activity is a more parsimonious explanation for the presence of the bones. One of the arguments against the burial hypothesis is the absence of a documented burial pit in any of the sections published by Okladnikov. The presence of a shallow pit was only inferred from the distribution of the bones. Gargett's strong opposition to the recognition of any kind of Middle Palaeolithic burials[21,22] has been criticized before (e.g. see CA* treatment of Gargett 1989, Hovers et al., 2000)[21,23], and as Gilman pointed out, using Gargett's criteria „…would sweep away the evidence for burials from virtually all pre-1960 excavations for periods prior to the Neolithic." (Gilman,1989, p.182)[24].

#### Teshik-Tash 1 child

##### Morphology

The cranium and mandible of Teshik-Tash 1 are quite well preserved[25], and a number of postcranial fragments were also discovered (Sinelnikov and Gremyatskiy, 1949)[26]. These include the atlas, an axis fragment, both clavicles, several rib fragments, the left humerus shaft, partial left femur and tibia, and both fibulae (fragmented). The first description of the human remains was by Aleš Hrdlička[27], who had the chance to study the remains in Moscow. He interpreted the remains as belonging to a normally developed Neandertal child 8 to 9 years of age. Rokhlin emphasizes the Neandertal-like characteristics of the skeleton, such as the robusticity of the long bones, the marked muscle insertions on the femur and humerus, and the massive browridges (Rokhlin, 1949, p.120)[28]. These traits are no longer seen as Neandertal traits, but as a general archaic morphology. Robust long bones with thick cortical bone are not only present in Middle Pleistocene *Homo*, but also in Early AMHs (e.g. the Jebel Irhoud 4 humerus[29], and the Qafzeh juveniles[30]).

Glantz and colleagues reassessed the affinities of the Teshik-Tash child using multinomial logistic regression and discriminant function analyses.[31] They found that metrically the mandible clustered with recent humans for both raw and size-standardized data, while the cranium showed Upper Palaeolithic affinities when size-standardized. In their opinion, the hypothesis that Teshik-Tash 1 is a typical Neandertal can not be further held up, but it is impossible to give a clear assessment of its affinities. A geometric morphometric analysis of the Teshik-Tash 1 frontal was performed by Gunz and Bulygina[32]. Using semilandmarks on surfaces, they not only compared juveniles, but they also predicted adult shapes for Teshik-Tash 1 using both Neandertal and modern human growth trajectories. The shape of the frontal bone of Teshik-Tash 1 is similar to Neandertal juveniles, and the adult morphologies predicted by both growth trajectories are similar to Neandertals, especially to La Chapelle aux Saints 1.

A specialized odontological analysis of the Teshik-Tash 1 identified three distinct morphological strata[33]. According to author, the dental system exhibits: (1) a pronounced suite of archaic characteristics, (2) a near-complete expression of Neandertal-derived traits, and (3) a number of evolutionary progressive features of the dental system (Khaldeeva, 2010, p. 123[33]). Multivariate analysis incorporating multiple anthropological character systems positions the specimen within the morphological range of Western European Neandertal variants[33].

The pursuit of additional taxonomic markers prompted researchers to conduct an analysis of frontal sinus morphology (size and shape) in different hominids. The results support the conclusion that neither masticatory load adaptation nor climatic factors determine frontal sinus morphology in them[34]. The analysis revealed that *H. neanderthalensis* exhibits no evidence of hyperpneumatization when compared to *H. sapiens*, *H. erectus* or other hominids. The analyzed *H. neanderthalensis* specimens demonstrate significantly reduced variability in sinus morphology (both size and shape) compared to other fossil hominin groups. Our investigation of the Teshik-Tash 1 specimen employed conventional X-ray and computed tomography (CT), which show complete absence of frontal sinus development (**Figure SI3.1**). This contrasts with modern human children of comparable age (~9 years), where frontal sinuses are typically present. This anatomical absence may reflect either developmental variation and/or taxonomic distinctiveness. Current interpretations remain constrained by insufficient data on sinus development in Neandertal juveniles, and lacking Pleistocene reference standards for this ontogenetic feature.

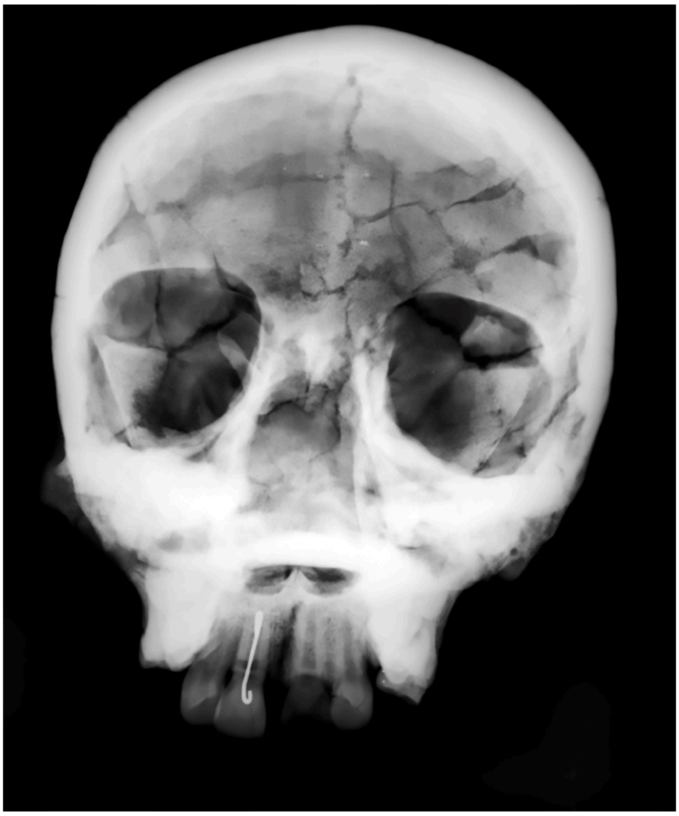

**Figure SI3.1:** The X-ray of the frontal part of the skull of Teshik-Tash 1.

##### Archaeology

The skeletal remains of Teshik-Tash 1 were found in association with Middle Palaeolithic lithic assemblages, but this industry was found to be distinct from the assemblages in other Neandertal sites such as the Chagyrskaya and Okladnikov Caves in the Altai Mountains, which are possibly contemporaneous with Teshik-Tash 1[35]. The assignment of the site to the Middle Palaeolithic was not changed by attempts of radiocarbon dating[32], which were not successful.

Recent archaeological evidence indicates that the Teshik-Tash industry is characterized by a combination of hierarchical and non-hierarchical core reduction strategies, the former including a distinct form of Levallois technology. Notably, triangular points – common elsewhere in the Middle Paleolithic of Southwest Asia – are exceptionally rare at this site. These findings establish the Teshik-Tash industry as a significant Neandertal cultural tradition in western Central Asia[35]. Another archaeological analysis confirms the predominance of centripetal reduction strategies in the Teshik-Tash lithic assemblage[36]. The study documents significant technological variability evident in both: (1) initial flake blank production, and (2) subsequent tool modification phases. Levallois reduction constitutes a minor component of the assemblage.

##### Genetics

Ancient DNA was previously obtained from the Teshik-Tash 1 individual, but was limited to the HVR1 region of the mitochondria[37]. Analyses of this placed Teshik-Tash 1 closer to earlier Neandertals such as Scladina and Altai (D5), rather than Late Neandertals in Europe or the possibly contemporaneous ones from Chagyrskaya and Okladnikov[37]. However, the mitochondrial DNA is only a single maternally inherited locus, and does not reflect the whole population history of an individual.

Teshik-Tash represents the southeastern-most range of the Neandertal distribution (**Figure SI3.2**), and falls close to the known geographical range of the Denisovans. Due to the absence of nuclear DNA, it has not been possible to investigate if Teshik-Tash 1 individual carried any genetic contribution from Denisovans.

In our study, we improved the quality of the mitochondrial genome of Teshik-Tash 1 as well as sequenced nuclear DNA from various specimens that were attributed to this individual. Due to extremely high levels of present-day human DNA contamination in the nuclear genome of this individual, the common methods for calculating *f*-statistics were not reliable. Because of this, we used *admixslug* to assess the population affinities of Teshik-Tash 1.

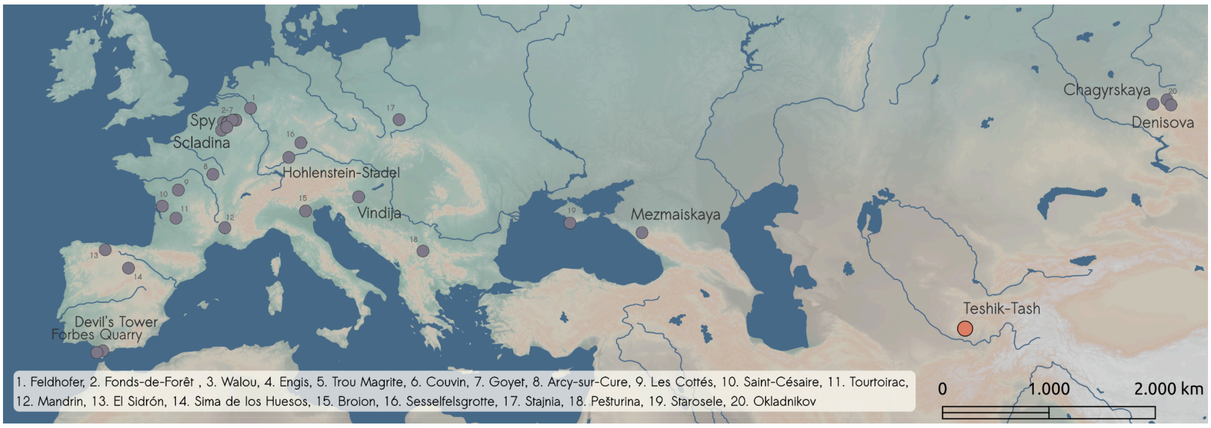


**Figure SI3.2:** Geographical distribution of the sites with Neandertals which yielded ancient DNA (mitochondrial and/or nuclear). The Teshik-Tash site is shown in pink and represents the southeastern-most extent of the Neandertal range to date. The base map was made with Natural Earth (www.naturalearthdata.com/).

#### SI3.1 - Data generation

We sampled two skeletal remains from the Teshik-Tash 1 individual, the petrous part of the left temporal bone (SP.A.3640) and a long bone fragment (SP.A.1047, SP.A.3029, SP.A.3030; different sample IDs were generated for different sampling spots on the same bone, see **Table SI3.1**). Sampling for ancient DNA analyses was carried out after the removal of a thin layer of surface material, using a sterile disposable dentistry drill. First, the petrous bone was sampled in Moscow in 2015, for a total of 32.0 mg of bone powder. Of this, an aliquot of 10.7mg was washed with a sodium phosphate buffer in order to remove contamination[38]. DNA was then extracted using a silica-based method optimized for the retrieval of short DNA molecules[39]. An aliquot of this extract was converted into a single-stranded DNA library[40], and barcoded with a pair of unique indexes[41], generating library L5387.

For an additional bone powder aliquot from the petrous bone (10.0mg) and for five subsamples from the long bone (between 5.9 and 31.2mg of powder), we followed the extraction protocol described in Rohland et al., 2018[42] on an automated handling platform and using binding buffer "D", without any pre-treatment. We then used an automated version of the single-stranded DNA library preparation protocol[43], generating libraries A25715 from the petrous bone and A29235-A29237 and A29025-A29026 from the long bone fragment. The automated single-stranded DNA library protocol was also used to generate a second library from the extract made from the petrous bone aliquot that had undergone a sodium phosphate wash, yielding library A24767. Three types of data were generated from these libraries: shallow shotgun screening data, mitochondrial DNA capture data and nuclear DNA capture data, which we detail in the next sections.

**Table SI3.1:** Summary of the specimens, subsamples and the downstream laboratory products obtained from the Teshik-Tash 1 individual.

| **Specimen ID** | **Other specimen ID** | **Specimen description** | **Sample weight (mg)** | **Extraction protocol** | **Subsample ID** | **Lysate ID** | **Extract ID** | **Library ID** |
|---|---|---|---|---|---|---|---|---|
| SP.A.1047 | SP999 | Teshik-Tash 123 1/30 long bone fragment | 10.3 | regular | Sub.A.51 | Lys.A.9423 | E.B.7837 | A29235 |
| SP.A.1047 | SP999 | Teshik-Tash 123 1/30 long bone fragment | 11.5 | regular | Sub.A.52 | Lys.A.9424 | E.B.7838 | A29236 |
| SP.A.1047 | SP999 | Teshik-Tash 123 1/30 long bone fragment | 5.9 | regular | Sub.A.53 | Lys.A.9425 | E.B.7839 | A29237 |
| SP.A.3029 | SP2977 | Teshik-Tash 123 1/30 long bone fragment proximal side | 31.2 | regular | Sub.A.393 | Lys.A.9303 | E.B.7731 | A29025 |
| SP.A.3030 | SP2978 | Teshik-Tash 123 1/30 long bone fragment distal side | 12.5 | regular | Sub.A.394 | Lys.A.9304 | E.B.7732 | A29026 |
| SP.A.3640 | SP3589 | Petrous bone | 10.7 | phosphate wash | NA | NA | E.A.3049 | L5387<br>A24767 |
| SP.A.3640 | SP3589 | Petrous bone | 10 | regular | Sub.A.863 | Lys.A.7568 | E.B.5991 | A25715 |

#### Screening

We produced shallow shotgun sequencing data (depth of 3-5 million reads) from all libraries after their amplification by using a 75 bp paired-end read configuration at the Max Planck Institute for Evolutionary Anthropology (MPI-EVA) in Leipzig, Germany, on HiSeq4000 and MiSeq platforms. We demultiplexed the resulting sequences based on perfect matching of the expected index combinations, and aligned the sequences to hg19 human reference genome using BWA (version 0.5.10-evan.9-1-g44db244, https://github.com/mpieva/network-aware-bwa) with the ancient DNA parameters ("-n 0.01 –o 2 –l 16500")[10]. We removed the PCR duplicates using bam-rmdup (https://github.com/mpieva/biohazard-tools/), and filtered the sequences for mapping quality of 25 and minimum length cutoff of 30 base pairs. We calculated the summary statistics reported in **Table SI3.2** using an in-house perl script, and the present-day human contamination using AuthentiCT[17]. We note that this method can not estimate contamination for a subset of sequences with terminal deaminations, which is why we use *admixslug* in a later section. Furthermore, AuthentiCT often requires 10,000 mapped sequences for reliable inference[17], which most of our libraries do not have (**Table SI3.2**). We investigated contamination also using the linear combination method[11,15], which relies on the frequency of derived alleles in present-day humans in a given ascertainment. This method can estimate the present-day human contamination in the deaminated sequences, and has been used frequently for low-coverage Neandertal genomes[11,15,44].

**Table SI3.2:** Summary statistics obtained from screening data generated by shallow shotgun sequencing, per library. Reported present-day human DNA contamination values were estimated by AuthentiCT[17]. The values in the brackets for contamination estimates represent the two standard errors of the mean.

| **Library ID** | **Number of sequenced fragments** | **Number of mapped fragments ≥30bp, MQ≥25** | **% of mapped fragments ≥30bp, MQ≥25** | **Number of unique fragments with ≥30bp, MQ≥25** | **Number of deaminated sequences** | **Present-day human DNA contamination %** |
|---|---|---|---|---|---|---|
| A29235 | 1,074,489 | 2,531 | 3.348 | 2,512 | 478 | 42.0 (37.2, 46.9) |
| A29236 | 735,827 | 5,155 | 7.424 | 5,136 | 730 | 50.6 (46.8, 54.4) |
| A29237 | 801,357 | 2,163 | 3.683 | 2,156 | 319 | 53.4 (47.9, 58.9) |
| A29025 | 2,290,959 | 1,536 | 0.155 | 1,514 | 156 | 61.8 (55.1, 68.5) |
| A29026 | 3,149,792 | 2,301 | 0.137 | 2,274 | 360 | 45.4 (39.8, 51.0) |
| L5387 | 7,039,580 | 46,361 | 1.204 | 46,227 | 2,220 | 48.0 (42.5, 53.5) |
| A24767 | 368,439 | 85,987 | 37.294 | 5,726 | 248 | 44.7 (36.5, 53.0) |
| A25715 | 2,627,729 | 14,457 | 1.51 | 14,392 | 629 | 54.4 (49.4, 59.5) |

#### Substitution patterns

Terminal ancient DNA damage is a measure of endogenous DNA preserved in the sequence libraries[45]. Higher rates of C-to-T substitutions indicate endogenous DNA is well preserved and present-day human contamination is low. A second measure, the conditional substitution rate, measures how much damage is found at one end, when the other end is conditioned to be carrying damage[13]. If the standard substitution rate is low, but conditional substitution rate is high, this can indicate that the library has high present-day human DNA contamination. In agreement with the contamination estimates from AuthentiCT, conditional substitution rates for all libraries showed drastic increase compared to the standard substitution rates (**Table SI3.3**, for complete table see **ST.1**). While this shows that the libraries are contaminated, it also indicates that a small portion of them are ancient, originating from the Teshik-Tash 1 child. Encouraged by this, we produced SNP enrichment data both from the mitochondrial and nuclear genome of Teshik-Tash 1.

**Table SI3.3:** Ancient DNA damage measured as deamination rates at the 5' and 3' ends of the sequences, with 95% CI, per library for screening data.

| Library ID | 5' C-to-T substitution rate (95% CI) | 3' C-to-T substitution rate (95% CI) | 5' C-to-T conditional substitution rate (95% CI) | 3' C-to-T conditional substitution rate (95% CI) |
|---|---|---|---|---|
| A29235 | 30.7 (26.7-35.0) | 31.3 (27.1-35.7) | 63.3 (43.9-80.1) | 57.6 (39.2-74.5) |
| A29236 | 21.3 (18.8-24.1) | 21.4 (18.9-24.1) | 40.5 (24.8-57.9) | 35.7 (21.6-52.0) |
| A29237 | 23.9 (19.9-28.1) | 21 (17.0-25.3) | 25 (5.5-57.2) | 20 (4.3-48.1) |
| A29025 | 19.3 (15.0-24.2) | 14.2 (10.3-18.9) | 40 (5.3-85.3) | 20 (2.5-55.6) |
| A29026 | 25.6 (21.5-30.0) | 29.2 (24.8-33.9) | 48 (27.8-68.7) | 50 (29.1-70.9) |
| L5387 | 6.3 (5.9-6.8) | 7.5 (6.9-8.1) | 24.2 (17.2-32.5) | 29.1 (20.8-38.5) |
| A24767 | 4.5 (3.5-5.7) | 4.6 (3.6-5.8) | 12.5 (1.6-38.3) | 10.5 (1.3-33.1) |
| A25715 | 7.2 (6.2-8.2) | 5.3 (4.5-6.1) | 43.3 (25.5-62.6) | 31.7 (18.1-48.1) |

#### Mitochondrial Capture

Data production and processing

We enriched seven libraries (not including the library prepared manually in 2015) from the two skeletal elements for human mitochondrial DNA (mtDNA) following protocols optimized for ancient DNA[46]. We used two rounds of in-solution hybridisation capture and a probe-set that tiled the revised Cambridge Reference Sequence (rCRS,NC_01290)[47]. We sequenced the capture libraries at the Core Unit facilities of the Max Planck Institute for Evolutionary Anthropology in Leipzig, on MiSeq and HiSeq platforms, on a total of four runs. We merged and trimmed the raw reads using leeHom[48], and mapped the resulting sequences to rCRS reference using BWA version 0.5.10[49] with ancient DNA parameters[10], as previously described in Bossoms Mesa et al.[44]. We demultiplex the sequences using unique 8-base pair index combinations, and removed PCR duplicates using *bam-rmdup* (https://github.com/mpieva/biohazard-tools). To filter the resulting sequences, we used a minimum mapping quality of 25 and minimum read length of 30 basepairs.

The resulting summary statistic can be found in **ST.2**. Out of the seven enriched libraries, only five had a minimum of 10% terminal deamination, our threshold for determining ancient DNA preservation. These five libraries had an average depth of coverage between 14.1-fold and 29.9-fold when considering all sequences, and between 2.2-fold and 4.2-fold when considering only sequences with deamination in the three terminal bases. We explored whether we could use all sequences versus only deaminated sequences by calculating the present-day human contamination in two ways: through lineage-diagnostic positions[13] and with AuthentiCT[17]. In both cases, contamination estimates exceeded 30% for all libraries. Thus, only deaminated sequences were used in downstream analyses.

Even after filtering for only deaminated sequences, we noted that one of the libraries (A29025) still had noticeable more support for present-day human alleles than the rest of the libraries, based on the mentioned diagnostic positions analysis (23% support; while all others were <15%). To assess the potential impact of contamination, we went forward with two different files for Teshik-Tash 1: one with the five libraries with evidence of ancient DNA, and one with A29025 excluded (**ST.2**).

We also explored how different parameters could affect the resulting consensus sequence for the mitochondrial genome of Teshik-Tash 1. More specifically, we tried requiring a minimum coverage of 3-fold and 66% consensus support, as well as a minimum coverage of 4-fold and 75% consensus support, resulting in four total consensus sequences. We defined the coordinates of this consensus sequence with rCRS, the same reference genome that had been used for mapping[50].

Results

None of the four tested consensus, regardless of the number of merged libraries or consensus criteria, showed any pairwise differences (calculated using MEGA version 10.1.7[51]). The number of unresolved bases in the coding region of the mitochondrial genome (defined following Fu et al., 2013[52]) of the consensus ranged from 879 (five libraries, minimum 3-fold coverage and 66% consensus support) to 1,761 (four libraries, minimum 4-fold coverage and 75% consensus support), and in the majority of the cases it was due to insufficient coverage. We present results for the merging of 4 libraries with a minimum 4-fold coverage and 75% consensus support here, but the other three consensus sequences yielded comparable results (not shown).

Firstly, we aligned these consensus sequences with a set of other reference mitochondrial genomes: 33 from Neandertals, one from a first-generation Neandertal-Denisovan offspring, four from Denisovans, one from the archaic hominin from Sima de los Huesos, 53 from present-day humans and one chimpanzee, using *mafft* with 1,000 iterations (version 7.453[53]). The phylogenetic relationship between these genomes was then explored by building a Maximum Likelihood (ML) tree using 100 bootstrap, once more relying on MEGA[51] . This revealed that the Teshik-Tash 1 genome was most closely related to Scladina I-4A, a ~170 – 120 kya juvenile Neandertal mandible from Belgium[9,54].

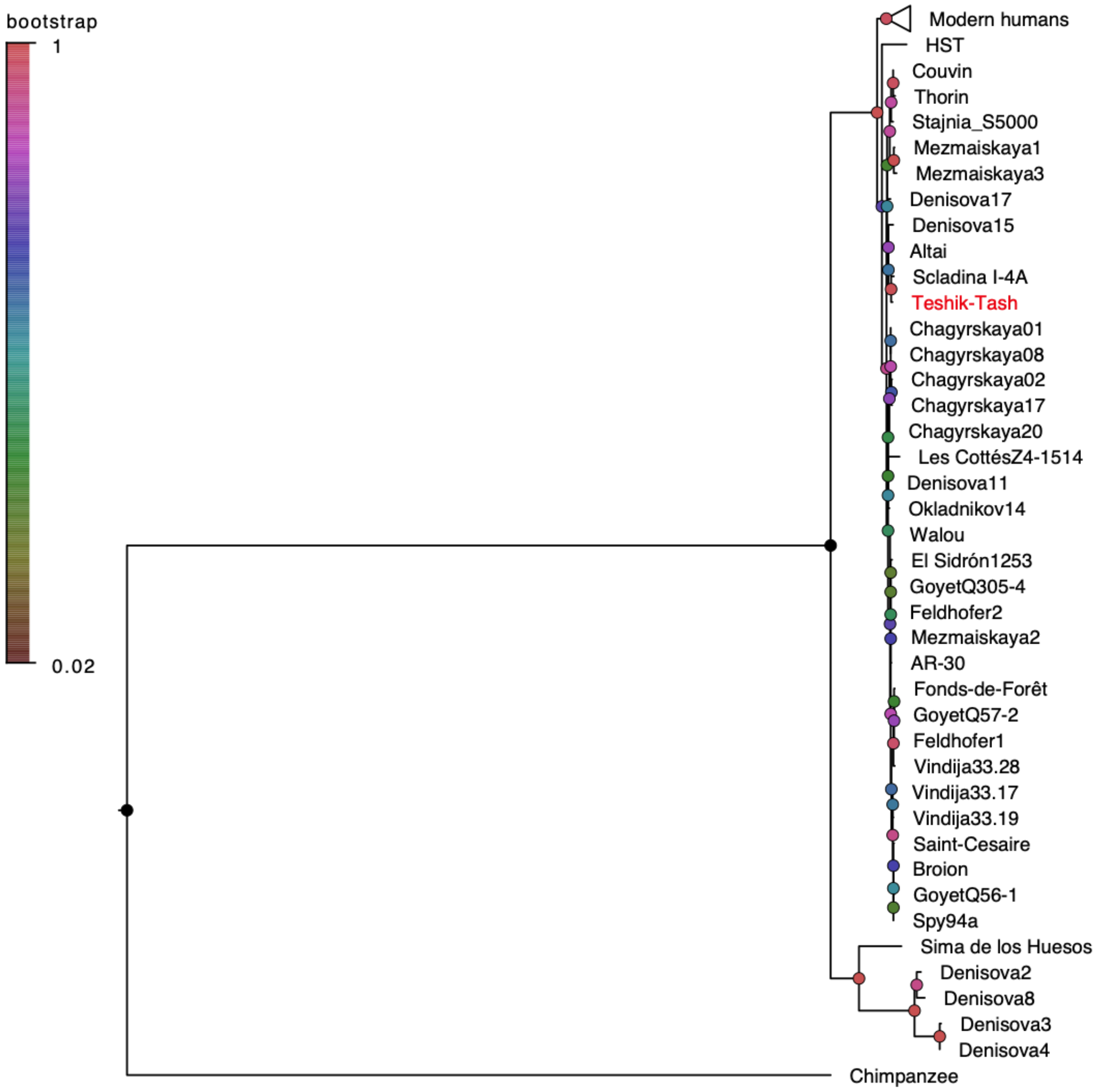


**Figure S3.3:** Maximum Likelihood tree of the described mitochondrial genomes, including the newly generated consensus for Teshik-Tash 1 (in red). The colours in the nodes indicate the bootstrap support for that branching (100 replicates). A total of 13,769 positions were used to calculate the tree using MEGA.

Secondly, we used Kallisto[55] to further explore the phylogenetic affinities of Teshik-Tash 1 to other archaic mtDNA genomes. Kallisto has been successfully applied for ancient DNA research both in sediments[56] and in skeletal remains[44]. This method is more robust to contamination because its "pseudo-aligner" matches the sample sequences (divided in k-mers of a specified size, which in our case was 21 bases) to a set of unique genomic references, thus separating contamination from any potential endogenous signals. This allowed us to use the complete data, and not only sequences filtered for deamination. The results are consistent with the ones described above: the closest match for the Teshik-Tash 1 mtDNA is that of Scladina I-4A (**Figures S3.3 and S3.4**).

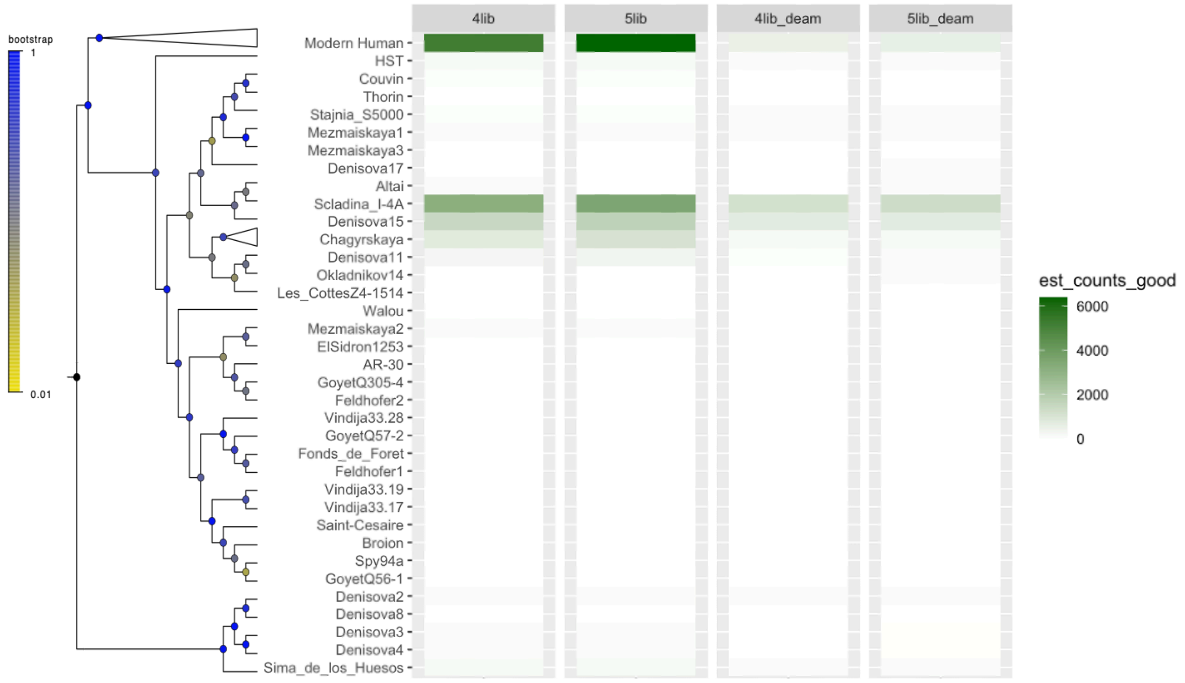


**Figure S3.4:** Kallisto mitochondrial DNA abundances for different Teshik-Tash 1 data (as estimated counts), either using all libraries with aDNA (i.e. 5 libraries or "5lib") or omitting the one with more contamination (i.e. 4 libraries or "4lib"), with and without filtering for deamination; relative to a reference of hominin mitochondrial genomes. On the left, the maximum likelihood tree of the reference dataset.

Unfortunately, because the Teshik-Tash 1 ancient DNA data was low-coverage, we were not able to reconstruct a complete consensus sequence for this Neandertal. The high number of unresolved bases (>1,500 "N" bases, regardless of the consensus and merging criteria selected) thus precluded us from performing molecular dating with methods such as BEAST2[57].

#### Nuclear Capture

Even though all the screened libraries were heavily contaminated with present-day human DNA, they also contained low levels of endogenous DNA from the Teshik-Tash 1 individual. Because of this, we enriched all libraries using an array designed specifically to capture nuclear variation in archaic populations[11] to obtain as much endogenous Neandertal DNA as possible. The capture array we used was introduced in Skov et al., 2022[11], and consisted of 643,472 biallelic transversions across the autosomes with the ascertainment described in SI5 of Skov et al., 2022[11]:

1- SNPs with derived allele frequency > 10% in African genomes from the 1000 genomes phase 3 dataset.

2- SNPs uniformly sampled from 2% frequency bins.

3- SNPs that vary among four high-coverage archaic human genomes that were available at the time of the array design (Altai (D5), Vindija (Vi33.19), Denisova (D3), Chagyrskaya (Chagy8)).
4- Fixed differences between these four archaic human genomes and the African genomes.
5- Sites that are fixed derived in the archaic humans, while not fixed derived in the African genomes.

Our ascertainment for the simulations was very similar, using all ascertainments listed above except SNPs uniformly sampled from 2% frequency bins.

Sequencing of the capture libraries was done at the Core Unit facilities as for the other types of data produced for this study, on HiSeq and MiSeq platforms. Demultiplexing and filtering were performed exactly as described for the shallow shotgun data obtained for screening. To minimize reference bias towards modern human alleles and to keep our data comparable with the previously published low-coverage capture data from Neandertals[11,14,15], we followed the alignment strategy introduced in Peyrégne et al., 2019[15]. Specifically, we aligned our sequences to both the hg19 human reference genome and to a modified reference genome that carries an alternative archaic allele for each of the variants on the array. We merged both alignments per library, and kept all sequences with a mapping quality of 25 or higher, and a length of 30 bp or longer (**Table SI3.4)**.

**Table SI3.4:** Summary statistics from the data obtained from nuclear DNA capture of each library. For the complete table, see **ST.3.**

| Library ID | Capture Library ID | Number of sequenced fragments | Number of mapped fragments ≥30bp, MQ≥25 | Number of unique fragments with ≥30bp, MQ≥25 | Number of deaminated sequences |
|---|---|---|---|---|---|
| A29235 | Cap.G.6907 | 1,802,181 | 680,882 | 24,787 | 4,585 |
| A29236 | Cap.G.6908 | 3,864,996 | 2,431,100 | 112,512 | 19,944 |
| A29237 | Cap.G.6909 | 2,679,565 | 1,349,929 | 43,439 | 6,780 |
| A29025 | Cap.G.6904 | 3,022,054 | 1,261,756 | 41,873 | 6,572 |
| A29026 | Cap.G.6905 | 2,403,057 | 1,005,565 | 34,954 | 6,172 |
| L5387 | Cap.G.6027 | 1,775,358 | 961,888 | 87,464 | 21,537 |
| A24767 | Cap.G.6683 | 5,593,540 | 3,850,085 | 69,412 | 10,019 |
| A25715 | Cap.G.6669 | 9,764,237 | 6,957,378 | 236,759 | 39,428 |

#### Substitution patterns

Similar to what we observed for the screening data, conditional substitution rates were increased compared to standard substitution rates for the capture data produced from all libraries (**Table SI3.5**, **ST.3**). This indicated that the data we produced contained high present-day human DNA contamination.

**Table SI3.5:** Ancient DNA damage measured as deamination rates at the 5' and 3' ends of the sequences, with 95% CI, per library for nuclear capture data.

| Library ID | Capture Library ID | 5' C-to-T substitution rate (95% CI) | 3' C-to-T substitution rate (95% CI) | 5' C-to-T conditional substitution rate (95% CI) | 3' C-to-T conditional substitution rate (95% CI) |
|---|---|---|---|---|---|
| A29235 | Cap.G.6907 | 22.3 (21.2-23.4) | 21.6 (20.5-22.7) | 44.3 (38.7-50.1) | 45.9 (40.2-51.8) |
| A29236 | Cap.G.6908 | 10.3 (9.9-10.7) | 8.7 (8.3-9.0) | 45.6 (41.2-50.0) | 41.1 (37.0-45.2) |
| A29237 | Cap.G.6909 | 10.1 (9.5-10.7) | 9.4 (8.9-10.1) | 42.9 (36.4-49.7) | 43.9 (37.2-50.7) |
| A29025 | Cap.G.6904 | 10.2 (9.6-10.8) | 8.6 (8.0-9.2) | 42.4 (36.0-49.1) | 41 (34.7-47.5) |
| A29026 | Cap.G.6905 | 17.4 (16.6-18.3) | 15.6 (14.8-16.4) | 43.8 (38.5-49.2) | 46.1 (40.6-51.6) |
| L5387 | Cap.G.6027 | 5 (4.7-5.3) | 5.4 (5.1-5.8) | 20.2 (15.2-26.0) | 21.4 (16.1-27.5) |
| A24767 | Cap.G.6027 | 5.1 (4.8-5.5) | 3.9 (3.6-4.3) | 15 (9.8-21.7) | 12.8 (8.3-18.6) |
| A25715 | Cap.G.6669 | 4 (3.9-4.2) | 3.3 (3.2-3.5) | 22.5 (18.6-26.7) | 20.1 (16.6-23.9) |

#### Present-day human DNA contamination

We quantified the levels of contamination introduced by humans living today using three methods, including *admixslug*. Here we present results from two of these, AuthentiCT[17] and linear combination method[13], and we will later compare these with the estimates from *admixslug* (see section **SI3.4**). When the deaminated sequences have minimal or no contamination, such as for libraries A29026 and A29235, AuthentiCT and linear combination estimates of contamination in all sequences are very similar. However, when there are high levels of contamination in the deaminated sequences, such as for libraries L5387 and A24767, the AuthentiCT estimates of contamination in all sequences is much lower than those obtained via linear combination using all sequences (**Table SI3.6**).

**Table SI3.6:** The percentage of present-day human contamination estimates for each library, using two different methods. One of the methods, AuthentiCT requires all sequences and can not be used to estimate contamination only for the deaminated sequences. The values in brackets stand for the 95% CI on each estimate, more specifically for AuthentiCT it is two standard errors of the mean.

| Library ID | AuthentiCT (all sequences) | Linear combination (all sequences) | Linear combination (only deaminated sequences) |
|---|---|---|---|
| A29235 | 60.7 (59.2 - 62.3) | 56.1 (51.3 - 61.0) | 1.1 (0 - 6.8) |
| A29236 | 73.6 (72.6 - 74.6) | 76.5 (74.1 - 79.0) | 10.7 (6.2 - 16.7) |
| A29237 | 80.1 (79.1 - 81.1) | 77.5 (73.6 - 81.4) | 9.7 (3.2 - 20.4) |
| A29025 | 67.4 (65.4 - 69.4) | 75.7 (71.8 - 79.7) | 14.3 (6.3 - 26.2) |
| A29026 | 66.1 (64.6 - 67.5) | 59.9 (55.7 - 64.1) | 5.2 (1.0 - 12.5) |
| L5387 | 53.3 (49.9 - 56.6) | 86.3 (83.5 - .89.2) | 51.8 (40.1 - 64.9) |
| A24767 | 47.5 (42.7 - 52.2) | 87.3 (84.1 - 90.5) | 44.9 (32.1 - 59.9) |
| A25715 | 64.4 (60.8 - 68.1) | 85.8 (84.1 - 87.5) | 39.1 (31.3 - 48.0) |

#### Merged Data

Next, we merged all sequences from all eight captured libraries to obtain the final Teshik-Tash 1 genome which we use in the downstream analyses. We estimated present-day human DNA contamination in the merged set to be 69.6% (68.2%-71.0%, 95% CI), using AuthentiCT[17]. The linear combination method provided a higher estimate for contamination in all sequences (80.7%: 95% CI: 79.6% - 81.8%), as well as detecting high levels of present-day human DNA contamination in the deaminated sequences (22.8%; 95% CI: 19.7% - 26.3%). The average depth of coverage was 0.96x on the target positions, and the breadth of coverage was 396,773 sites. Even though we observed elevated C-to-T substitution rates compared to other types of substitutions, indicating the presence of ancient DNA, these rates were low (~6%), in line with the high estimates of present-day human DNA contamination (**Figure SI3.5**). Additionally, we filtered this set for SNPs that are at least 52 bp apart, which is the length of the probes in the capture array used in this study[11]. This filtering aims to prevent reference bias caused by overlapping flanking parts of probes, as observed in Skov et al., 2022[11]. This removed a total of 42,618 SNPs from the Teshik-Tash 1 genome and left us with 354,155 SNPs (average coverage: 0.84x). For comparative analyses using ADMIXTOOLS and the genotype-mode of *admixslug*, we also generated a final filtered set of Teshik-Tash 1 sequences with C-to-T substitutions at three terminal positions at either end, which we refer to as deaminated sequences here after. The subset of deaminated sequences contained 29,353 SNPs (average depth of coverage 0.044x).

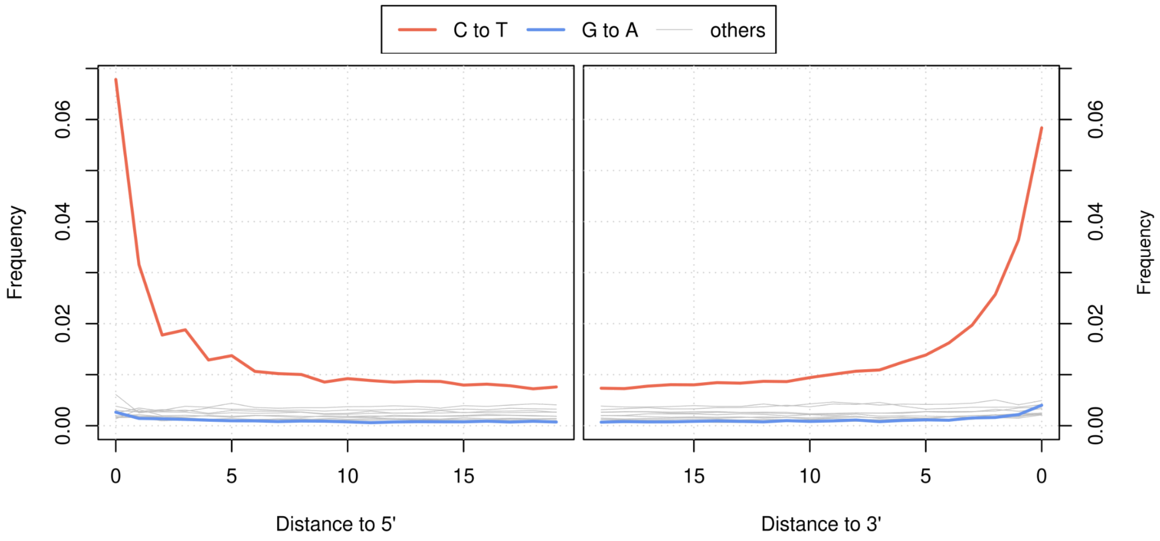


**Figure SI3.5:** Substitution patterns for the Teshik-Tash 1 genome after merging the data from eight libraries together. C-to-T and G-to-A substitutions are plotted in red and blue, respectively, while other substitutions are plotted in gray.

#### SI3.2 - Genetic sex of Teshik-Tash 1

Teshik-Tash 1 is a child that died before their morphological characters informative of sexual dimorphism could be formed. Due to this, to our knowledge, it had not been possible to determine if the child was a male or a female. To investigate this, we calculated the ratio of the sequences mapped to the X chromosome versus the autosomes. Because there are two copies of the X chromosome in females and only one in males, we would expect this ratio to be around one if the Teshik-Tash 1 child was a female, and 0.5 if male. To resolve this, we separately analysed sequences obtained from both SNP capture and shallow shotgun sequencing, and found concordant results.

##### Capture data

The capture array used in this and previous studies (Skov et al. 2022[11], Bossoms Mesa et al. 2026[44]) targets positions on the autosomes and the X chromosome. Previously, 11,886 sites on the X chromosome and 402,801 sites on the autosomes were used for genetic sex determination of Neandertals[11]. Following the same approach, we estimated the X-to-autosome ratio for Teshik-Tash 1 genome, using only the deaminated sequences. This resulted in a proportion of 0.024 (290 out of 11,886 target sites covered) on the X chromosome and of 0.048 on the autosomes (19,410 out of 402,801 target positions covered) **ST.4**. Therefore, the X-to-autosomes ratio was 0.506, suggesting that Teshik-Tash 1 was a male. We estimated the binomial confidence interval on this ratio as 0.450 - 0.568.

##### Shallow-shotgun data obtained for screening

The capture array we use for the enrichment of the SNPs in nuclear genomes does not target positions on the Y chromosome. Due to this, we repeated our analyses with the sequences obtained through shallow shotgun sequencing performed during the screening of libraries from Teshik-Tash 1. To reduce the present-day human DNA contamination in the libraries (**Table SI3.6**), we restricted this analysis to the sequences with terminal C-to-T substitutions, which resulted in only 305 positions covered on the Y-chromosome, and 6,207 positions on the larger X chromosome. The chromosome 7, which is similar in size to the X chromosome, had 12,127 positions covered. Overall, the X chromosome to autosome ratio was ~0.54 ($4.11x10^{-5}/7.63x10^{-5}$) indicating that Teshik-Tash 1 was a boy (**ST.5**). We investigated the 305 positions covered on the Y-chrosomosome to see if they provide any support for one of the known Neandertal Y-chromsome lineages[11,58], however none of the sites covered was among the informative positions.

#### SI3.3 - Comparative data set for nuclear DNA analyses

For comparison, we analysed low-coverage genome-wide data from two European Neandertals lived ~100ky ago in Hohlenstein-Stadel, and Scladina[15], two Neandertals from Gibraltar from Forbes

Quarry and Devil's Tower[14], one ~65ky old Neandertal from Mezmaiskaya[6], and two late Neandertals from Spy and Mezmaiskaya[9]. We subsetted these published low-coverage genomes to the positions targeted by the capture array we use for Teshik-Tash 1. While this reduces the number of sites covered for analyses for the published genomes, the final coverage is still within the range *admixslug* can perform, except maybe for the Devil's Tower child. For this individual, even though the coverage was very low, we wanted to test how *admixslug* performed.

#### SI3.4 - *Admixslug* results

We ran *admixslug* on the Teshik-Tash 1 data we generated, as well as the previously published low coverage Neandertal genomes described above, with parameters:

```
admixslug --in input_file \
--states VIN CHA ALT DEN \
--ref reference_file \
--ancestral PAN --cont-id EUR \
-o output_name \
--ptol 0.001 --ll-tol 0.01 --max-iter 100 \
 --filter-ancestral --len-bin-size 2000 \
--jk-resamples 500 --output-jk-sfs --output-fstats
```

First, we compare the previously published present-day human DNA contamination estimates for the four genomes in our comparative set, with the new estimates obtained from *admixslug*. We also present contamination estimates in the sequences with deamination.

##### Present-day human DNA contamination

*admixslug* estimates present-day human DNA contamination in the target genome, for several different categories, such as read-groups (sequence libraries), deaminated or non-deaminated sequences, different read length groups, etc. For the values reported here, we combined the estimates per library, and for all sequences or deaminated sequences for simplicity. We find that the eight libraries from Teshik-Tash 1 carried between 58% and 95% contamination, when all sequences are used (**Figure S3.6**). These values were higher than those obtained through AuthentiCT, and aligned better with the estimates of the linear combination method (**Table SI3.6**). Furthermore, we observe for some of the libraries, significant levels of contamination in the deaminated sequences, in line with the results of the linear combination method. Often it is not possible to detect this type of contamination, with sequences carrying terminal deamination considered endogenous. This assumption biases downstream analyses in the presence of contamination in deaminated sequences.

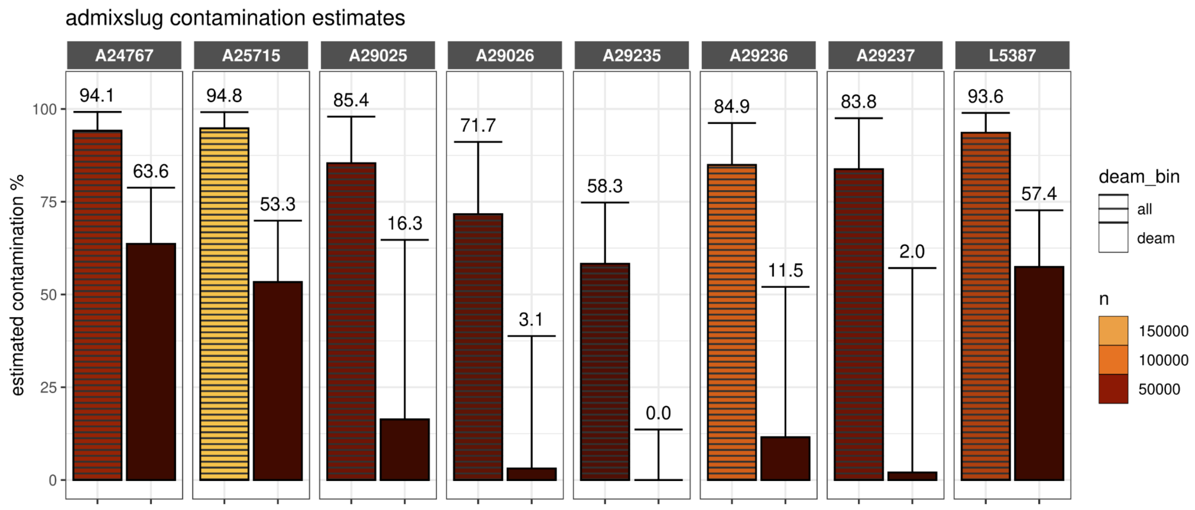


**Figure S3.6:**Present-day human DNA contamination estimated by *admixslug*, for all sequences and those with terminal deamination (striped and non-striped, respectively). The values on the bars indicate the mean values of the estimates, while error bars represent the 95% CI. The number of sequences used in each estimate is colour-coded.

Overall, for the genome of Teshik-Tash 1 after merging data from all libraries, *admixslug* estimated present-day human DNA contamination to be 89.0% (78.6%-97.2%, 95% CI) overall and 27.2% (16.4%-56.2%, 95% CI) for deaminated sequences. We also investigated how the levels of present-day human contamination changes for different read-lengths. The parameter `--len-bin-size` controls the number of sequences clustering in each bin. We set this value to 2,000 in order to observe if there is a difference in contamination in deaminated sequences as they get longer. For 6 out of 8 libraries, the number of deaminated sequences were too low to cluster in two groups (less than 2,000 sequences). Two libraries with more deaminated sequences (A25715 and A29236) resulted in two groups, in which we observe higher contamination in longer sequences (**Figure S3.7**). Gene-flow from an ancient modern human into the ancestors of Teshik-Tash 1 individual could have resulted in modern-human sequences with deamination, which could be interpreted as present-day human contamination with ancient DNA signal. However, observing an increase in contamination with increasing sequence length also in the deaminated sequences indicates that this is likely not the case.

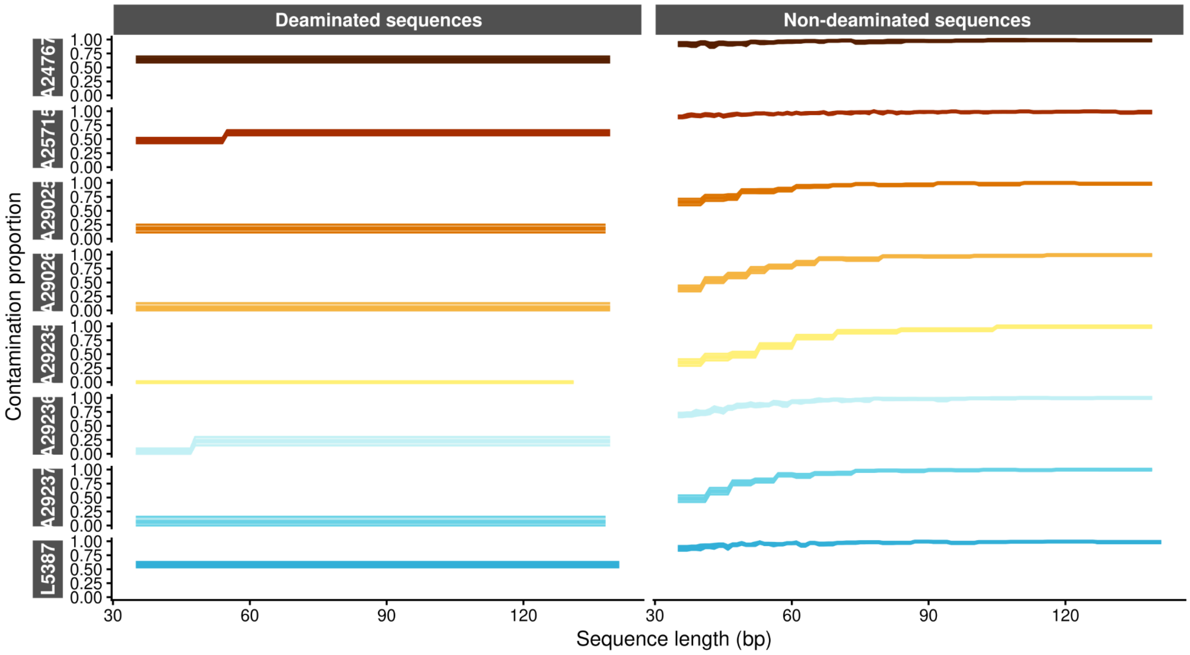


**Figure S3.7:** Present-day human DNA contamination estimated by *admixslug* for different length bins, for sequences with and without terminal C-to-T substitutions (deaminated and non-deaminated, respectively). The y-axis shows the contamination proportion, and the x-axis represents sequence lengths in each library.

**Table S3.7:** The present-day human contamination estimates and average depth of coverage in all sequences and deaminated sequences of all low-coverage Neandertal genomes included in this study. (plotted in the main **Figure 1B**). The contamination estimates are estimated by *admixslug*.

| Specimen | Average depth of coverage | Sequences | Present day human contamination mean (%) | Present day human contamination low (%) | Present day human contamination high (%) |
|---|---|---|---|---|---|
| Mezmaiskaya1 | 1.9-fold | all | 2.6 | 2.3 | 3 |
| Mezmaiskaya2 | 1.7-fold | all | 0.66 | 0.45 | 0.89 |
| Scladina_I-4A | 0.042-fold | all | 68.4 | 65 | 71.9 |
| Spy94a | 1-fold | all | 3.63 | 3.33 | 3.94 |
| Forbes Quarry | 0.075-fold | all | 6.7 | 4 | 9.7 |
| Devil's Tower | 0.003-fold | all | 69.1 | 60 | 78 |
| Hohlenstein-Stadel | 0.09-fold | all | 28.6 | 25 | 32 |
| Teshik-Tash 1 | 0.84-fold | all | 89 | 78.60 | 97.2 |
| Teshik-Tash 1 | 0.044-fold | deaminated | 27.2 | 16.4 | 56.2 |
| Mezmaiskaya1 | 0.394-fold | deaminated | 0.48 | 0.33 | 0.63 |
| Mezmaiskaya2 | 0.56-fold | deaminated | 0.4 | 0.17 | 0.63 |
| Scladina_I-4A | 0.0065-fold | deaminated | 3.57 | 0.35 | 6.99 |
| Spy94a | 0.22-fold | deaminated | 0.3 | 0.06 | 0.5 |
| Forbes Quarry | 0.038-fold | deaminated | 1.6 | 0.2 | 4 |

| | | | | | |
|---|---|---|---|---|---|
| Devil's Tower | 0.0002-fold | deaminated | 0.1 | 6.4 | 0 |
| Hohlenstein-Stadel | 0.028-fold | deaminated | 2.53 | 0.9 | 4.28 |

#### Unbiased *f*-statistics and F(A|B)

##### ***$F_4$-statistics*** to test if Teshik-Tash 1 was closer to Altai or Vindija

Teshik-Tash 1 represents the currently known edge of south-eastern dispersal of the Neandertals (**Figure SI3.2)**. On the mitochondrial level, he clusters together with an early Neandertal from Belgium (Scladina), and with a ~120,000 years old Neandertal from the Altai mountains (D5), rather than with the later Neandertals from Europe often represented by the Vindija Neandertal (Vi33.19) (**Figures S3.3 and S3.4**). Taken together with the uncertainty on the age of the specimen, the genetic background of Teshik-Tash 1 is difficult to predict.

We used *admixslug* to test if Teshik-Tash 1 is genetically more similar to the geographically closer Altai Neandertal, which he also clusters with on the mitochondrial level, or to the Vindija Neandertal from Croatia who lived ~45,000 years ago. We obtained a significantly positive value for the statistics *$f_4$(Vindija, Altai, target, Chimpanzee),* showing that Teshik-Tash 1 is genetically closer to the Vindija Neandertal when compared with the Altai Neandertal. When we repeat the test to ask if he was closer to Vindija or Chagyrskaya, the statistic is again positive, indicating closer affinity to Vindija, however not significantly so (**Figure S3.8**).

For comparison, we included the shotgun genomes detailed in the beginning of SI3.3. Our results for this comparative set aligned with results previously published for these individuals. For the Vindija-like late Neandertals such as Spy 94a and Mezmaiskaya 2, we observed the highest values for the *$f_4$-statistics $f_4$(Vindija, Altai, target, Chimpanzee),* which tests if the target is closer to the Vindija or Altai Neandertal. This was also true to a lower extent for the other Neandertals in our comparative set, except for the Devil’s Tower individual from which data were not enough to obtain meaningful statistics (**Figure S3.8**).

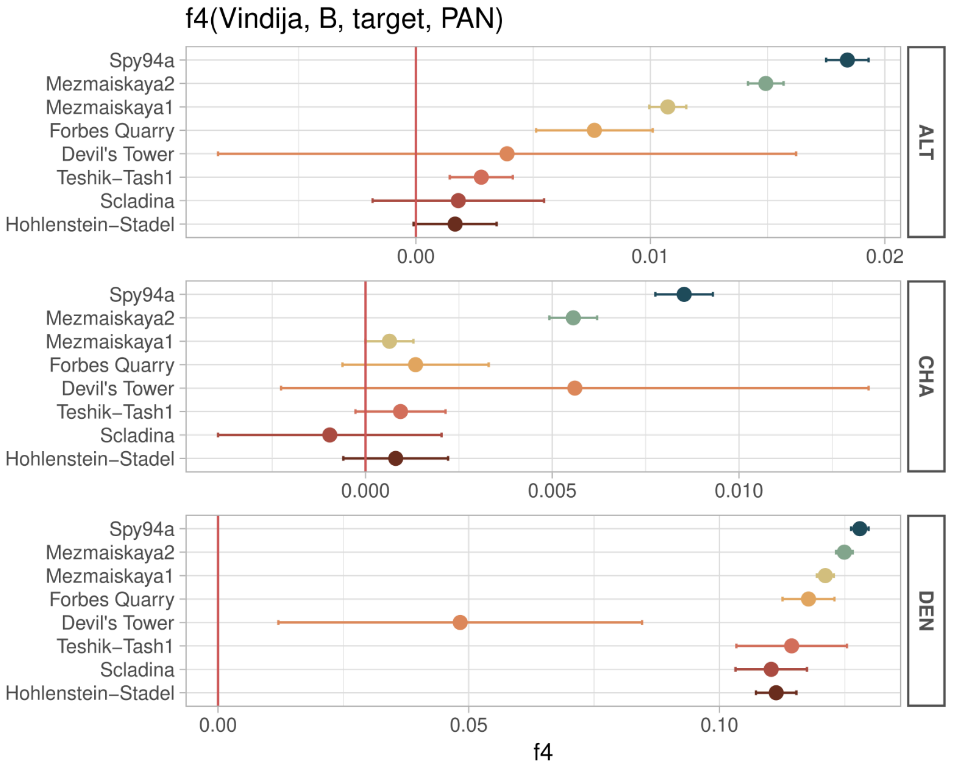


**Figure S3.8:** $f_4$*-statistics* in the form of $f_4$*(Vindija, B, target, Chimpanzee)*, where the high-coverage genome used in position B is denoted on the right y-axis and low-coverage genomes used in the target position on the left y-axis. The x-axis shows the $f_4$ estimates, with error bars corresponding to 95% confidence interval (calibrated as 1.5 standard errors of the mean), obtained from 500 jackknives using a bin size of 2,000.

##### $F_4$*-statistics* to detect Denisovan ancestry

Because Teshik-Tash 1 geographically and temporally overlaps with the inferred range of the Denisovans, he could have recent Denisovan ancestry in his genome. Denisovan ancestry in Neandertal genomes was previously investigated using admixfrog, a tool developed for segment calling from ancient DNA data[59]. Results from previous studies indicate that while traces of Denisovan ancestry can be found in Neandertals older than 100,000 years who lived in the Altai Mountains, this was not the case for later Neandertals in Europe[6,9,44,60]. Unfortunately, due to the ultra-low coverage of the Teshik-Tash 1 genome, it was not possible to call ancestry segments using this method.

We used *admixslug* to calculate $f_4$-statistics to investigate Denisovan ancestry in Teshik-Tash 1 and three Neandertal genomes without Denisovan ancestry (Mezmaiskaya 1[6], Mezmaiskaya 2 and Spy 94a[9]). For comparability to the Teshik-Tash 1 genome, we generated downsampled and contaminated versions of these three published genomes following the same methodology detailed in section SI2.1.

We ran *admixslug* with parameters:

```
admixslug --infile input_file \
--ref reference_file \
-o output_name \
--states VIN CHA DEN --cont-id EUR --ancestral PAN \
--ptol 0.001 --ll-tol 0.01 --max-iter 100 --filter-ancestral \
--jk-resamples 100 --output-jk-sfs --output-fstats
```

Note that we removed ALT (Altai Neandertal, D5) from the states in these runs, because this Neandertal genome carries Denisovan ancestry[7,59] and could potentially bias our results. We calculated two $f_4$-*statistics* that inform us about the existence of Denisovan ancestry in the Teshik-Tash 1 genome: statistics $f_4$*(Denisova 3, B, target, Chimpanzee)* and $f_4$*(target, B, Denisova 3, Chimpanzee)*. B is either the Vindija 33.19 or Chagyrskaya 8 high-coverage Neandertal genomes that do not carry any Denisovan ancestry, and target genome is Teshik-Tash 1 or the test genomes of Mezmaiskaya 1, Mezmaiskaya 2 and Spy 94a.

The first statistics test if the Denisovan or B is closer to the target. If Teshik-Tash 1 carries Denisovan ancestry, we expect to observe a higher value of this statistics when compared to the test genomes that do not carry any Denisovan ancestry (**Figure SI3.9, Panel A**). Estimates of these statistics did not differ for Teshik-Tash 1 and other test genomes, indicating that Teshik-Tash 1 does not have Denisovan ancestry.

The second statistics, $f_4$*(target, B, Denisova 3, Chimpanzee)* tests if the target or the genome in B is closer to Denisova 3. Both Neandertal genomes in position B, Chagyrskaya 8 and Vindija 33.19 are known not to carry any Denisovan ancestry. Our results were centered around 0, indicating equal affinity of the target and B to Denisova 3 (**Figure SI3.9, Panel B**). When taken together, these two statistics point to the absence of Denisovan ancestry in the Teshik-Tash 1 genome, similar to the Mezmaiskaya 1, Mezmaiskaya 2 and Spy 94a genomes.

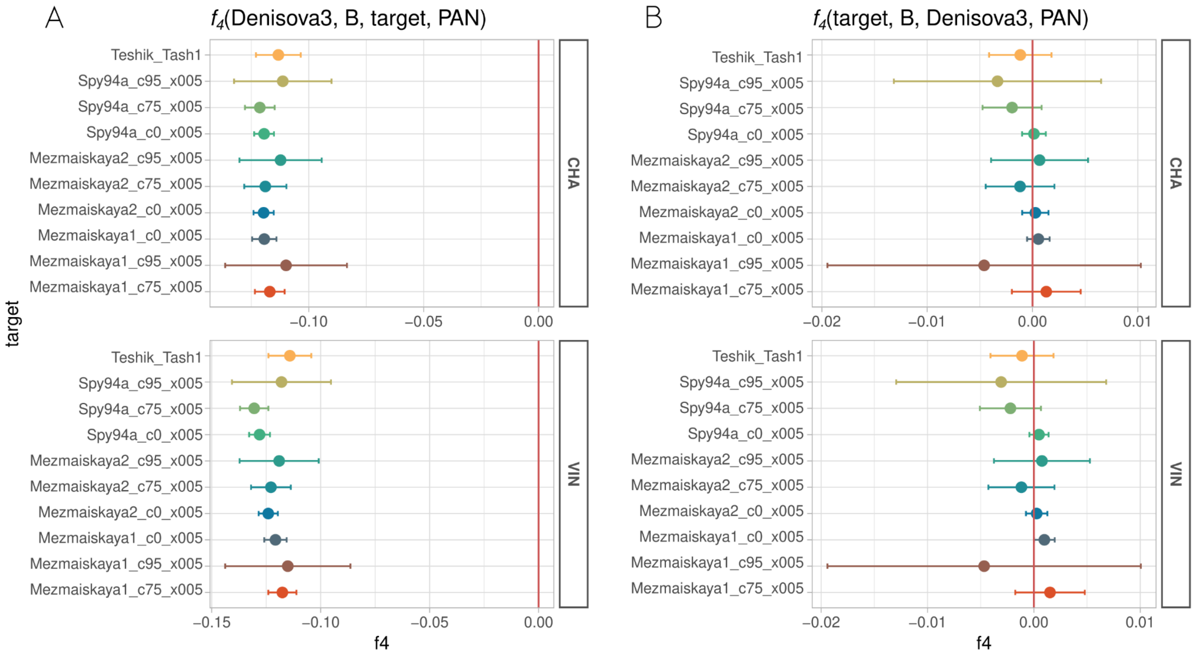


**Figure SI3.9**: *$f_4$-statistics* testing **A.** if Denisova 3 or the high-coverage genome listed on the right y-axis (Vindija or Chagyrskaya) is closer to the target, and **B.** if target or the high-coverage genome listed on the right y-axis is closer to Denisova 3. The error bars represent 95% CI (calibrated as 1.5 se) of the mean estimate.

#### F(A|B) statistics for inferring split order

The output of *admixslug* can be used to calculate F(A|B) statistics, which measures the proportion of heterozygous sites in a diploid genome *B* that share a derived allele with a pseudohaploid genome *A*[7]. Here, genome *B* represents the high-coverage archaic human genome in our reference panel, and genome *A* is the low-coverage genome we are interested in. Because the F(A|B) estimate does not rely on the demography of the population of individual *A*, this method was previously used for estimating split-times from low-coverage Neandertal genomes[15].

Similar to our observations from the $f_4$-statistics, we find that the Devil's Tower genome is too low-coverage to provide meaningful results (**Figure SI3.10**). The rest of the genomes yield comparable estimates of the F(A|B) statistics where genome *B* is the Denisovan (D3) or Altai Neandertal (D5), showing that they all belong to a lineage that split from the two, before they split from each other.

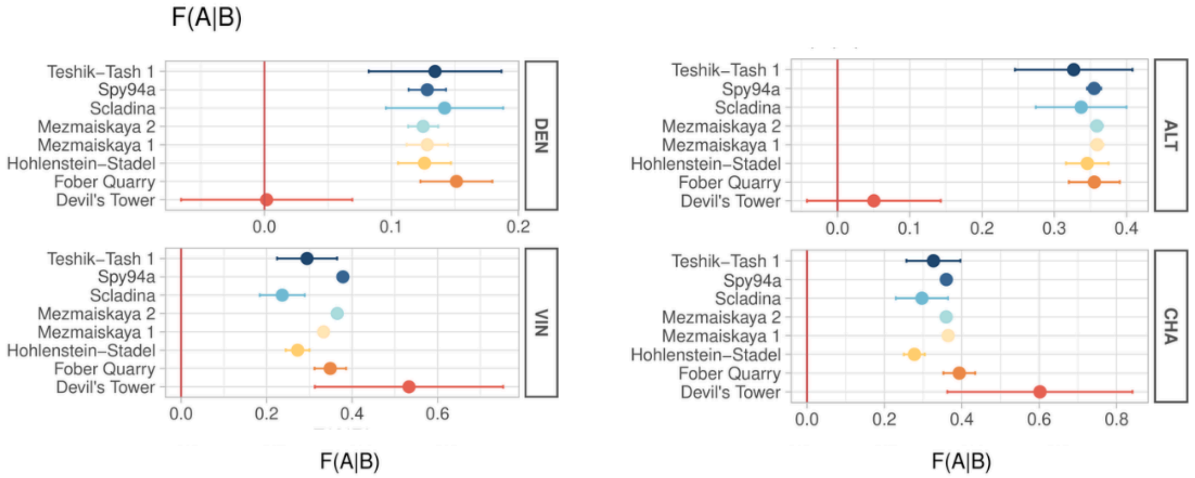


**Figure SI3.10:** F(A|B) statistics where the high coverage archaic human genomes Denisovan (D3), Vindija, Chagyrskaya and Altai (D5) are in position "B", and low-coverage genomes listed on the y-axis are in position "A". The error bars represent the 95% confidence interval (2 standard errors).

When the genome of the Vindija Neandertal is *B*, we see different values of the statistics for the target genomes. As expected, the late Neandertals that split from Vindija latest have the highest F(A|B) values. The succession of the values of this statistics gives us the splitting order on the Vindija lineage, where lower values of F(A|Vindija) correspond to deeper splits, and higher values to more recent splits. We find that Teshik-Tash 1 splits from the Vindija lineage after Scladina and Hohlenstein-Stadel, but before Mezmaiskaya 1, Forbes Quarry and the late Neandertals Spy 94a and Mezmaiskaya 2 (**Table SI3.8**). This places the split time of this individual approximately between 80 ka and 100 ka, based on the previously published split time estimates[8,14,15].

**Table SI3.8:**F(A|B) statistics plotted in **Figure SI3.9**.

| | **Vindija** | | **Chagyrskaya** | | **Altai** | | **Denisova** | |
|---|---|---|---|---|---|---|---|---|
| **Individual** | **Mean F(A\|B)** | **Standard error** | **Mean F(A\|B)** | **Standard error** | **Mean F(A\|B)** | **Standard error** | **Mean F(A\|B)** | **Standard error** |
| Teshik-Tash 1 | 0.295 | 0.0351 | 0.327 | 0.0349 | 0.327 | 0.0407 | 0.134 | 0.0262 |
| Mezmaiskaya 2 | 0.365 | 0.00377 | 0.36 | 0.00384 | 0.359 | 0.00419 | 0.125 | 0.00591 |
| HST | 0.273 | 0.0139 | 0.277 | 0.0135 | 0.346 | 0.0147 | 0.126 | 0.0104 |
| Scladina | 0.237 | 0.0262 | 0.297 | 0.0338 | 0.337 | 0.0314 | 0.142 | 0.0231 |
| Forbes Quarry | 0.349 | 0.0184 | 0.394 | 0.0205 | 0.355 | 0.0176 | 0.151 | 0.0337 |
| Devil's Tower | 0.533 | 0.11 | 0.602 | 0.12 | 0.0503 | 0.0463 | 0.00185 | 0.0337 |
| Spy 94a | 0.379 | 0.0048 | 0.36 | 0.00537 | 0.355 | 0.00486 | 0.128 | 0.00735 |
| Mezmaiskaya 1 | 0.333 | 0.00452 | 0.365 | 0.00418 | 0.359 | 0.00442 | 0.128 | 0.00812 |

#### SI4: Sex determination based on proteomics

In this section, we summarize the findings of another paper that is currently under review, from some of the authors of this manuscript (R.Z. and A.B.) (Buzhilova and Ziganshin, in review).

A permanent mandibular incisor of Teshik-Tash 1 was used as the sample for sex determination using proteomics. A method validated by the authors—chromatography-mass spectrometry—was employed to analyze dental enamel peptides. This method was optimized by acid-etching tooth enamel for 8 minutes, followed by hydrolysis of extracted proteins by chymotrypsin and desalting the peptide products using SDB-RPS StageTips microcolumns. Subsequent analysis of one-third of the desalted sample via liquid chromatography-mass spectrometry (LC-MS) enabled reliable sex determination of fossil remains. The approach proved effective across a broad range of archaeological and biological ages while preserving the structural integrity of the teeth[61].

The biological sex of an individual is determined by detecting amelogenin Y-chromosome specific peptides (≥2) within the identified peptide profile. For conclusive results, a minimum of 30 amelogenin X peptide fragments must also be detected in the sample to ensure accurate sex differentiation.

The quantitative results of unique peptide identification from tooth enamel proteins are summarized in **ST.6**. According to methodological recommendations, the number of unique Y-chromosome specific peptides (more than two) is sufficient to confirm male sex. Additionally, we verified the presence of the amelogenin Y-chromosome specific peptide sequence [SM(+15.99)IRPPYSP]. This peptide was found in all male-derived samples without exception in previous experiments[61]. Y-chromosome specific peptides identified for Teshik-Tash 1 in our analysis (**ST.7**) show that the child was male.
